# Hypermethylation and small RNA expression are associated with increased age in almond (*Prunus dulcis* [Mill.] D.A. Webb) accessions

**DOI:** 10.1101/2021.05.02.442365

**Authors:** Katherine M. D’Amico-Willman, Chad E. Niederhuth, Michael Sovic, Elizabeth S. Anderson, Thomas M. Gradziel, Jonathan Fresnedo Ramírez

## Abstract

- The focus of this study is to profile changes in DNA methylation and small RNA expression occurring with increased age in almond breeding germplasm to identify possible biomarkers of age that can be used to assess the potential of individuals to develop aging-related disorders.
- To profile DNA methylation in almond germplasm, 70 methylomes were generated from almond individuals representing three age cohorts (11, 7, and 2 years old) using an enzymatic methyl-seq approach followed by analysis to call differentially methylated regions (DMRs) within these cohorts. Small RNA (sRNA) expression was profiled in three breeding selections, each from two age cohorts (1 and 6 years old) using sRNA-Seq followed by differential expression analysis.
- Weighted chromosome-level methylation analysis reveals hypermethylation in 11-year old almond breeding selections when compared to 2-year-old selections in the CG and CHH contexts. Seventeen consensus DMRs were identified in all age contrasts. sRNA expression differed significantly between the two age cohorts tested, with significantly decreased expression in sRNAs in the 6-year-old selections compared to the 1-year-old.
- Almond shows a pattern of hypermethylation and decreased sRNA expression with increased age. Identified DMRs and differentially expressed sRNAs could function as putative biomarkers of age following validation in additional age groups.

## Introduction

The study of aging has centered primarily around mammalian systems with a focus on humans (Kirkwood, 2005; Ferrucci *et al*., 2020); however, the aging process has also been shown to impact plants with emphasis placed on long-lived perennials (Munné-Bosch, 2007; Brutovská *et al*., 2013; Thomas, 2013; Woo *et al*., 2018). These impacts can include diminished growth and reduced flower and fruit production, as well as the development of aging-related disorders (Kester & Jones, 1970; Van Dijk, 2009). In perennial plants and other organisms such as humans, causal mechanisms underlying the development of age-related phenotypes include genetic alterations such as somatic mutations or differential chromatin marks as well as changes in expression of small non-coding RNAs (sRNAs) (Jaligot *et al*., 2000; Kato *et al*., 2011; Dubrovina & Kiselev, 2016; Ogneva *et al*., 2016; Xiao *et al*., 2019; Wang *et al*., 2020). In fact, DNA methylation has been proposed as a biomarker of aging in many systems, serving as a biological “clock” that can be used to track aging and predict aging outcomes (Runov *et al*., 2015; Jylhävä *et al*., 2017; Xiao *et al*., 2019).

Profiling genome-wide DNA methylation is one approach to quantify differential epigenetic marks and model alterations associated with advanced age. Identification of specific regions of the genome showing changes in methylation associated with aging provides the opportunity to develop biomarkers to track aging and information on those genes or genic regions that might contribute to the development of age-related phenotypes (Xiao *et al*., 2019). Alterations in DNA methylation patterns associated with aging have been recently reported in *Pinus*, suggesting DNA methylation could serve as a biomarker for tracking age in plants (Gardner *et al*., 2023).

In addition to DNA methylation, sRNAs are known to modulate plant development and have been previously shown to be associated with aging (Chen, 2009; D’Ario *et al*., 2017; You *et al*., 2022). The function of sRNAs, specifically microRNAs, in these processes occurs primarily through regulating target gene expression and via post-transcriptional control (Chen, 2009; D’Ario *et al*., 2017; You *et al*., 2022). Due to their direct association and involvement in regulating plant developmental processes, including the juvenile-to-adult transition, microRNAs represent putative targets for developing biomarkers of age in a clonally propagated species such as almond (Raihan *et al*., 2021; You *et al*., 2022).

Studying alterations like differential DNA methylation and sRNA expression associated with advanced age in perennial plant systems can: (1) provide a means to track aging in these systems and (2) lead to an increased understanding of the development of age-related disorders or degeneration of important physiological processes in plants. This information is valuable to agricultural industries that rely on sustained production of perennial crops, including fruit and nut trees. Almond (*Prunus dulcis* [Mill.] D.A. Webb) is an example of a perennial nut crop that is negatively affected by the aging process through the exhibition of non-infectious bud failure, an aging-associated disorder (Kester & Jones, 1970; Micke, 1996; Kester *et al*., 2004). Additionally, almond trees are primarily produced by clonal propagation for orchard establishment, meaning age and thus susceptibility to age-related impacts is difficult to determine (Ally *et al*., 2010; de Witte & Stöcklin, 2010; Salguero-Gomez, 2018). A method to track aging, particularly in crops like almond produced by clonal propagation or shown to exhibit age-related disorders, would benefit growers, producers, and consumers by helping to protect the supply chain of these valuable commodities.

In this study, we utilize almond breeding germplasm produced in pedigreed crosses as part of the almond breeding program at the University of California, Davis. The individuals used in this study are grown from seed and thus are of known age, making them particularly useful to generate models to track aging in this species where clonal propagation is standard. The goal of this study was to examine DNA methylation patterns in the genome of a productive perennial crop by performing an exhaustive methylome profiling of ∼70 almond individuals from three distinct age cohorts. The hypothesis is that the almond breeding selection cohorts will exhibit, on average, divergent DNA methylation profiles associated with age. Further, we performed small RNA sequencing almond selections from two age cohorts to identify differentially expressed sRNAs associated with aging. Our overall aim is to identify variability in the almond methylome and transcriptome that could enable model development to track aging in this clonally propagated crop and provide targets (i.e., differentially methylated regions, differentially expressed sRNAs) for further investigation into mechanisms influencing age-related phenotypes such as non-infectious bud failure or the juvenile-to-adult transition. This work also serves as a model to explore aging and its impacts on other important perennial crops.

## Materials and Methods

### Plant Material

Almond leaf samples were collected in May 2019 from the canopy of 30 distinct breeding selections planted in 2008, 2012, and 2017, totaling 90 individuals sampled. These selections represent three almond age cohorts aged 11, 7, and 2 years at the time of sampling. Almond breeding germplasm sampled for this study is maintained at the Wolfskill Experimental Orchards (Almond Breeding Program, University of California – Davis, Winters, CA). Leaf samples were collected in the field and immediately stored on ice and then at -20 °C until shipping. Samples were shipped on ice to the Ohio Agricultural Research and Development Center (OARDC; The Ohio State University, Wooster, OH, USA) and immediately stored at -20 °C until sample processing. Almond leaf tissue from the canopy of three breeding selections planted in 2015 and 2020 was collected in May 2021 from the Wolfskill Experimental Orchards. Samples were collected, stored on dry ice, and immediately sent to Novogene for processing and small RNA sequencing.

### DNA Extraction

High-quality DNA was extracted from leaves following a modified version of the protocol outlined in Vilanova et al., 2020. Briefly, samples were ground to a fine powder with a mortar and pestle in liquid nitrogen, and 50 mg of the ground material was added to 1 mL of extraction buffer (2% w/v CTAB; 2% w/v PVP-40; 20 mM EDTA; 100 mM Tris-HCl [pH 8.0]; 1.4 M NaCl), 14 µL beta-mercaptoethanol, and 2 µL RNase (10 mg/mL). The solution was incubated at 65°C for 30 mins and on ice for 5 mins, followed by a phase separation with 700 µL chloroform:isoamyl alcohol (24:1). The aqueous phase (∼800 µL) was recovered, and 480 µL binding buffer (2.5 M NaCl; 20% w/v PEG 8000) was added followed by 720 µL 100% ice-cold ethanol.

A silica matrix buffer was prepared by adding 10 g silicon dioxide to 50 mL ultra-pure water prior to incubation and centrifugation steps. Silica matrix buffer (20 µL) was added to each sample, and samples were gently mixed for 5 mins. Samples were spun for 10 secs, and the supernatant was removed. To resuspend the remaining mucilaginous material (but not the pellet), 500 µL of cold 70% ethanol was used, and the supernatant was removed. Another 500 µL cold 70% ethanol was added to resuspend the silica pellet, the tubes were spun for 5 secs, and the supernatant was removed. The pellet was allowed to dry at room temperature for 5 mins and was resuspended in 100 µL elution buffer (10 mM Trish HCl [pH 8.0]; 1 mM EDTA [pH 8.0]) followed by a 5 min incubation at 65°C. Samples were centrifuged at 14,000 rpm for 10 mins at room temperature, and 90 µL of supernatant was transferred to a new tube. DNA concentration was assessed by fluorometry using a Qubit™ 4 and Qubit™ 1X dsDNA HS Assay Kit (ThermoFisher Scientific, Waltham, MA, USA).

### RNA Extraction

For small RNA sequencing, RNA was isolated from almond leaves using the Quick-RNA Plant Miniprep Kit (Zymo cat# R2024) with minor modifications. Leaf tissue was transferred into 800 µl of DNA/RNA Shield in the BashingBead tubes (instead of 800 µl of RNA Lysis Buffer as written in the protocol) and homogenized using an MP FastPrep-24 with the following settings: 6m/s for 60s, repeat for a total of 2 cycles. After centrifuging to pellet debris, 400 µl of cleared lysate was taken out and mixed thoroughly with 1200 µl (3 volumes) of RNA Lysis Buffer. This mixture was then added to the ZymoSpin IIICG column as described in Step 4 of the protocol, and then the rest of the protocol was followed as written. DNase treatment was performed in the column according to the appendix on page 6.

### Enzymatic Methyl-Seq Library Preparation and Illumina Sequencing

Whole-genome enzymatic methylation sequencing is equivalent to the “gold standard” bisulfite sequencing approach to profile the methylome at the nucleotide level (Feng *et al*., 2020). Utilizing this approach provides information on both genome-wide methylation in each context (CG, CHG, and CHH [H = A, T, or C]) and allows for the identification of differentially methylated regions (DMRs), pinpointing regions of the genome showing dynamic patterns of methylation associated with increased age (Feng *et al*., 2020; Vaisvila *et al*., 2020). Whole genome enzymatic methyl-seq libraries were prepared using the NEBNext® Enzymatic Methyl seq kit (New England BioLabs® Inc., Ipswich, MA, USA) according to the manufacturer’s instructions. Each sample was prepared using 100 ng input DNA in 48 μL TE buffer (1 mM Tris-HCl; 0.1 mM EDTA; pH 8.0) with 1 μL spikes of both the CpG unmethylated Lambda and CpG methylated pUC19 control DNA provided in the kit. The samples were sonicated using a Covaris® S220 focused-ultrasonicator in microTUBE AFA Fiber Pre-Slit Snap-Cap 6×16 mm tubes (Covaris®, Woburn, MA, USA) with the following program parameters: peak incident power (W) = 140; duty factor = 10%; cycles per burst = 200; treatment time (s) = 80. Following library preparation, library concentration and quality were assessed by fluorometry using a Qubit™ 4 and Qubit™ 1X dsDNA HS Assay Kit (ThermoFisher Scientific) and by electrophoresis using a TapeStation (Agilent, Santa Clara, CA, USA). Library concentration was further quantified by qPCR using the NEBNext® Library Quant Kit for Illumina® (New England BioLabs® Inc.). Libraries were equimolarly pooled in batches of ∼15 (five libraries per age cohort) and cleaned using an equal volume of NEBNext® Sample Purification Beads (New England BioLabs® Inc.). The library pools were eluted in 25 μL TE buffer (1 mM Tris-HCl; 0.1 mM EDTA; pH 8.0), and concentration and quality were assessed by fluorometry and electrophoresis as above. Library pools were sequenced on two lanes of the Illumina® HiSeq4000 platform to generate 150-bp paired-end reads.

### small RNA-Seq Library Preparation and Illumina Sequencing

The NEBNext Small RNA Library Prep Set for Illumina was used according to the manufacturer’s instructions. Briefly, 3’ and 5’ adaptors were ligated to the 3’ and 5’ ends of the small RNA, respectively. Then the first-strand cDNA was synthesized after hybridization with a reverse transcription primer. The double-stranded cDNA library was generated through PCR enrichment. After purification and size selection, libraries with insertions between 18∼40 bp were ready for single-end sequencing 50-nt read length with Illumina NovaSeq, 10 million raw reads per sample.

Detailed analysis methods for enzymatic methyl-seq libraries, including identification of differentially methylated regions (DMRs) and annotation of genetic features and transposable elements, and analysis of small RNA libraries, including examining differential expression, are presented in Methods S1. The general methodology, including software packages and references used for this study, is illustrated in Figure 1.

**Fig 1.**
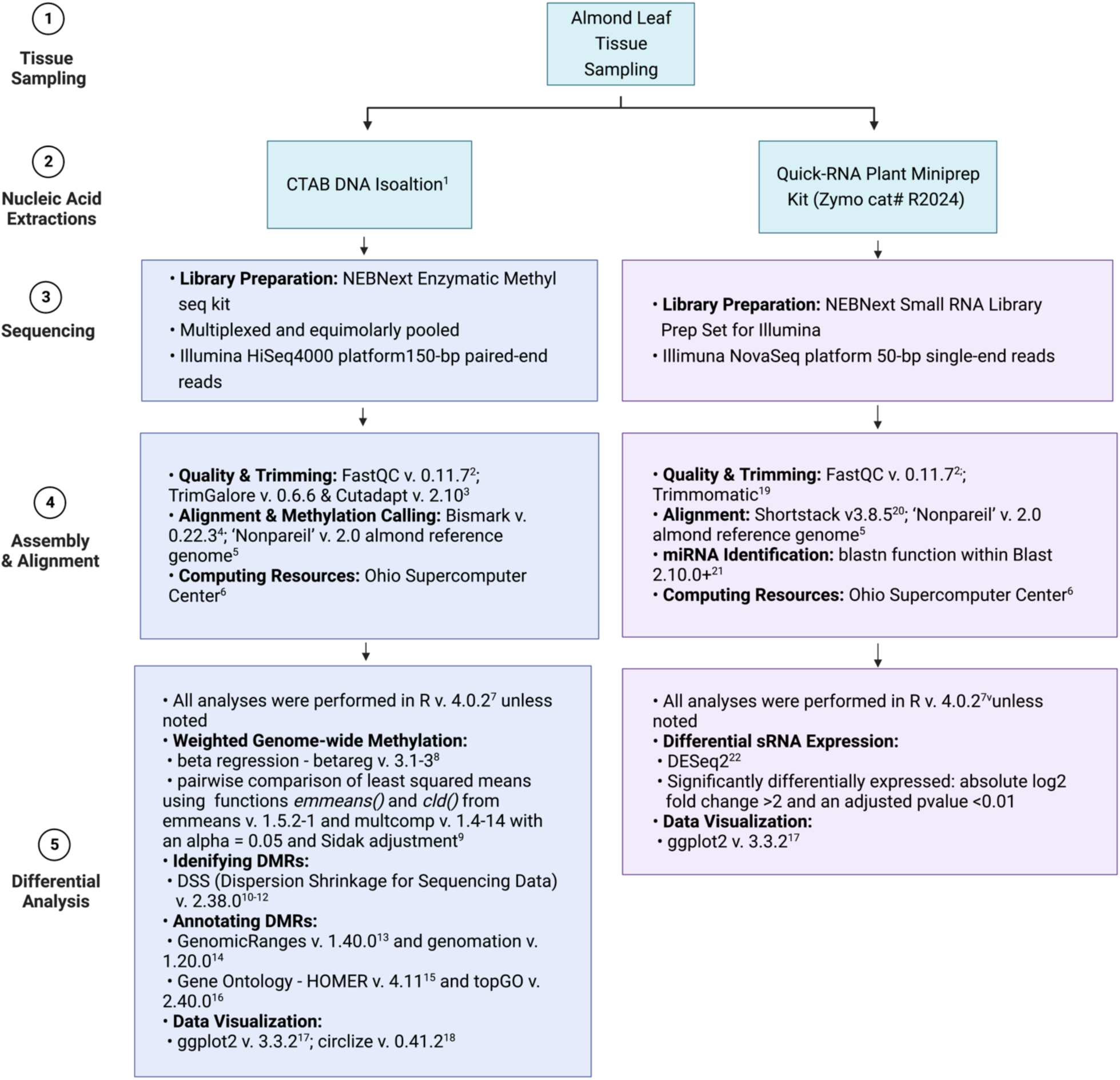
Pipeline of methodology following almond leaf tissue sampling, including DNA and RNA isolations, sequencing library preparation, Illumina sequencing platform, and data analysis approaches and tools. (^1^Vilanova *et al*., 2020; ^2^Andrews, 2010; ^3^Krueger, 2016; ^4^Krueger & Andrews, 2011; ^5^D’Amico-Willman *et al*., 2022; ^6^Ohio Supercomputer Center, 1987; ^7^R Core Team, 2020; ^8^Cribari-Neto & Zeileis, 2010; ^9^Hothorn *et al*., 2008; ^10^Wu *et al*., 2013; ^11^Feng *et al*., 2014; ^12^Park & Wu, 2016; ^13^Lawrence *et al*., 2013; ^14^Akalin *et al*. 2015; ^15^Heinz *et al*., 2010;^16^Alexa A, 2020; ^17^Wickham, 2016; ^18^Gu *et al*., 2014; ^19^Bolger, 2014; ^20^Johnson *et al*., 2016;^21^Camacho, 2009; ^22^Love *et al*., 2014) (Image created with BioRender.com)

## Results

### Genome-wide methylation analysis in almond accessions representing three age cohorts

Following DNA isolation, library preparation, and Illumina sequencing, a total of 21 almond breeding selections were used for subsequent analysis in the 2-year-old age cohort, 25 in the 7 year-old age cohort, and 24 in the 11-year-old age cohort. Sequencing results show aligned coverage for almond accessions ranged from 3.85 – 50.41X (**average:** 12.7 X; **median:** 11.3X) with an average of 49.8 % reads uniquely aligned reads (Data S1). Conversion efficiency was greater than 98% based on alignment to the Lambda reference sequence file (Data S1).

Analysis of weighted genome-wide percent methylation within all methylation contexts (CG, CHG, and CHH) revealed a significant increase in weighted methylation in the 11-year-old age cohort compared to the 2-year-old in the CG (p-value = 0.0105) and CHH (p-value = 0.0399) contexts, respectively (Fig. 2a, c). There was also a significant increase in CG methylation in the 11-year-old age cohort compared to the 7-year-old age cohort (p-value = 0.0115; Fig. 2a). There was not a significant difference in weighted genome-wide methylation in the CHG context when comparing age cohorts (Fig. 2b).

**Fig 2.**
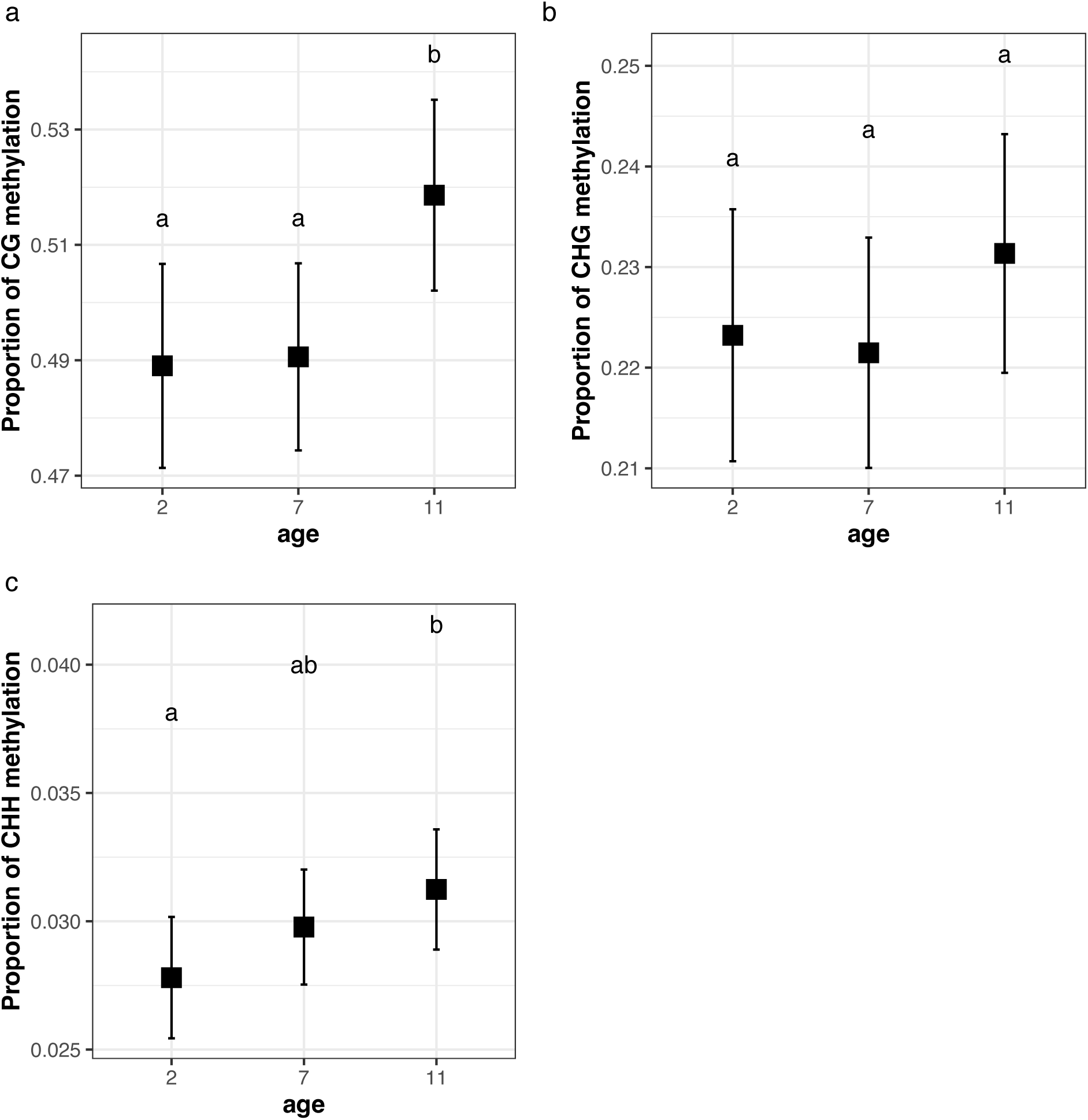
Proportion of weighted genome-wide methylation in the CG **(a)**, CHG **(b)**, and CHH **(c)** methylation-contexts for each age cohort (2, 7, and 11 years old). Letter groups represent significant differences based on pairwise comparisons using least squared means (alpha = 0.05).

To further analyze weighted methylation in these samples, methylation data for each individual was processed per chromosome, and weighted methylation was analyzed at the chromosome level for each methylation context. (Fig. S1a-c). Pairwise comparisons of DNA methylation within each chromosome revealed significant differences in cytosine methylation on distinct chromosomes for each methylation context (Table S1). In the CG context, both the 2 – 11 year and the 7 – 11 year age contrasts were significant on chromosomes 1, 3, 5, 7, and 8 (Table S1). In the CHG context, both the 2 – 11 year and the 7 – 11 year age contrasts were significant on chromosome 5, the 7 – 11 year age contrast was significant on chromosome 7, and the 2 – 11 year age contrast was significant on chromosome 8 (Table S1). Finally, in the CHH context, the 2 – 11 year age contrast was significant on chromosomes 5, 7, and 8 (Table S1). Overall, significant differences in chromosome-level DNA methylation between age cohorts tend to occur on chromosomes 5, 7, and 8.

### Identification and classification of differentially methylated regions (DMRs) between age cohorts

DMRs were identified based on comparisons between the age cohorts in each methylation context. Most DMRs identified are in the CG context, followed by CHH and CHG, respectively (Table 1). These DMRs were further classified as hyper- and hypomethylated where hypermethylated indicates the older cohort is more methylated than the younger cohort and hypomethylated indicates the older cohort is less methylated than the younger cohort for each DMR. In the CG context, 96%, 94%, and 64% of the identified DMRs were hypermethylated in the older trees for the 11 – 2 year, 11 – 7 year, and 7 – 2 year age contrasts, respectively (Table 1). In the CHG context, 68%, 52%, and 64% of DMRs were hypermethylated in the older trees for the 11 – 2 year, 11 – 7 year, and 7 – 2 year age contrasts, respectively (Table 1). Finally, in the CHH context, 82%, 38%, and 82% of DMRs were hypermethylated in the older trees for the 11 – 2 year, 11 – 7 year, and 7 – 2 year age contrasts, respectively (Table 1). The cumulative binomial probability of the occurrence of hypermethylated DMRs was less than 1×10^-6^ for all age contrasts except 11 – 7 in the CHG and CHH contexts, suggesting there are more hypermethylated DMRs than would be expected given an equal probability of hyper- and hypomethylated DMRs in the genome. Identified DMRs ranged in length from 51- 4,824 base pairs, with an average length of 200 base pairs and a median of 139 base pairs (Fig. S2a-l) The average length of CG, CHG, and CHH DMRs is 212, 201, and 177 base pairs, respectively. The average length of a gene in the ‘Nonpareil’ genome is 2,912 bp, so most of the identified DMRs are much shorter than the average gene.

**Table 1.** Number of identified differentially methylated regions (DMRs) in each methylation context when comparing the three age cohorts. DMRs were identified with a threshold of p :: 0.0001. DMRs are classified as hypermethylated if the percent methylation in that region is greater in the older age cohort compared to the younger age cohort within each contrast. DMRs are classified as hypomethylated if the percent methylation in that region is lesser in the older age cohort compared to the younger age cohort within each contrast. Hypermethylated DMR values in bold represent those with a cumulative binomial probability < 1×10^-6^.

| Methylation Context | Contrast | Number of DMRs | Hypermethylated DMRs | Hypomethylated DMRs |
| --- | --- | --- | --- | --- |
| CG | 11-2 | 6479 | **6232** | 247 |
|  | 11-7 | 2105 | **1983** | 122 |
|  | 7-2 | 1211 | **985** | 226 |
| CHG | 11-2 | 1129 | **763** | 366 |
|  | 11-7 | 485 | 252 | 233 |
|  | 7-2 | 813 | **524** | 289 |
| CHH | 11-2 | 2059 | **1690** | 369 |
|  | 11-7 | 567 | 216 | 351 |
|  | 7-2 | 2259 | **1849** | 410 |

The distribution of CG-context DMRs showed a similar pattern across all chromosomes, where the 11 – 2 year age contrast has the highest number of DMRs per chromosome, followed by the 11 – 7 year and 7 – 2 year age contrasts (Fig. S3a). In the CHG and CHH contexts, the distribution of DMRs showed greater variability, with the 11 – 7 year age contrast typically showing the lowest number of DMRs across all chromosomes, while the 11 – 2 and 11 – 7 year age contrasts oscillate in number of DMRs occurring on each chromosome across the genome (Fig. S3b,c).

### Classification of DMRs as hyper- or hypomethylated in the age cohort comparisons

Using the classifications of hyper- and hypomethylated, DMRs were plotted across the eight chromosomes of the ‘Nonpareil’ genome revealing unique distributions based on both methylation context and age contrast, as well as indicating DMR enrichment in specific chromosomes (Fig. 2). In the CG context, DMR enrichment occurs in the 11 – 2 year age contrast, with predominantly hypermethylated DMRs, though enrichment of hypomethylated DMRs appears on chromosome 5 (Fig. 2a). The CHG context represents the lowest overall enrichment of DMRs compared to the other methylation contexts with regions throughout the genome showing a concentrated enrichment of DMRs across the genome (Fig. 2b). Finally, DMRs in the CHH context show similar patterns in enrichment for both the 7 – 2 year and 11 – 2 year age contrasts, with evident DMR enrichment occurring on chromosomes 3 and 8 (Fig. 2c). The 11 – 7 year age contrast in the CHH methylation context is the only contrast to have a higher number of hypomethylated DMRs compared to hypermethylated (Table 1; Fig. 2c).

Following the classification of DMRs as either hyper- or hypomethylated in each age contrast, DMRs were compared among age contrasts to identify overlapping genomic regions. This analysis revealed several overlapping DMRs from distinct age contrasts. The highest number of overlaps occurred in CG context hypermethylated DMRs, particularly when comparing the 11 – 2 by 11 – 7 and 11 – 2 by 7 – 2 age contrasts (Table 2). Interestingly, the 11 – 7 by 7 – 2 age contrast revealed very few overlaps in hypermethylated DMRs, and no overlapping hypomethylated DMRs (Table 2). Finally, a comparison was performed to identify DMRs with overlapping genomic regions among all three age contrasts, showing the 11 – 2 age contrast contains DMRs that share a genomic region with the overlapping DMRs in the 11 – 7 by 7 – 2 comparison (Table 2). This final analysis revealed 17 overlapping DMRs among the three age contrasts, meaning these DMRs share overlapping genomic coordinates (Table S2). These 17 DMRs are the longest (ranging from 106 – 1,504 base pairs) in the 11 – 2 age contrast, followed by the 11 – 7 (56 – 713 base pairs) and 7 – 2 (52 – 1,037 base pairs) age contrasts (Table S2). Analysis of the average percent methylation of cytosines within the genomic regions identified in the 11 – 7 age contrasts show that in most of these regions, cytosines become more methylated with increased age across the three age cohorts (Figs. S4a-c, S5a-m, S6a-c).

**Table 2.** Number of occurrences of overlap when comparing differentially methylated regions (DMRs) identified in each contrast to those identified in the other contrasts. The number of overlaps means the number of times a DMR in a particular age-contrast (e.g., 11-2) overlaps the genomic region of a DMR in one of the other age-contrasts (e.g., 11-7). The overall comparison indicates the number of DMRs occurring in overlapping genomic regions in all three contrasts. DMRs are classified as either hyper- or hypomethylated in each methylation context.

| Methylation Context | Hyper/Hypo | 11-2 * 11-7 | 11-2 * 7-2 | 11-7 * 7-2 | 11-2 * 11-7 * 7-2 |
| --- | --- | --- | --- | --- | --- |
| CG | Hyper | 677 | 646 | 13 | 13 |
|  | Hypo | 37 | 58 | 0 | 0 |
| CHG | Hyper | 75 | 161 | 3 | 3 |
|  | Hypo | 74 | 62 | 0 | 0 |
| CHH | Hyper | 95 | 607 | 1 | 1 |
|  | Hypo | 78 | 146 | 0 | 0 |

### Annotation of hyper- and hypomethylated differentially methylated regions (DMRs)

Annotation was performed by classifying all hyper- and hypomethylated DMRs in each methylation context into four categories (gene, exon, 5’ untranslated region [UTR], and 3’ UTR) based on their association with features in the ‘Nonpareil’ genome annotation. CG context DMRs generally tended to have higher associations with genes and exons compared to the other methylation contexts, while CHG DMRs tended to have higher associations with 5’ UTRs and CHH DMRs with 3’ UTRs compared to the other contexts (Table S3).

Identified DMRs were then annotated using the ‘Nonpareil’ genome annotation file to determine the closest gene associated with each DMR. Enrichment analysis was performed for both hyper- and hypomethylated DMR-associated genes in all methylation contexts for each age contrast, revealing a suite of biological process, molecular function, and cellular component gene ontology (GO) terms associated with each contrast (Tables S4-S9). Comparing annotations between age contrasts identified GO terms unique to each age contrast in each methylation context and degree of methylation (i.e., hyper or hypo). For example, a subset of genes associated with hypermethylated DMRs in the CG context from all three age contrasts was assigned the molecular function GO terms transmembrane transporter activity, protein serine/threonine kinase activity, and DNA-binding transcription factor activity (Table S4b).

### Annotation of genes associated with 17 hypermethylated DMRs identified in all three age cohort age contrasts

Of the DMRs identified in each age contrast, 17 hypermethylated DMRs were found to share genomic regions in all three age contrasts, meaning these regions showed consistent significant increases in methylation in the older age cohort relative to the younger in each age contrast. The 17 DMRs were annotated using the ‘Nonpareil’ genome annotation to identify the closest associated gene. In total, eight previously annotated genes, including *FAR1-RELATED SEQUENCE 5* (*FRS5*), a receptor-interacting serine/threonine-protein kinase 4 (*Ripk4*), and dCTP pyrophosphatase 1 (*Dctpp1*) were identified as associated with nine of the DMRs (Table 3). One CG DMR and one CHG DMR are associated with the same gene, Tryptophan aminotransferase-related protein 3 (*TAR3*) (Table 3). The remaining eight DMRs are associated with genes of unknown function (Table 3). These eight unknown protein sequences were used as input into three programs to determine properties, including predicted motifs, localization, and weight. Two of these unknown proteins contained transposase_24 motifs, and four are predicted to be localized to the nucleus (Table S10).

**Table 3.**
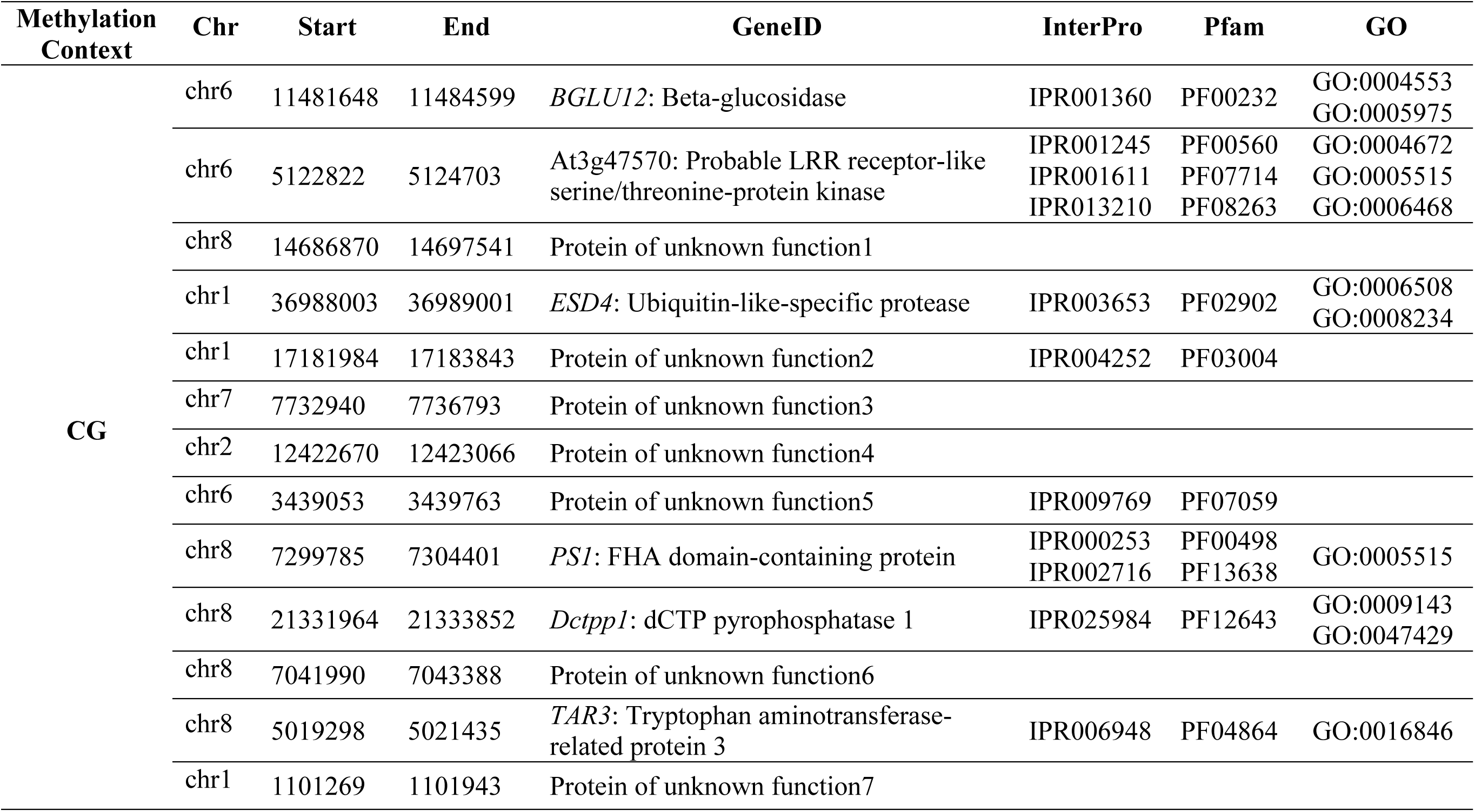
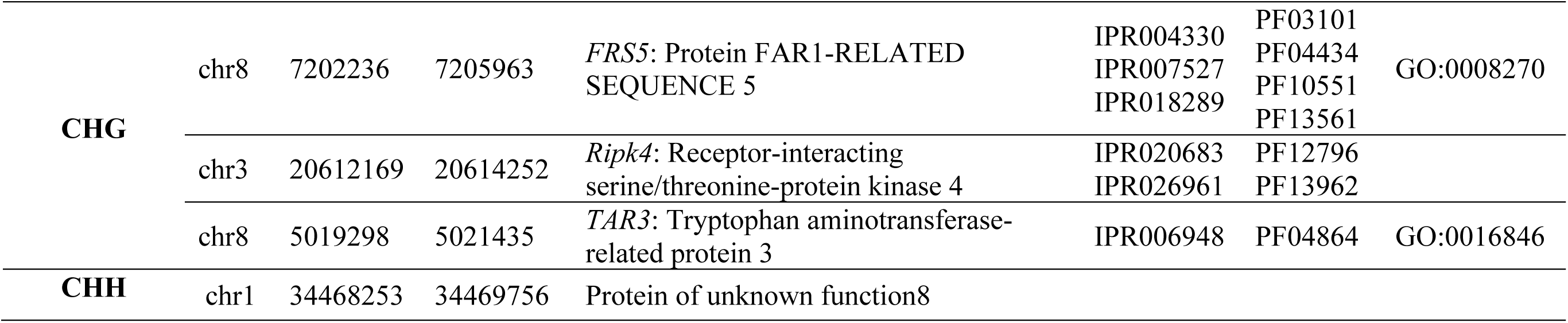
Annotation of genes associated with 17 hypermethylated differentially methylated regions (DMRs) occurring in all three age cohort contrasts. The chromosome (chr) and genomic coordinates (start and end) of each gene are listed along with the gene identification from the ‘Nonpareil’ genome annotation. Protein identifiers from InterPro and Pfam databases are also included, as well as gene ontology (GO) terms associated with the gene.

In addition to identifying genes associated with these 17 shared DMRs, the DMR genomic sequences (Data S2) were searched against the modified miRBase database using blastn to identify any potential miRNAs within these regions. Results from this analysis showed that 12 of the DMRs contained at least one previously identified miRNA sequence (Table S11).

### Profiling of DMRs overlapping transposable elements in two age cohorts

The 2- and 7-year-old age cohorts were used to determine overlap of DMRs with transposable elements in the almond genome, as this contrast had the largest sample size and represents the typical transition from juvenile to mature adult in almond (occurring ∼ 5 years old). Of the DMRs identified in the 2-7 age contrast, 493 overlapped with 495 predicted transposable elements (Data S3). These overlaps occurred among transposable elements in 11 categories (Fig. S7. Out of those transposable elements with a defined taxonomy, the majority of these overlaps occurred within Mutator Terminal Inverted Repeat (TIR) transposons (132) and Long Terminal Repeat (LTR) retrotransposons (87). The smallest proportion of DMRs occurred within Long Interspersed Nuclear Elements (LINEs). The most common methylation context occurring within transposable elements was CG (252), followed by CHG (156) and CHH (87). The length of the DMRs varied from 50 to 978 base pairs, and the overlap with transposable elements varied between 41 to 381 base pairs, with 90% of the overlaps occurring between 1 and 121 base pairs and 95% of the transposable elements being between 71 and 1171 base pairs long. Interestingly, most of the DMRs overlapping with transposable elements occurred along chromosome 4 (91), followed by chromosome 2 (71), chromosome 1 (70) and chromosome 8 (67), chromosome 3 (56), chromosome 7 (48), chromosome 6 (46) and chromosome 5 (42); four additional DMRs occurred in unassembled contigs. This pattern deviates from the expectation that longer chromosomes tend to have more transposable elements and, therefore, higher chances of overlapping with DMRs.

### Profiling small RNA expression and identifying differentially expressed small RNAs in two age cohorts

Small RNA expression was profiled in two almond breeding selection cohorts aged 1 and 6 years at the time of sampling. Three individual almond breeding selections were sequenced from each of the two cohorts. Analysis of the sRNA sequencing data by principal component analysis revealed clustering based on age in 20, 21, 23, and 24 nucleotide sRNAs, with the age cohort explaining 30%, 35%, 36%, and 37% of variance in the small RNA expression profiles, respectively (Fig. 4). Differential expression analysis showed 1,768 significantly differentially expressed sRNAs, and 1,450 of those have a negative log2fold change value, indicating overall down regulation in the 6-year-old compared to the 1-year-olds. This trend is particularly apparent in the 21 and 22 nucleotide sRNAs, where most of the significantly differentially expressed sRNAs are downregulated in the 6-year-olds (Fig. 5). Interestingly, the 23 and 24 nucleotide sRNAs show the opposite pattern, where most of the significantly differentially expressed sRNAs are upregulated in the 6-year-olds compared to the 1-year-olds (Fig. 5). None of the 20 nucleotide sRNAs show significant differential expression between the two age cohorts (Fig. 5).

**Fig 3a-c.**
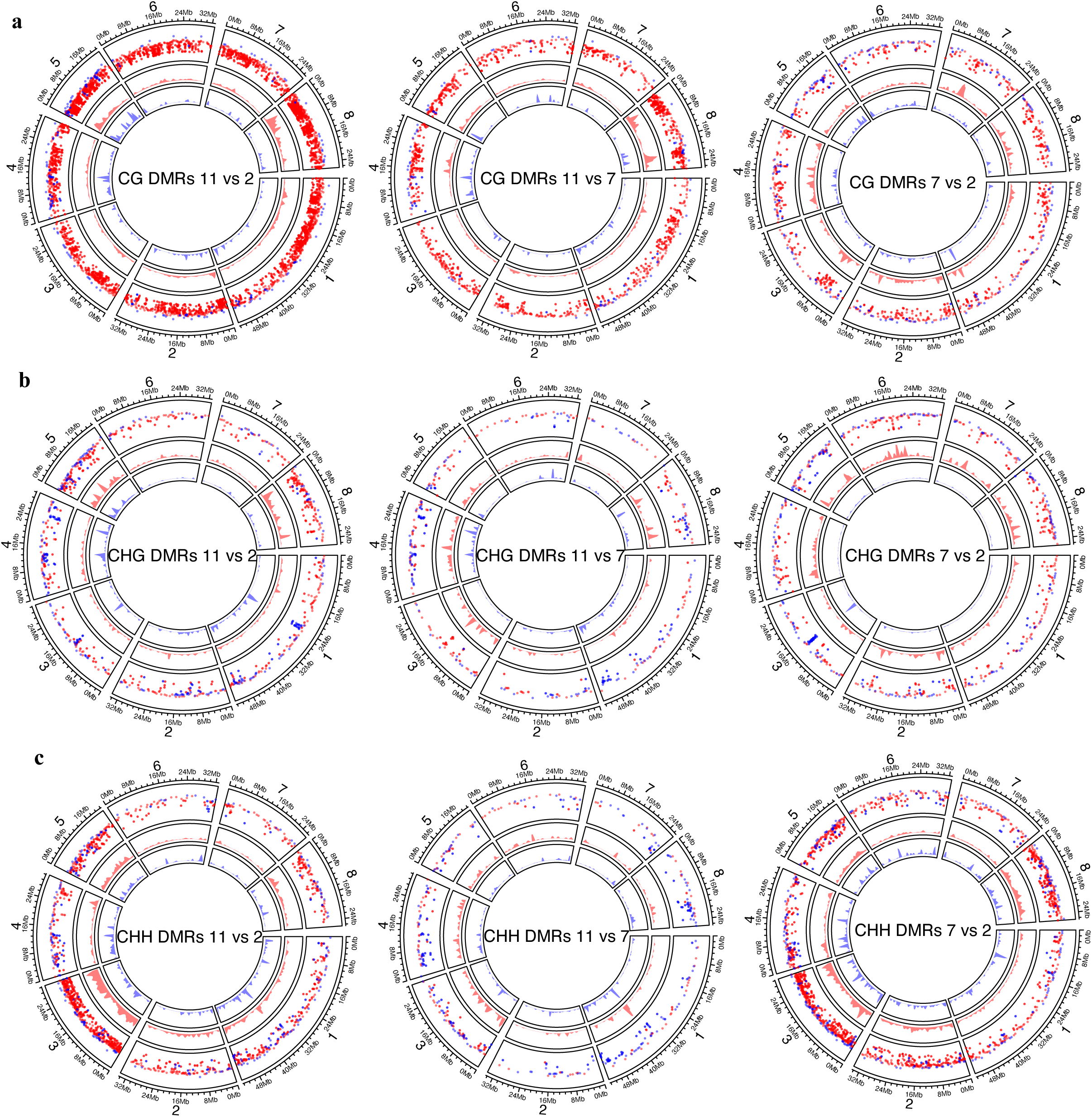
Circos plots depicting individual hyper (red) and hypo (blue) methylated differentially methylated regions (DMRs) identified in each contrast and methylation-context. The outer ring of each plot gives approximate location of the individual DMRs on each of the eight ‘Nonpareil’ chromosomes represented by red and blue dots. The middle ring of each plot represents enrichment of hypermethylated DMRs across each chromosome, and the innermost ring of each plot represents enrichment of hypomethylated DMRs across each chromosome. Panel **a** shows the distribution of DMRs in the CG context, panel **b** shows distribution of DMRs in the CHG context, and panel **c** shows distribution of DMRs in the CHH context.

**Fig 4.**
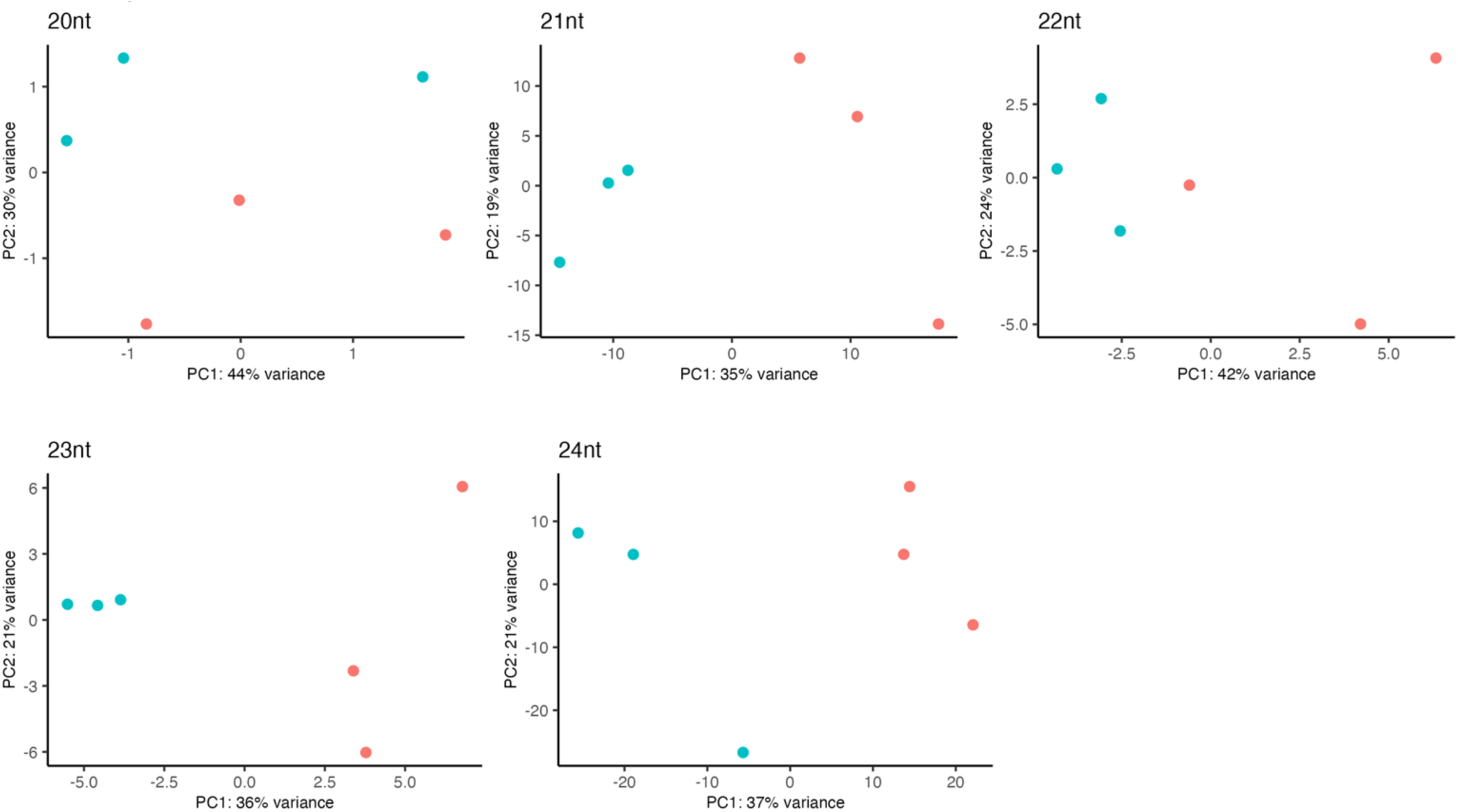
Analysis of small RNA expression profiles using principal component analysis following rlog transformation subset by the small RNA size (20, 21, 22, 23, and 24 nucleotides [nt]). All small RNA sizes with the exception of 22 nt show a cluster of 1-year-old individuals (pink) and 6-year-old (teal) individuals.

**Fig 5.**
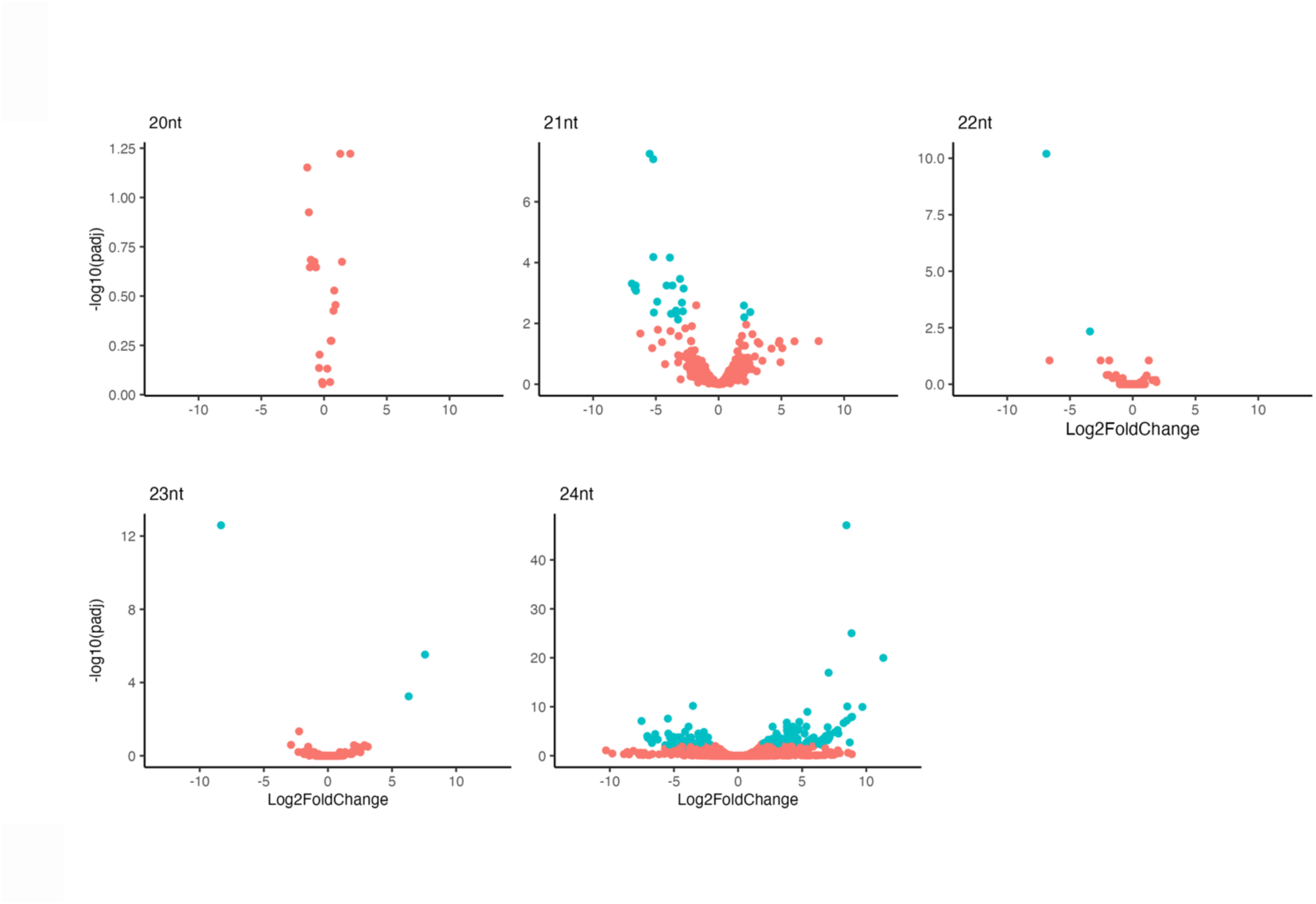
A volcano plot shows differential expression patterns of individual small RNAs based on size (20, 21, 22, 23, and 24 nucleotides [nt]), where a positive fold change indicates upregulation of small RNAs in the 6-year-old cohort compared to the 1-year-old cohort. Pink dots represent small RNAs that are not significantly differentially expressed, and teal dots represent small RNAs that are significantly differentially expressed (p < 0.01).

Subsequent BLAST analysis of the sRNAs against the miRbase database showed significant alignment of 455 sRNAs to previously identified microRNA families. Among those, 18 are significantly differentially expressed and align to a total of 9 microRNA families within the miRbase database (Fig. 6). The microRNA families include miR1511, mir159, miR171, miR395, miR477, miR482, miR5742, miR6274, and miR7125 (Fig. 6).

**Fig 6.**
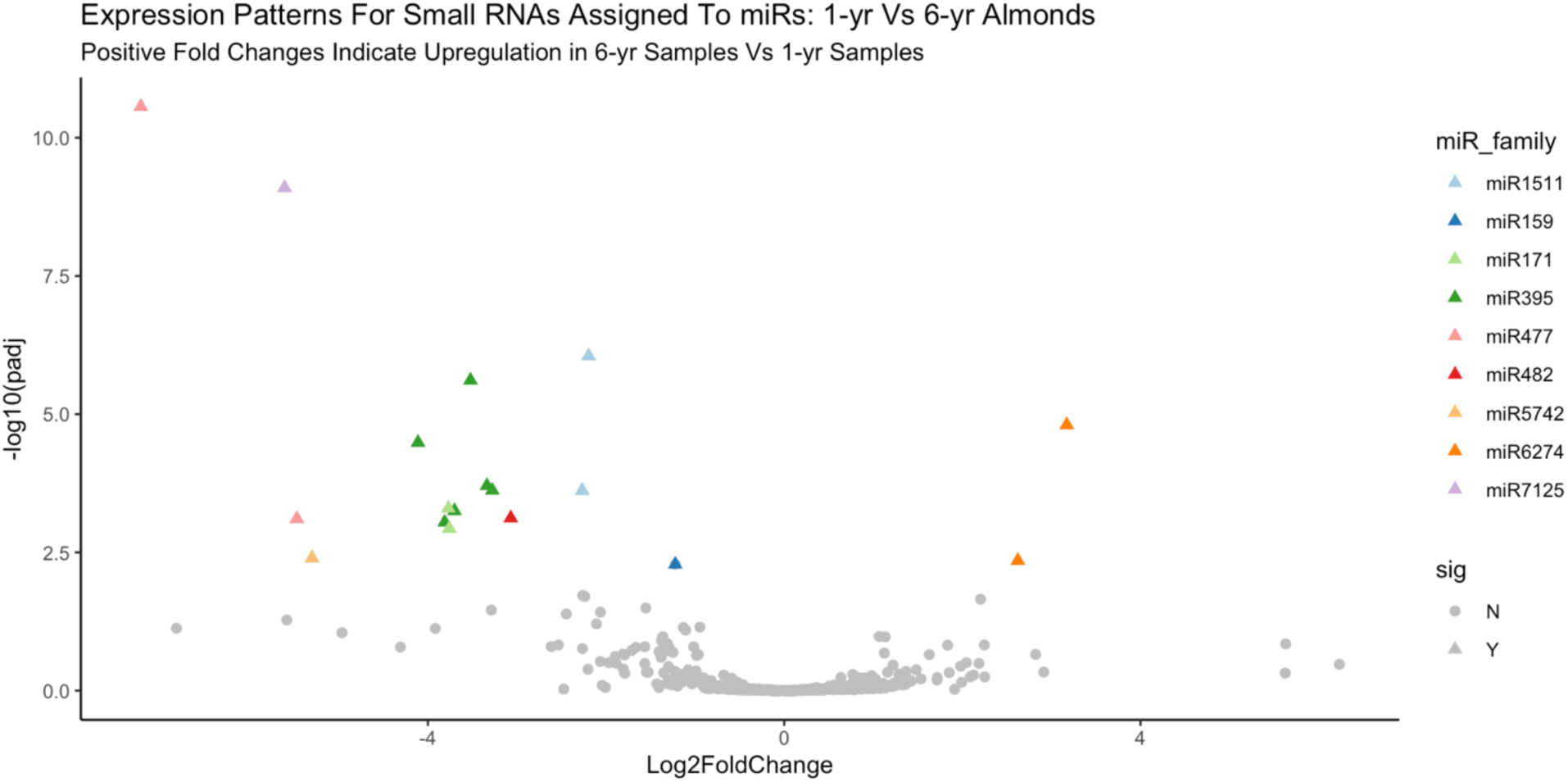
Volcano plot depicting the 455 differentially expressed small RNAs that significantly align to known microRNA families in the miRbase database based on BLAST analysis. Significantly differentially expressed small RNAs are identified as triangles, and the associated microRNA family based on significant BLAST alignment is indicated by color of the triangle.

## Discussion

Perennial plant aging and the impacts of this process, particularly on productive fruit and nut crops, is a neglected area of research with potential applications for agricultural production and crop improvement. The ability to track age in clonally propagated crops could aid in the mitigation of age-related disorders like non-infectious bud failure (Kester *et al*., 2004) and, more broadly, in overcoming the decrease in plant performance resulting from intense production systems that affect orchard/vineyard longevity. Biomarkers of age in these species, however, are lacking. The aim of this study was to test the hypothesis that, on average, almond breeding selection cohorts will exhibit divergent DNA methylation profiles associated with age. The long term goals of this work are to develop biomarkers of age in almond, a clonally propagated crop, and to further our understanding of the aging process in perennial species. To address this, whole-genome DNA methylation profiles were generated for ∼70 almond individuals from three distinct age cohorts, and comparisons were made between cohorts to identify regions of interest for further study into their involvement in the aging process and utility as biomarkers of age in almond.

### DNA hypermethylation in the CG and CHH contexts is associated with increased age in almond

The DNA methylation profiles generated for individuals in the three age cohorts (11, 7, and 2 years old) were compared at the whole-genome and chromosome level, which showed that hypermethylation in the CG and CHH methylation contexts is associated with increased age in almond. Further, the probability of identifying the number of hypermethylated DMRs observed in this study was very low for most age contrasts, suggesting there was a disproportionately high number of hypermethylated DMRs identified compared to hypomethylated DMRs. This result supports previous work theorizing an increase in total genomic DNA methylation with increased age in plants (Dubrovina & Kiselev, 2016). Previous studies have also shown that genome-wide hypermethylation can result in a high number of identified hypermethylated DMRs in subsequent analyses, as was reported in several species, including Monterey pine (*Pinus radiata* D.Don), peach (*P. persica*), and coast redwood (*Sequoia sempervirens* [D.Don] Endl.) (Bitonti *et al*., 2002; Fraga *et al*., 2002b; Huang *et al*., 2012).

DNA methylation has been proposed as a “biological clock” capable of predicting the true, ontogenetic age of an individual due to observed patterns of increased methylation with increased age in a variety of species (Runov *et al*., 2015). Results in this study suggest that almond fits this pattern of hypermethylation, and thus DNA methylation may serve as a biomarker of age in this species. Whole-genome hypermethylation with increased age represents an opportunity to develop high-throughput screening methods that do not require whole-genome sequencing. These methods could include high-performance liquid chromatography or capillary electrophoresis (Stach *et al*., 2003; Armstrong *et al*., 2011).

Regulation by DNA methylation, histone modifications, and chromatin remodeling has also been shown to modulate the juvenile-to-adult phase transition in plants, including in gymnosperms like Monterey pine and angiosperms such as peach (Bitonti *et al*., 2002; Fraga *et al*., 2002a; Xu *et al*., 2018). The juvenile period in almond is approximately 3-4 years; thus, the differential patterns of methylation observed in this study between the 2- and 7-year cohorts and the 2- and 11-year cohorts could be associated with the juvenile-to-adult transition as has been documented in other plants (Dubrovina & Kiselev, 2016). Patterns of differential methylation identified in this study and associated with specific regions of the genome further demonstrate the potential involvement of DNA methylation in regulating this transition in almond. Further investigation is needed focusing on the involvement of DNA-methylation in the juvenile-to-adult transition in almond and other perennial species, including those with available transgenic germplasm exhibiting reduced juvenility, such as apple (Flachowsky *et al*., 2011; Kumar *et al*., 2020).

### Differentially methylated regions (DMRs) in the CG and CHH contexts are enriched on specific chromosomes in the almond genome

Following the identification of DMRs in the three age contrasts, these regions were plotted across the almond genome showing enrichment of DMRs on specific chromosomes, particularly in the CG and CHH methylation contexts. These so-called “hotspots” of differential DNA methylation could suggest loci in these regions are prone to chromatin alterations, including methylation. Transposable elements (TEs) tend to be heavily methylated and have been reported to be involved in developmental processes in almond, such as the juvenile-to-adult transition (Han *et al*., 2018; Corso-Díaz *et al*., 2020; Wyler *et al*., 2020). In our study, we found that in the 2-7 age contrast, there is a high proportion of TEs overlapping with DMRs in chromosomes 3, 4, 7, and 8, which corresponds with DMR enrichment hotspots observed in the age cohort comparisons. This type of pattern has been documented in other species, such as rice, where a study on salt tolerance identified DMRs that tended to cluster on specific chromosomes and were typically associated with TEs on these chromosomes (Ferreira *et al*., 2019).

In this study, hypermethylated CG DMRs were enriched on chromosome 8, and hypermethylated CHH DMRs on chromosomes 3 and 8. As we have shown, increased levels of methylation tend to occur in regions rich in TEs, which seem to become increasingly methylated with age. A study in almond characterized the TE landscape in the ‘Texas’ cultivar and compared the distribution of TEs in the genome to that of peach (Alioto *et al*., 2020). This study revealed not only an increased involvement of TEs in trait diversity in almond compared to peach but also showed enrichment of TEs on almond chromosomes 3 and 8 (Alioto *et al*., 2020), corresponding to the chromosomes identified in this study to show enrichment of hypermethylated CG and CHH DMRs. Chromosomes 3 and 8 are targets for further exploration of biomarkers of age and to understand their association with the juvenile-to-adult transition in almond.

Interestingly, a study in *Brachypodium distachyon* (L.) P.Beauv. found that DMRs were highly correlated with genetic diversity as classified by the presence of single nucleotide polymorphisms (SNPs) throughout the genome (Eichten *et al*., 2016). This genetic diversity was found to be related to the presence of TEs at these sites, potentially contributing to the formation of SNPs as well as leading to differential levels of methylation between the lines tested (Eichten *et al*., 2016). Given the heterozygosity and diversity in almond germplasm, it may be relevant to compare regions enriched in DMRs from the age contrasts to SNP data in almond to test for a correlation between increased methylation and genetic diversity, particularly for traits associated with growth and development and length of juvenility.

### DMRs as potential biomarkers of age in almond

The DMRs identified in this study represent those regions in the genome that showed either an increase or a decrease in cytosine methylation with increased age in almond. The results herein suggest that DMRs tend to be hypermethylated in older trees. This pattern fits with the weighted genome-wide methylation patterns showing significant increases in methylation in the CG and CHH contexts between the 2- and 11-year-old age cohorts. Unique, hypermethylated DMRs were identified in each age contrast, providing information on DNA methylation dynamics associated with age. Interestingly, no hypomethylated DMRs were identified in each age contrast, suggesting that hypermethylation is more prominent in aging in almond. Regions that show increased methylation in each age contrast are of particular interest due to their potential suitability as biomarkers of age since these regions show incremental increases in methylation from 2-to-7 years and again from 7-to-11 years old. There were 17 hypermethylated DMRs common to all three contrasts that may be particularly useful as biomarkers. Once these regions are validated via DNA methylation profiling in additional almond cohorts of known age, a predictive model based on these regions could be developed and applied to clonal germplasm to predict ontogenetic age, providing a basis to screen germplasm for susceptibility to undesirable, age-related phenotypes. These tools may have implications for germplasm management in breeding, production (orchard), propagation (nursery), and conservation (repository) settings.

The genetic features associated with these specific DMRs could also be involved in developmental processes, including the juvenile-to-age transition. Of the 17 DMRs, nine were found to be associated with eight annotated genes. These genes include *FRS5*, a *FAR1-related* protein in the *FAR1* gene family which is involved in light perception and was demonstrated to be involved in plant development and regulation of aging processes in Arabidopsis (Lin & Wang, 2004; Ma & Li, 2018; Xie *et al*., 2020). Further, *FAR1-related* genes have been previously found to be domesticated TEs (Lin *et al*., 2007). In the ‘Nonpareil’ genome annotation, 528 genes are found to have the annotation of *FAR1-related*. Given the occurrence of this annotation in the genome, this suggests that these genes may also be domesticated TEs. The gene *TAR3* was associated with two DMRs, one in the CG context and one in the CHG context. This gene is part of the *TRYPTOPHAN AMINOTRANSFERASE (TAR)* gene family, whose members are a component of one of the major auxin biosynthetic pathways (Hofmann, 2011). Auxin is a well-known regulator of plant development and senescence processes (Ljung, 2013; Khan *et al*., 2014; Mueller-Roeber & Balazadeh, 2014), and disruption of *TAR3* expression could impede auxin production (Hofmann, 2011). Two receptor-like kinase (RLK) genes were also found to be associated with DMRs in the CG and CHG context. RLKs have been extensively studied in plants and are known to be involved in cell signaling and related to plant defense and plant developmental processes (Afzal *et al*., 2008). Additionally, eight proteins of unknown function were associated with the overlapping DMRs. These genes and the others identified represent interesting targets for future study on their potential involvement in aging processes in almond, including in the vegetative transition.

### Putative role of miRNAs in almond aging

In addition to identifying nearby genes associated with these DMRs, microRNAs (miRNAs) were also surveyed in these regions. CGDMR5 contained two putative miRNA sequences with best hits to *Glycine max* miR172e and miR172d, members of the miR172 family of microRNAs. The miR172 family are major regulators of development and phase transition in plants (Wu *et al*., 2009; Tang *et al*., 2018; Zhang & Chen, 2021). MiR172 increases in abundance with age, acting antagonistically with miR156 to regulate phase transition from vegetative to reproductive in plants (Auckerman & Sakai, 2003; Bastías *et al*., 2016). If miR172 activity is involved in the juvenile-to-adult transition in almond, as suggested in other plants, significant upregulation of this miRNA would be expected in the 6-year-old cohort compared to the 2-year-old cohort in this study. The sRNA expression analysis in this study revealed several significantly differentially expressed miRNAs; however, miR172 was not found to be significantly differentially expressed. It is possible that expression was not detected during the time points sampled because the miRNA is not active or because the correct time points were not chosen. Further work is needed in almond to examine miR172 activity, including expression, during the vegetative transition to understand its involvement in aging. Among the differentially expressed small RNAs, miR171 was found to be significant. This microRNA has been previously shown to be involved in regulating the GRAS transcription factor family which is involved in regulating flowering and meristem development (Wang *et al*., 2010). Work in barley has shown that miR171 activity is also involved in phase transitions regulated by the miR156/miR172 pathway (Curaba *et al*., 2013).

In addition to specific microRNA expression patterns, overall small RNA expression profiles in the two age cohorts revealed an association between small RNA expression and aging. An association between sRNAs and aging has been identified in other organisms, and small RNA expression has been suggested as a biomarker of aging and aging-associated disorders (Chen *et al*., 2010; Gao *et al*., 2012; Proshkina *et al*., 2020). The correlation between small RNA expression and age in almond suggests that sRNA expression profiles are potential biomarkers of age and could be used along with DMRs to generate models of aging in almond and other *Prunus* species.

Biomarkers of age are valuable for clonally propagated crops, whose ontogenetic age can be difficult to determine, to screen and select germplasm with low potential for developing age related disorders. Further, the DMRs and small RNAs identified in this study can be used to guide future studies aimed at increasing our understanding of plant aging and vegetative phase transition in perennials. Perennial plants, including almond, are known for having long juvenile periods, which can inhibit breeding and improvement efforts. Alteration of DNA methylation or sRNA expression provides another approach for manipulating phase transition and developing perennial crops with shorter juvenile periods, dramatically shortening breeding cycles for these species.

## Acknowledgements

We would like to acknowledge Matthew Willman for his assistance with the statistical analysis and preparation of the scripts used to perform the computational analyses for this manuscript. We would also like to acknowledge the Ohio Supercomputer Center for access to computing resources and the Translational Plant Sciences Graduate Program for the fellowship for KMDW. This work was supported by The Ohio State University CFAES-SEEDS program grant # 2019-125, the Almond Board of California Grant HORT35, the U.S. Department of Health and Human Services National Institutes of Health National Cancer Institute Cancer Center Support Grant (CCSG) P30CA016058, the USDA National Institute of Food and Agriculture AFRI-EWD Predoctoral Fellowship 2019-67011-29558.

## Author contributions

KMDW and JFR conceptualized and designed the study. KMDW and TMG performed the tissue sampling. KMDW and ESA performed all the laboratory portions of the project. KMDW performed all analyses with the assistance of JFR and CEN. KMDW prepared the manuscript with the assistance of JFR and CEN. All authors contributed to editing the manuscript prior to submission. All the authors approved the submission and revised version of this manuscript.

## Data availability

The ‘Nonpareil’ almond reference genome can be found at https://www.rosaceae.org/Analysis/13738196. All sequencing data for this project has been deposited to the NCBI Sequence Read Archive under Bioproject PRJNA933198 including biosamples: SAMN33228162 to SAMN33228237. All code used to perform analyses reported in the manuscript can be found at (https://github.com/kmdamico/AgeCohortAnalysis).

## Processing and Alignment of Enzymatic Methyl-Seq Libraries

Methyl-Seq read quality was initially assessed using FastQC v. 0.11.7 (Andrews, 2010), and reads were trimmed using TrimGalore v. 0.6.6 and Cutadapt v. 2.10 with default parameters (Krueger, 2016). Forward read fastq and reverse read fastq files from the two HiSeq4000 lanes were merged for each library to produce single fastq files for both read one and read two. Reads were aligned to the ‘Nonpareil’ v. 2.0 almond reference genome (D’Amico-Willman *et al*., 2022), deduplicated, and methylation calls were generated using Bismark v. 0.22.3 (Krueger & Andrews, 2011) with default parameters in paired-end mode. To test conversion efficiency, reads were also aligned to both the Lambda and pUC19 nucleotide sequence fasta files provided by NEB (https://www.neb.com/tools-and-resources/interactive-tools/dna-sequences-and-maps-tool).

All analyses were performed using the Ohio Supercomputer Center computing resources (Ohio Supercomputer Center, 1987).

## Weighted Genome-wide Methylation Analysis of Age Cohorts

Weighted genome-wide percent methylation values were calculated for each individual within each cohort by taking the total number of methylated reads at each cytosine and dividing this by the total number of reads (methylated + unmethylated) at each cytosine. Weighted values were calculated for each methylation context. These values were used as input to R v. 4.0.2 (R Core Team, 2020) to perform beta regression using the package betareg v. 3.1-3 (Cribari-Neto & Zeileis, 2010). Pairwise comparison of least squared means was completed by the functions *emmeans()* and *cld()* from the R packages emmeans v. 1.5.2-1 and multcomp v. 1.4-14 with an alpha = 0.05 and Sidak adjustment (Hothorn *et al*., 2008). The R package ggplot2 v. 3.3.2 was used to create plots for weighted percent methylation within each methylation context (Wickham, 2016). Files were then subset by chromosome (chr1 – chr8), and weighted percent methylation values were calculated for all individuals by chromosome using the same formula as above for each methylation context. These values were used as input in R v. 4.0.2 (R Core Team, 2020) to perform beta regression and subsequent pairwise comparison of least squared means as performed above for the genome-wide weighted percent methylation values.

## Differential Methylation Analysis of Age Cohorts

Coverage files for each methylation context produced by Bismark were prepared for input into the R package DSS (Dispersion Shrinkage for Sequencing Data) v. 2.38.0 (Wu *et al*., 2013; Feng *et al*., 2014; Park & Wu, 2016). The functions *DMLtest()* and *callDMR()* were used with a significance p.threshold set to 0.0001 to identify differentially methylated regions (DMRs) through pairwise comparisons between the three age cohorts. Comparisons were made relative to the oldest cohort in each DMR test (i.e., 11-year-old cohort relative to 2-year-old cohort).

Following identification of DMRs in each age contrast (11 – 2 years; 11 – 7 years; 7 – 2 years) and methylation context, DMRs were further characterized based on the directionality of differential methylation. Hypermethylated DMRs are those that show increased methylation in the oldest cohort in each contrast, and hypomethylated DMRs are those that show decreased methylation in the oldest cohort in each contrast. The cumulative binomial probability of identifying an equal or greater number of hypermethylated DMRs in each age contrast by methylation context was calculated using the R base package stats command *pbinom()* where x = the number of hypermethylated DMRs in each age contrast by methylation context, size = the total number of DMRs identified in each age contrast/methylation context, p = 0.5, and lower.tail = FALSE.

To visualize enrichment of DMRs across the eight chromosomes in the ‘Nonpareil’ genome, circos plots were generated with one track depicting each DMR classified as either hyper- or hypomethylated and two additional tracks depicting DMR enrichment across the genome. To create the circos plots, the R package circlize v. 0.41.2 (Gu *et al*., 2014) was used along with the bed files for all hyper- and hypomethylated DMRs in each methylation context for all age contrasts. The command *circos.genomicRainfall()* was used to create the first track with dots representing each individual DMR (red – hypermethylated, blue – hypomethylated) and positioned based on the number of DMRs occurring in that location. The command *circos.genomicDensity()* was used to create the two additional tracks representing enrichment of hyper- and hypomethylated DMRs on each chromosome where the taller the peak, the higher the number of DMRs occurring in the specific region (Gu *et al*., 2014).

Following classification into hyper- and hypomethylated regions, bed files were generated for these DMRs using genomic coordinates. These bed files were used as input along with the ‘Nonpareil’ genome annotation file into the R v. 4.0.2 (R Core Team, 2020) packages GenomicRanges v. 1.40.0 (Lawrence *et al*., 2013) and genomation v. 1.20.0 (Akalin *et al*., 2015) to prepare a GRanges object and annotate DMRs using the command *annotateWithFeatures()*.

Initial annotation of DMRs by features includes the percentage of DMRs overlapping one of four features: gene, exon, 5’ untranslated region (UTR), and 3’ UTR. The DMRs were further annotated, and gene ontology (GO) enrichment was performed using the software HOMER v. 4.11 (Heinz *et al*., 2010) and the R package topGO v. 2.40.0 (Alexa A, 2020). Initially, all DMRs were annotated by assigning the gene with the closest transcriptional start site to each DMR using *annotatePeaks.pl -noann* with the ‘Nonpareil’ genome and genome annotation files. This produced a list of gene identifiers from the genome annotation file that are associated with each DMR. The GO terms assigned to each DMR-associated gene were used as input along with the ‘Nonpareil’ genomic annotation file to determine enrichment of GO terms in each age contrast DMR-associated gene set. The DMR-associated genes in each methylation context were classified based on biological process, molecular function, and cellular component GO term to produce tables depicting the number of DMR-associated genes assigned to each descriptor.

The DMRs were then further classified based on the occurrence of overlapping genomic regions among DMRs when comparing age contrasts. The bed files generated above were used as input in the bedtools v. 2.29.2 (Quinlan & Hall, 2010) command *intersect -wao* to identify overlaps in DMRs from each of the age contrasts (i.e., 11-7 contrast compared to 11-2 contrast). Finally, genomic regions were identified that contain significant DMRs in all three age contrasts using bedtools *intersect*. A similar process was used to determine DMRs overlapping transposable elements between two age cohorts (2-year-old *vs.* 7-year-old) using the annotation file to determine the most likely transposable element using EDTA ver. 2.0.0 (Ou et al., 2019) The DMRs overlapping genes were annotated using the *annotatePeaks.pl* script as above to find DMR-associated genes as well as GO terms, Pfam identifiers, and Interpro identifiers associated with each gene. The genomic DMR sequence was extracted from the ‘Nonpareil’ genome fasta file, and individual DMR fasta files were searched against the miRbase database v. 22.1 (Kozomara *et al*., 2019) using blastn (Camacho *et al*., 2009) to identify any putative microRNAs (miRNAs) within those regions. Searches were performed using an e-value cutoff of 1 after filtering the blast database to include only the following genera: *Arabidopsis*, *Prunus*, *Malus*, *Glycine*, *Medicago*, *Populus*, and *Solanum*.

Hits were subsequently filtered based on length of the putative microRNA within each DMR sequence with a cutoff of n – 2 base pairs based on the length of the identified putative microRNA within the database. A further filtering step was performed based on the e-value of the hits returning only those microRNAs with the lowest e-value for each potential match to the DMR sequences. This process resulted in at least one microRNA representing the best hit for each putative microRNA sequence within each DMR. In some cases, more than one best hit is returned if the e-values are equal for more than one hit.

## Annotation of unknown protein sequences associated with DMRs

A total of eight unknown proteins were identified as associated with the DMRs shared between the three age contrasts. Several programs were used to annotate these protein sequences and determine additional information about their putative functions. The program ProtParam was used to characterize protein properties, including molecular weight (Gasteiger *et al*., 2005). To predict subcellular localization, the program YLoc was used (Briesemeister *et al*., 2010a,b).

Finally, the Motif tool on the GenomeNet website (https://www.genome.jp/tools/motif/) was used to search a protein query against several databases to identify putative alignments (Marchler-Bauer *et al*., 2013; Sigrist *et al*., 2013; Finn *et al*., 2014).

## Small RNA-Seq Differential Expression Analysis

Following sequencing, Trimmomatic (Bolger, 2014) was run in single end mode to remove adaptor sequences and trim low-quality bases from the ends of raw reads. The expected adaptor sequence was ‘*AGATCGGAAGAGCACACGTCTGAACTCCAGTCAC.*’ Values for the seed mismatch, palindrome clip threshold, and simple clip threshold parameters within the ILLUMINACLIP argument were set to 2, 30, and 6, respectively. Other options passed to Trimmomatic included LEADING:3, TRAILING:3, SLIDINGWINDOW:6:20, and MINLEN:12. Trimmed reads were aligned to the ‘*Prunus dulcis* Nonpareil’ reference genome (D’Amico-Willman *et al*., 2022) with Shortstack v3.8.5 (Johnson *et al*., 2016) using default parameters. All aligned small RNA sequences were obtained and used as input for a Blast analysis.

A custom Blast database was generated by downloading the mature miRNA database from miRbase and filtering it for entries associated with the genera *Prunus*, *Arabidopsis*, *Malus*, *Glycine*, *Medicago*, *Populus*, or *Solanum*. The aligned small RNA sequences were searched against this custom database using the blastn function within Blast 2.10.0+ (Camacho, 2009) with the following settings: -evalue: 1, -task: blasn-short, -word_size: 4, -gapopen: 25, -gapextend: 10, -reward: 5, -penalty: 4, -dust: no, and -soft_masking: false. The initial set of Blast hits was filtered to retain only those that had an aligned length no less than (mature miR length – 2). For each unique query sequence, the hit with the best (lowest) e-value was retained, and results were categorized into miRNA families.

The counts table obtained from ShortStack was used as input to DESeq2 (Love *et al*., 2014) to test each aligned small RNA for evidence of differential expression between the two age groups. Significant differentially expressed genes were defined as those with an absolute log2 fold change >2 and an adjusted p-value <0.01.

---

**Figure S1.**
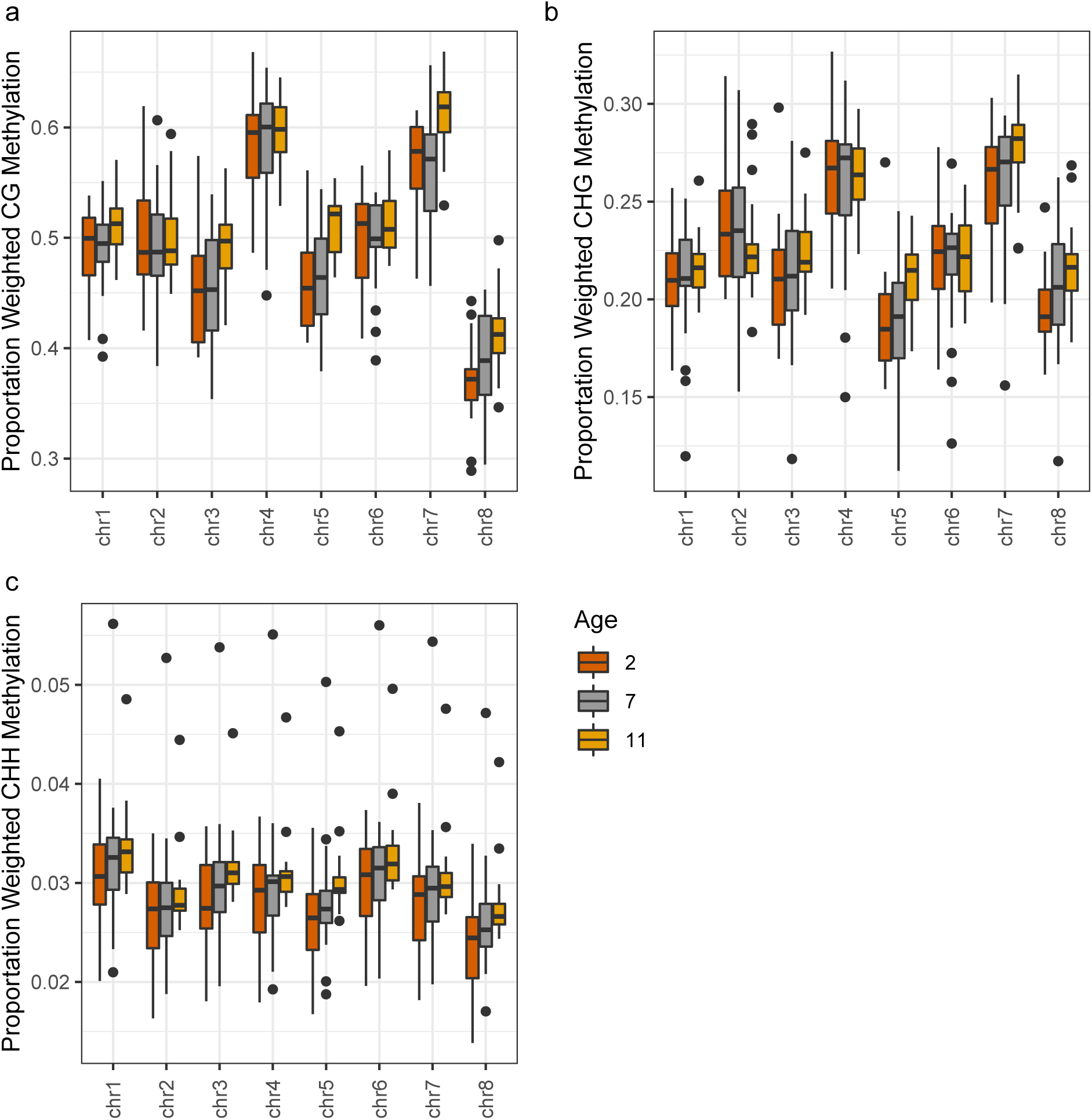
Boxplots depicting the proportion of weighted methylation in each age cohort (2 years old – red; 7 years old – grey; 11 years old – yellow) across the three methylation contexts: **(a)** CG, **(b)** CHG, and **(c)** CHH. The black dots represent outliers.

**Figure S2a-l.**
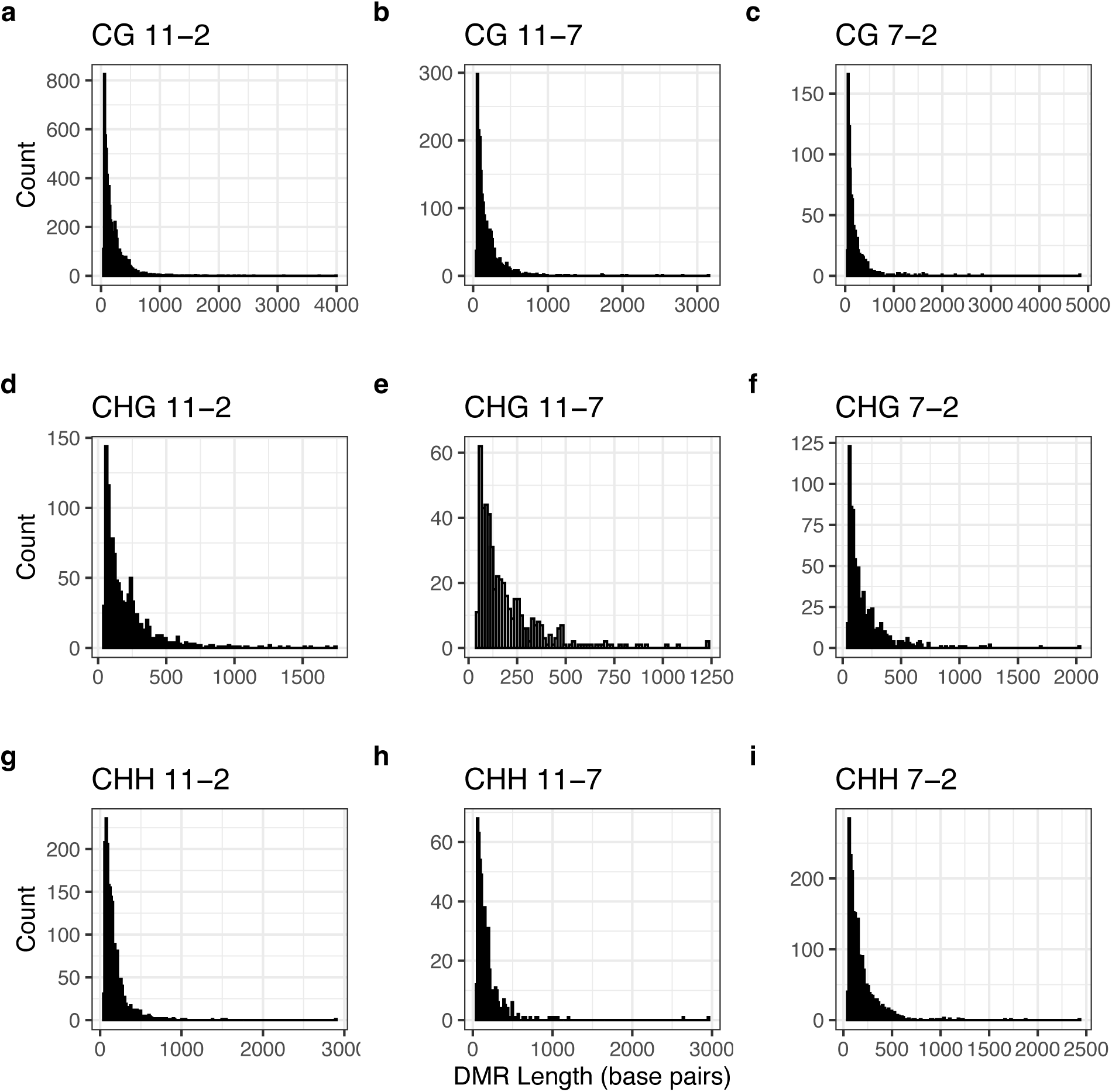
Distribution of lengths in base pairs of differentially methylated regions (DMRs) identified in each age contrast and methylation context. Panels **a-c** show distribution of DMRs identified in the CG context, panels **d-f** show distribution in the CHG context, and panels **g-l** show distribution in the CHH context. The values listed next to the methylation context indicate the age-contrast (11 – 2 year, 11 – 7 year, and 7 – 2 year).

**Figure S3a-c.**
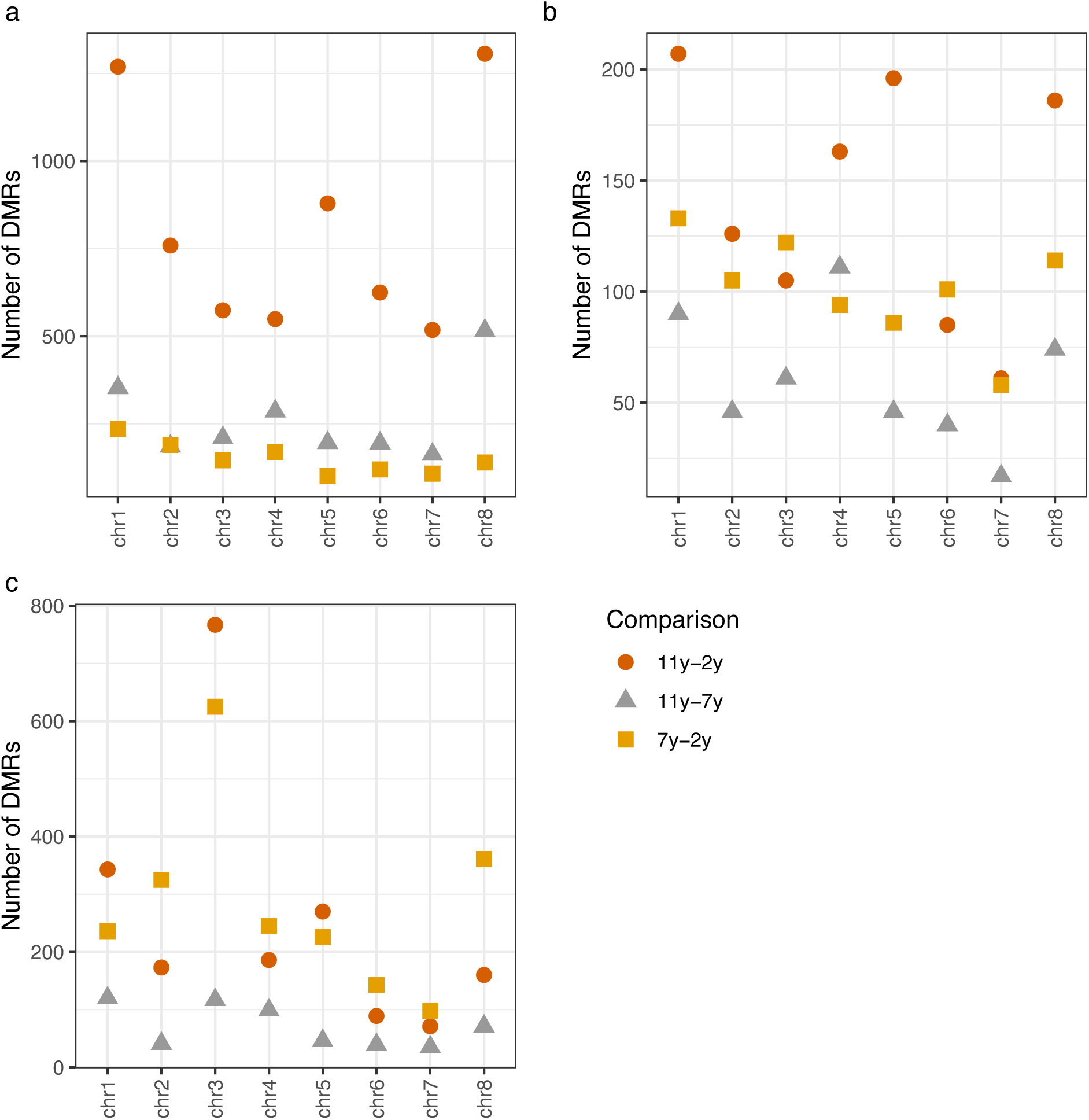
The dot plots represent the number of significant (p < 0.0001) differentially methylated regions (DMRs) identified in each of the contrasts (11 – 2 years: red; 11 – 7 years: grey; 7 – 2 years: yellow) in each methylation-context: **(a)** CG, **(b)** CHG, and **(c)** CHH.

**Figure S4a-m.**
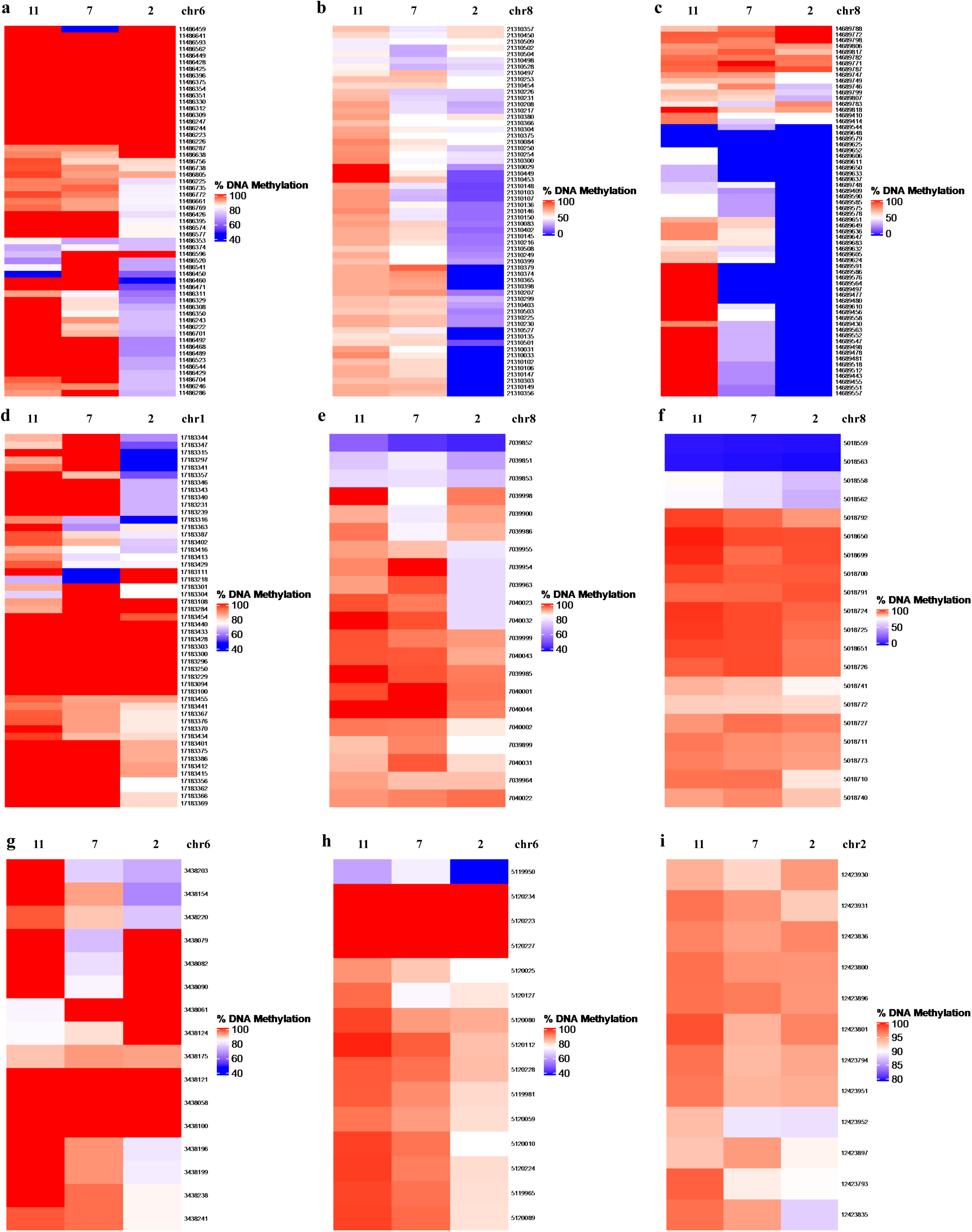
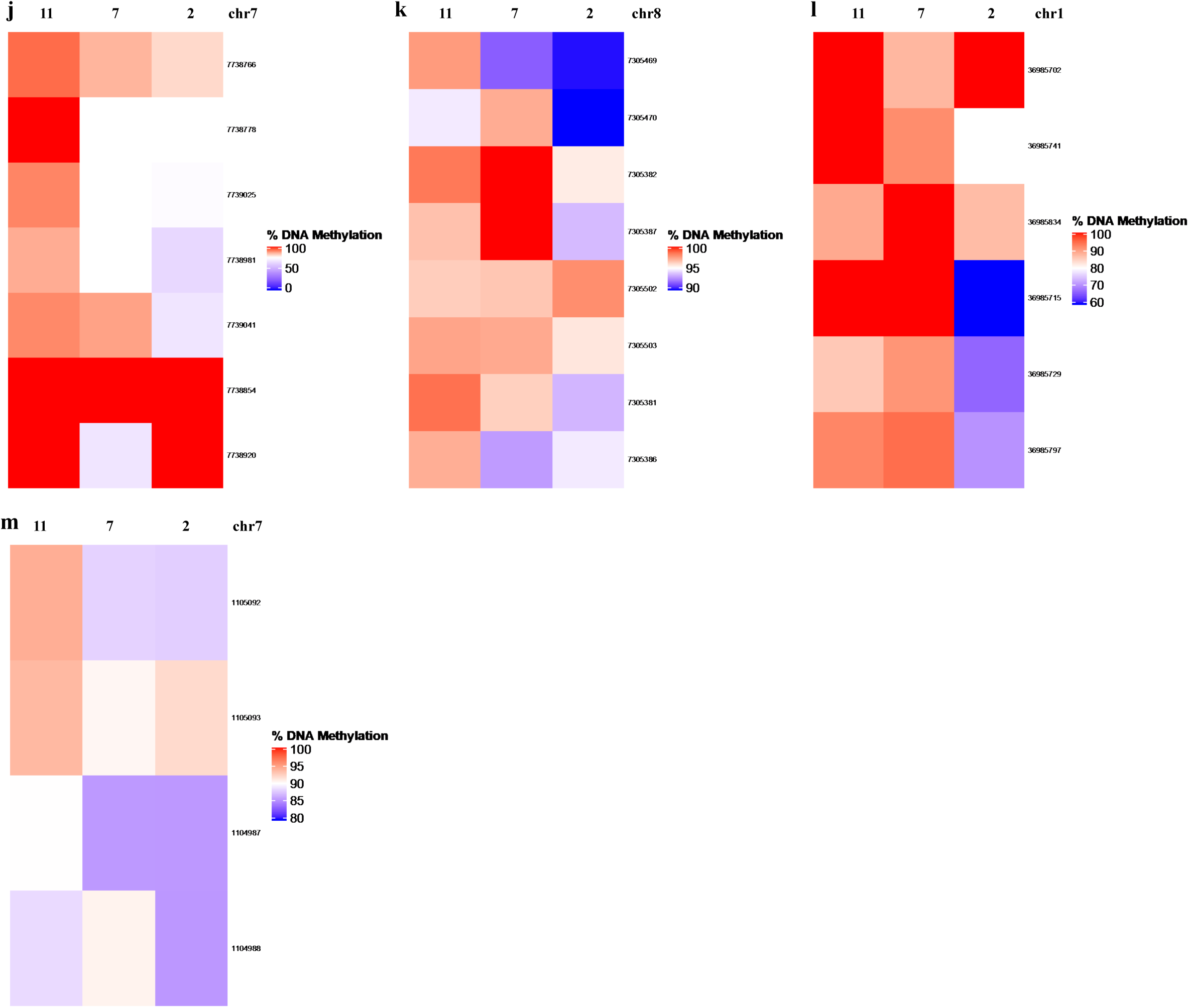
Heatmaps displaying average percent DNA methylation across cytosines in the 11-year, 7-year, and 2-year age cohorts within the genomic range of 13 overlapping differentially methylated regions (DMRs) in the CG context identified in the three age-contrasts. The regions correspond to CGDMR1-13 **(a-m; see Table S2),** and the values to the right of each heatmap represent the genomic position of each cytosine on the respective chromosome.

**Figure S5a-c.**
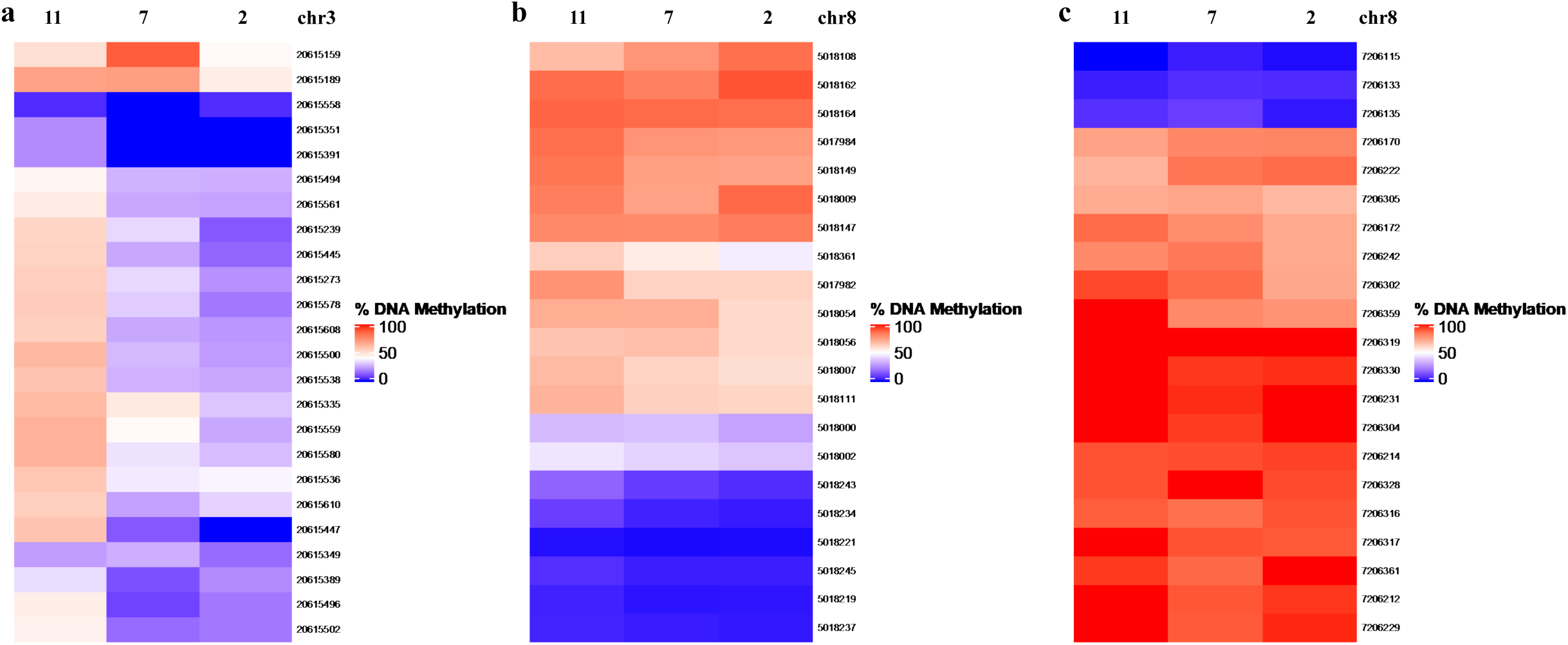
Heatmaps displaying average percent DNA methylation across cytosines in the 11-year, 7-year, and 2-year age cohorts within the genomic range of 3 overlapping differentially methylated regions (DMRs) in the CHG context identified in the three age-contrasts. The regions correspond to CHGDMR1-3 **(a-c; see Table S2),** and the values to the right of each heatmap represent the genomic position of each cytosine on the respective chromosome.

**Figure S6.**
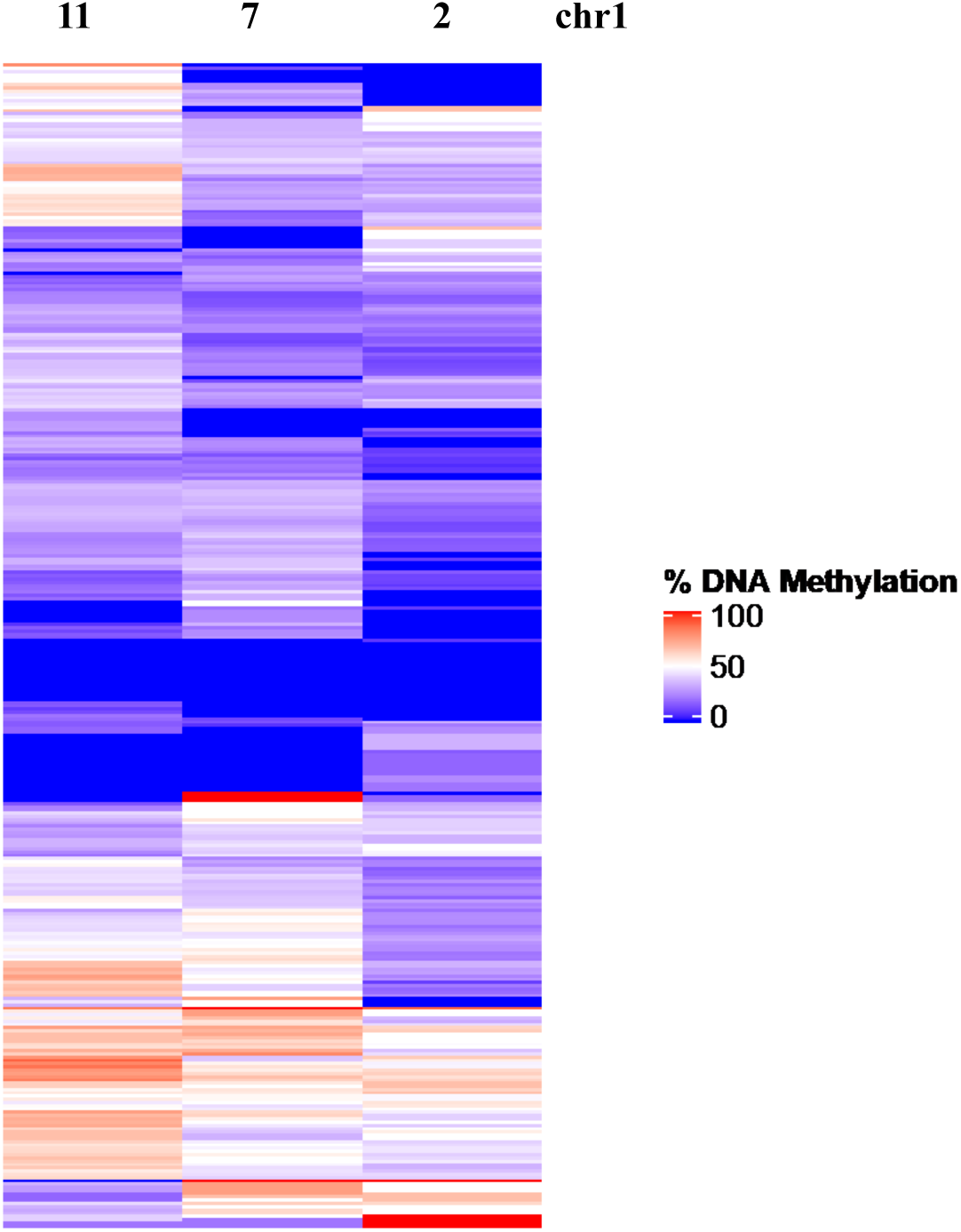
Heatmap displaying average percent DNA methylation across cytosines in the 11-year, 7-year, and 2-year age cohorts within the genomic range of the overlapping differentially methylated region (DMR) in the CHH context identified in the three age-contrasts. The regions correspond to CHHDMR1 **(see Table S2)**.

**Figure S7.**
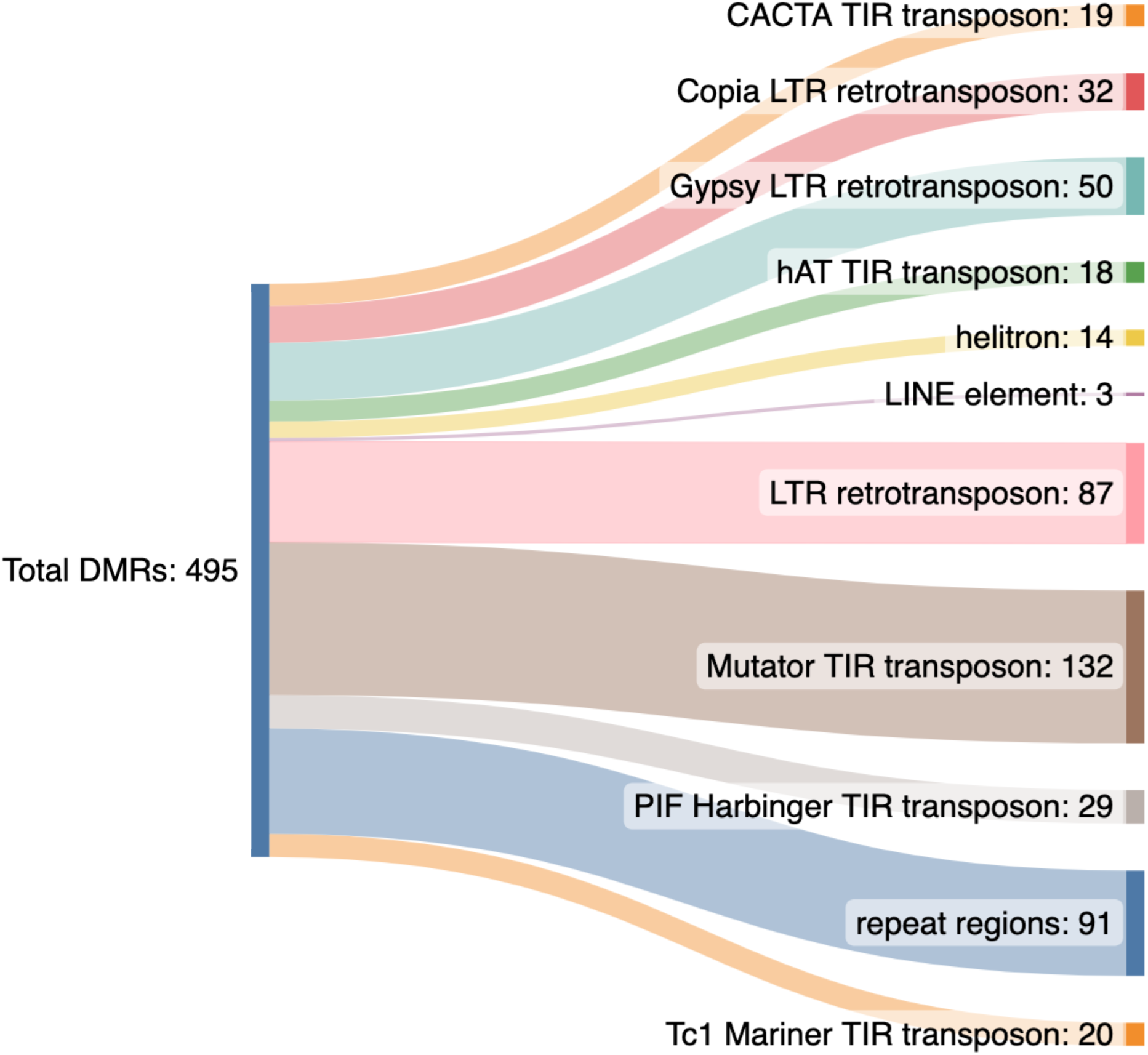
Distribution by type of transposable elements overlapping with 493 differentially methylated regions (DMRs) identified in the 2 – 7-year-old age contrast.

**Table S1.** Pairwise comparison of least squared means of weighted percent methylation in the CG, CHG, and CHH contexts for each chromosome in the ‘Nonpareil’ almond genome. Age cohort contrasts include the 2 – 11, the 7 – 11, and the 2 – 7-year contrasts. Significant contrasts are represented in bold with an alpha = 0.05.

| Chromosome | Age-Contrast | CG context<br>p-value | CHG context<br>p-value | CHH context<br>p-value |
| --- | --- | --- | --- | --- |
| Chr1 | 11-2 | **0.0235** | 0.5329 | 0.0595 |
|  | 11-7 | **0.0422** | 0.4425 | 0.7068 |
|  | 7-2 | 0.9436 | 0.9949 | 0.2780 |
| Chr2 | 11-2 | 0.9973 | 0.3587 | 0.1320 |
|  | 11-7 | 0.7356 | 0.9063 | 0.6008 |
|  | 7-2 | 0.7924 | 0.5948 | 0.5698 |
| Chr3 | 11-2 | **0.0020** | 0.2137 | **0.0138** |
|  | 11-7 | **0.0046** | 0.1647 | 0.3543 |
|  | 7-2 | 0.9267 | 0.9980 | 0.2883 |
| Chr4 | 11-2 | 0.6659 | 0.9917 | 0.0972 |
|  | 11-7 | 0.5390 | 0.8626 | 0.6030 |
|  | 7-2 | 0.9867 | 0.8071 | 0.4789 |
| Chr5 | 11-2 | **<.0001** | **0.0056** | **0.0020** |
|  | 11-7 | **<.0001** | **0.0049** | 0.1935 |
|  | 7-2 | 0.8706 | 0.9954 | 0.1873 |
| Chr6 | 11-2 | 0.3472 | 0.9922 | 0.0675 |
|  | 11-7 | 0.1645 | 0.7924 | 0.5824 |
|  | 7-2 | 0.9342 | 0.8679 | 0.4042 |
| Chr7 | 11-2 | **0.0003** | 0.0658 | 0.0956 |
|  | 11-7 | **0.0002** | **0.0263** | 0.7183 |
|  | 7-2 | 0.9963 | 0.9676 | 0.3716 |
| Chr8 | 11-2 | **0.0004** | **0.0163** | **0.0055** |
|  | 11-7 | **0.0295** | 0.1837 | 0.4246 |
|  | 7-2 | 0.3440 | 0.5279 | 0.1329 |

**Table S2.** Genomic coordinates and length in base pairs for the 17 overlapping DMRs occurring in each of the three age-contrasts.

| DMR_ID | Chr | 11-2 |  |  | 11-7 |  |  | 7-2 |  |  |
| --- | --- | --- | --- | --- | --- | --- | --- | --- | --- | --- |
|  |  | Start | End | Length (bp) | Start | End | Length (bp) | Start | End | Length (bp) |
| CGDMR1 | 6 | 11486222 | 11486807 | 585 | 11486637 | 11486807 | 170 | 11486637 | 11486807 | 170 |
| CGDMR2 | 8 | 21310028 | 21310528 | 500 | 21310028 | 21310149 | 121 | 21310028 | 21310403 | 375 |
| CGDMR3 | 8 | 14689409 | 14689818 | 409 | 14689430 | 14689807 | 377 | 14689477 | 14689547 | 70 |
| CGDMR4 | 1 | 17183093 | 17183455 | 362 | 17183249 | 17183343 | 94 | 17183249 | 17183315 | 66 |
| CGDMR5 | 8 | 7039851 | 7040044 | 193 | 7039954 | 7040044 | 90 | 7039899 | 7040044 | 145 |
| CGDMR6 | 8 | 5018558 | 5018792 | 234 | 5018650 | 5018792 | 142 | 5018740 | 5018792 | 52 |
| CGDMR7 | 6 | 3438057 | 3438262 | 205 | 3438026 | 3438089 | 63 | 3438057 | 3438174 | 117 |
| CGDMR8 | 6 | 5119949 | 5120234 | 285 | 5119949 | 5120126 | 177 | 5119949 | 5120126 | 177 |
| CGDMR9 | 2 | 12423793 | 12423952 | 159 | 12423835 | 12423931 | 96 | 12423896 | 12423952 | 56 |
| CGDMR10 | 7 | 7738726 | 7739046 | 320 | 7738980 | 7739046 | 66 | 7738980 | 7739046 | 66 |
| CGDMR11 | 8 | 7305381 | 7305503 | 122 | 7305386 | 7305503 | 117 | 7305381 | 7305503 | 122 |
| CGDMR12 | 1 | 36985701 | 36985875 | 174 | 36985796 | 36985875 | 79 | 36985796 | 36985875 | 79 |
| CGDMR13 | 1 | 1104987 | 1105093 | 106 | 1104987 | 1105093 | 106 | 1104987 | 1105093 | 106 |
| CHGDMR1 | 3 | 20615159 | 20615610 | 451 | 20615337 | 20615502 | 165 | 20615337 | 20615391 | 54 |
| CHGDMR2 | 8 | 5017982 | 5018361 | 379 | 5018000 | 5018056 | 56 | 5018000 | 5018056 | 56 |
| CHGDMR3 | 8 | 7206115 | 7206361 | 246 | 7206170 | 7206242 | 72 | 7206170 | 7206231 | 61 |
| CHHDMR1 | 1 | 34468252 | 34469756 | 1504 | 34469043 | 34469756 | 713 | 34468083 | 34469120 | 1037 |

**Table S3.** Annotation of hyper- and hypomethylated differentially methylated regions (DMRs) in each methylation context and for each age-contrast. The ‘Nonpareil’ genome annotation was used to classify the DMRs into four categories: gene, exon, five prime untranslated region (5’ UTR), and three prime untranslated region (3’ UTR). The percentages under each classification represent the percentage of DMRs from each methylation-context and contrast in each of the four categories.

| **Methylation Context** | **Hyper/Hypo** | **Contrast** | **%Gene** | **%Exon** | **%5’ UTR** | **%3’ UTR** |
| --- | --- | --- | --- | --- | --- | --- |
| CG | Hyper | 11 – 2 | 28.0 | 20.6 | 0.91 | 0.96 |
|  |  | 11 – 7 | 34.2 | 25.1 | 1.51 | 1.46 |
|  |  | 7 – 2 | 27.8 | 21.4 | 0.81 | 1.32 |
|  | Hypo | 11 – 2 | 39.7 | 31.2 | 7.29 | 1.21 |
|  |  | 11 – 7 | 22.1 | 14.8 | 0.82 | 3.28 |
|  |  | 7 – 2 | 29.2 | 22.1 | 2.65 | 0.88 |
| CHG | Hyper | 11 – 2 | 27.7 | 20.1 | 2.49 | 1.70 |
|  |  | 11 – 7 | 24.6 | 18.3 | 3.17 | 1.98 |
|  |  | 7 – 2 | 24.8 | 17.2 | 1.91 | 0.76 |
|  | Hypo | 11 – 2 | 23.5 | 17.8 | 3.28 | 1.37 |
|  |  | 11 – 7 | 27.9 | 18.5 | 3.00 | 2.58 |
|  |  | 7 – 2 | 19.4 | 13.8 | 2.77 | 0 |
| CHH | Hyper | 11 – 2 | 18.0 | 9.41 | 1.60 | 2.13 |
|  |  | 11 – 7 | 19.0 | 12.0 | 0.46 | 1.39 |
|  |  | 7 – 2 | 17.9 | 9.20 | 1.35 | 2.16 |
|  | Hypo | 11 – 2 | 27.4 | 14.6 | 3.25 | 1.63 |
|  |  | 11 – 7 | 24.5 | 14.0 | 3.99 | 1.99 |
|  |  | 7 – 2 | 24.4 | 14.2 | 1.22 | 1.22 |

**Table S4a-c.** Number of CG hypermethylated DMR-associated genes associated with each biological process (**A)**, molecular function **(B)**, and cellular component **(C)** gene ontology (GO) terms for each of the three age-contrasts (i.e., 11 – 2 year, 11 – 7 year, and 7 – 2 year). Values in each column represent the number of DMR-associated genes that are associated with each GO term. Black squares indicate no genes associated with that contrast were assigned the particular GO term.

| **A. Biological Process GO Term** | **11-2** | **11-7** | **7-2** |
| --- | --- | --- | --- |
| proteolysis | 37 | 36 | 8 |
| protein phosphorylation | 29 | 28 | 7 |
| obsolete oxidation-reduction process | 28 | 22 | 4 |
| transmembrane transport | 18 | 16 | 4 |
| regulation of transcription, DNA-templated | 9 | 13 | 2 |
| carbohydrate metabolic process | 10 | 11 | 4 |
| steroid biosynthetic process | 5 |  |  |
| tRNA aminoacylation for protein translation | 3 |  |  |
| signal transduction | 1 | 8 |  |
| translation | 2 | 8 |  |
| biosynthetic process | 1 | 7 |  |
| transcription, DNA-templated | 1 | 7 | 1 |
| DNA topological change |  | 6 |  |
| cation transport | 2 | 4 |  |
| endoplasmic reticulum to Golgi vesicle-mediated transport |  | 3 |  |
| nitrogen compound metabolic process |  | 3 | 1 |
| proteolysis involved in cellular protein catabolic process |  | 3 |  |
| tRNA processing | 3 | 3 |  |
| ATP synthesis coupled proton transport |  | 2 |  |
| attachment of GPI anchor to protein |  | 2 |  |
| cell redox homeostasis |  | 2 |  |
| cell wall modification |  | 2 |  |
| defense response |  | 2 |  |
| fatty acid biosynthetic process |  | 2 |  |
| histone lysine methylation | 2 |  |  |
| lipid metabolic process | 1 | 2 |  |
| metal ion transport |  | 2 |  |
| nucleotide-excision repair | 2 |  |  |
| proteasome-mediated ubiquitin-dependent protein catabolic process | 2 |  |  |
| recognition of pollen |  | 2 |  |
| translational initiation |  | 2 |  |
| cell wall macromolecule catabolic process |  | 1 |  |
| cellular glucan metabolic process |  | 1 |  |
| cellular modified amino acid biosynthetic process | 1 | 1 |  |
| cellulose biosynthetic process | 1 |  | 1 |
| chitin catabolic process |  | 1 |  |
| dephosphorylation |  | 1 |  |
| DNA replication |  | 1 |  |
| DNA-templated transcription, termination | 1 |  |  |
| drug transmembrane transport |  | 1 |  |
| endoplasmic reticulum organization |  | 1 |  |
| fatty acid beta-oxidation | 2 | 1 |  |
| glutamine biosynthetic process |  | 1 |  |
| intermembrane lipid transfer |  | 1 |  |
| intracellular protein transport |  | 1 |  |
| intracellular transport |  | 1 |  |
| ion transport |  | 1 |  |
| malate transport | 1 |  |  |
| mitochondrial pyruvate transmembrane transport |  | 1 |  |
| mRNA processing |  | 1 |  |
| nucleobase-containing compound metabolic process |  | 1 |  |
| nucleotide transmembrane transport |  | 1 |  |
| nucleotide transport |  |  | 1 |
| nucleosome assembly |  | 1 |  |
| phosphatidylserine biosynthetic process |  | 1 |  |
| photosynthesis | 1 |  |  |
| photosynthesis, light reaction |  | 1 |  |
| photosynthetic electron transport in photosystem II |  | 1 |  |
| phytochromobilin biosynthetic process |  | 1 |  |
| potassium ion transport |  | 1 |  |
| protein folding | 2 | 1 |  |
| protein kinase C-activating G protein-coupled receptor signaling pathway | 1 |  |  |
| protein neddylation |  | 1 |  |
| protein ubiquitination | 1 | 1 |  |
| proton transmembrane transport | 1 |  |  |
| pseudouridine synthesis |  |  | 1 |
| pyrimidine nucleotide-sugar transmembrane transport |  | 1 |  |
| regulation of translational fidelity | 1 |  |  |
| response to auxin |  |  | 1 |
| response to hormone |  | 1 |  |
| RNA modification |  |  | 1 |
| RNA processing |  | 1 |  |
| rRNA processing |  | 1 |  |
| sucrose metabolic process |  | 1 | 1 |
| translation termination |  |  | 1 |
| terpenoid biosynthetic process | 1 |  |  |
| trehalose biosynthetic process | 1 |  |  |
| tRNA wobble uridine modification |  | 1 |  |

| **B. Molecular Function GO Term** | **11-2** | **11-7** | **7-2** |
| --- | --- | --- | --- |
| ATP binding | 41 | 33 | 11 |
| cysteine-type peptidase activity | 36 | 32 | 8 |
| hydrolase activity, acting on ester bonds | 26 | 28 | 4 |
| protein binding | 35 | 27 | 10 |
| protein kinase activity | 18 | 26 | 5 |
| DNA binding | 13 | 20 | 5 |
| zinc ion binding | 20 | 20 | 7 |
| ADP binding | 3 | 18 |  |
| nucleic acid binding | 18 | 16 | 3 |
| oxidoreductase activity | 3 | 13 | 1 |
| transferase activity, transferring hexosyl groups |  | 11 | 3 |
| catalytic activity | 4 | 9 |  |
| DNA-binding transcription factor activity | 7 | 8 | 1 |
| structural constituent of ribosome | 2 | 8 |  |
| DNA topoisomerase activity |  | 6 |  |
| transmembrane transporter activity | 3 | 6 | 1 |
| hydrolase activity, hydrolyzing O-glycosyl compounds | 6 | 5 | 2 |
| polygalacturonase activity | 4 | 5 | 2 |
| RNA binding | 1 | 5 | 1 |
| sequence-specific DNA binding | 1 | 5 |  |
| transferase activity, transferring acyl groups other than amino-acyl groups | 3 | 5 | 1 |
| acetylglucosaminyltransferase activity | 1 | 4 |  |
| aminoacyl-tRNA ligase activity | 3 |  |  |
| DNA-directed 5'-3' RNA polymerase activity | 1 | 4 | 1 |
| heme binding | 10 | 4 |  |
| iron ion binding | 10 | 4 |  |
| lyase activity | 4 | 4 |  |
| magnesium ion binding | 4 | 4 |  |
| nucleotide binding | 3 |  |  |
| oxidoreductase activity, acting on paired donors, with incorporation or reduction of molecular oxygen | 9 | 4 |  |
| phosphatidylinositol binding |  |  | 4 |
| protein dimerization activity | 4 | 4 | 1 |
| RNA-DNA hybrid ribonuclease activity | 10 | 4 | 2 |
| solute:proton antiporter activity | 1 | 4 |  |
| terpene synthase activity | 4 | 4 |  |
| GTP binding | 4 | 3 | 1 |
| GTPase activity |  | 3 | 1 |
| hydrolase activity | 1 | 3 |  |
| oxidoreductase activity, acting on the CH-OH group of donors, NAD or NADP as acceptor | 5 | 3 |  |
| threonine-type endopeptidase activity |  | 3 |  |
| 3-beta-hydroxy-delta5-steroid dehydrogenase activity | 5 |  |  |
| ATPase activity | 7 | 2 | 1 |
| calcium ion binding | 1 | 2 |  |
| copper ion binding | 1 | 2 |  |
| damaged DNA binding | 2 |  |  |
| electron transfer activity |  | 2 |  |
| enzyme inhibitor activity | 2 | 2 |  |
| histone-lysine N-methyltransferase activity | 2 |  |  |
| metal ion binding |  | 2 |  |
| methyltransferase activity | 2 | 2 |  |
| nickel cation binding |  | 2 |  |
| nutrient reservoir activity | 2 |  | 1 |
| O-methyltransferase activity | 1 | 2 |  |
| obsolete coenzyme binding | 2 | 2 |  |
| oxidoreductase activity, acting on NAD(P)H, oxygen as acceptor | 1 | 2 |  |
| pectinesterase activity |  | 2 |  |
| peroxidase activity | 2 | 2 |  |
| protein disulfide oxidoreductase activity |  | 2 |  |
| protein serine/threonine kinase activity | 11 | 2 | 2 |
| proton transmembrane transporter activity |  | 2 |  |
| serine-type carboxypeptidase activity |  | 2 |  |
| serine-type endopeptidase activity | 2 | 2 |  |
| transcription factor binding | 2 |  |  |
| transferase activity, transferring acyl groups | 2 |  | 1 |
| translation initiation factor activity |  | 2 |  |
| 2-C-methyl-D-erythritol 2,4-cyclodiphosphate synthase activity | 1 |  |  |
| 3'-5' exonuclease activity |  | 1 |  |
| acetyl-CoA carboxylase activity |  | 1 |  |
| acyl-CoA oxidase activity | 2 | 1 |  |
| antioxidant activity |  | 1 |  |
| antiporter activity | 1 | 1 |  |
| ATPase-coupled transmembrane transporter activity | 5 | 1 | 1 |
| ATP:ADP antiporter activity |  |  | 1 |
| cation transmembrane transporter activity | 1 |  |  |
| cellulose synthase (UDP-forming) activity | 1 |  | 1 |
| channel activity | 1 | 1 |  |
| chitin binding |  | 1 |  |
| chitinase activity |  | 1 |  |
| cobalt ion binding |  | 1 |  |
| diacylglycerol kinase activity | 1 |  |  |
| DNA-directed DNA polymerase activity |  | 1 |  |
| electron transporter, transferring electrons within the cyclic electron transport pathway of photosynthesis activity |  | 1 |  |
| endonuclease activity |  | 1 |  |
| endopeptidase inhibitor activity | 1 | 1 |  |
| FAD binding | 1 |  |  |
| glutamate-ammonia ligase activity |  | 1 |  |
| glutamate-cysteine ligase activity | 1 | 1 |  |
| glycerol-3-phosphate O-acyltransferase activity | 1 |  |  |
| GTPase activator activity | 1 | 1 |  |
| helicase activity |  | 1 |  |
| inorganic diphosphatase activity | 1 |  |  |
| intramolecular lyase activity |  | 1 |  |
| intramolecular transferase activity, phosphotransferases |  | 1 |  |
| ion channel activity |  | 1 |  |
| lipid binding |  | 1 |  |
| lipid transfer activity |  | 1 |  |
| iron-sulfur cluster binding | 1 |  |  |
| malate dehydrogenase (decarboxylating) (NAD<sup>+</sup>) activity |  | 1 |  |
| metal ion transmembrane transporter activity |  | 1 |  |
| metallopeptidase activity |  | 1 |  |
| NAD binding |  | 1 |  |
| NEDD8 activating enzyme activity |  | 1 |  |
| nucleoside transmembrane transporter activity |  | 1 |  |
| oxidoreductase activity, acting on single donors with incorporation of molecular oxygen, incorporation of two atoms of oxygen |  | 1 |  |
| oxidoreductase activity, acting on the CH-CH group of donors | 2 | 1 |  |
| oxidoreductase activity, acting on the CH-CH group of donors, iron-sulfur protein as acceptor |  | 1 |  |
| peptide-methionine (S)-S-oxide reductase activity | 4 | 1 |  |
| peptidyl-prolyl cis-trans isomerase activity |  | 1 |  |
| phospho-N-acetylmuramoyl-pentapeptide-transferase activity |  | 1 |  |
| polysaccharide binding |  | 1 |  |
| protein tyrosine/serine/threonine phosphatase activity |  | 1 |  |
| pseudouridine synthase activity |  |  | 1 |
| pyridoxal phosphate binding | 1 | 1 |  |
| pyrimidine nucleotide-sugar transmembrane transporter activity |  | 1 |  |
| pyrophosphate hydrolysis-driven proton transmembrane transporter activity | 1 |  |  |
| racemase activity, acting on amino acids and derivatives |  |  | 1 |
| strictosidine synthase activity |  | 1 |  |
| sucrose synthase activity |  | 1 | 1 |
| translation release factor activity |  |  | 1 |
| ubiquitin-like modifier activating enzyme activity |  | 1 |  |
| ubiquitin-protein transferase activity | 1 | 1 |  |
| unfolded protein binding | 2 | 1 |  |
| uridylyltransferase activity |  | 1 |  |
| xyloglucan:xyloglucosyl transferase activity |  | 1 |  |

| **C. Cellular Component GO Term** | **11-2** | **11-7** | **7-2** |
| --- | --- | --- | --- |
| integral component of membrane | 16 | 18 | 4 |
| membrane | 10 | 16 | 4 |
| nucleus | 5 | 9 | 2 |
| ribosome | 2 | 8 |  |
| endoplasmic reticulum | 2 |  |  |
| endoplasmic reticulum membrane |  | 4 |  |
| proteasome core complex |  | 3 |  |
| cytoplasm |  | 2 |  |
| GPI-anchor transamidase complex |  | 2 |  |
| mitochondrial proton-transporting ATP synthase complex, coupling factor F(o) |  | 2 |  |
| nucleolus | 2 |  |  |
| nucleosome |  | 2 |  |
| acetyl-CoA carboxylase complex |  | 1 |  |
| apoplast |  | 1 |  |
| cell wall |  | 1 |  |
| chromosome |  | 1 |  |
| elongator holoenzyme complex |  | 1 |  |
| Golgi membrane |  | 1 |  |
| Las1 complex |  | 1 |  |
| mitochondrial inner membrane |  | 1 |  |
| nuclear pore |  | 1 |  |
| peroxisome | 2 | 1 |  |
| photosystem I | 1 |  |  |
| photosystem I reaction center | 1 |  |  |

**Table S5a-c.**
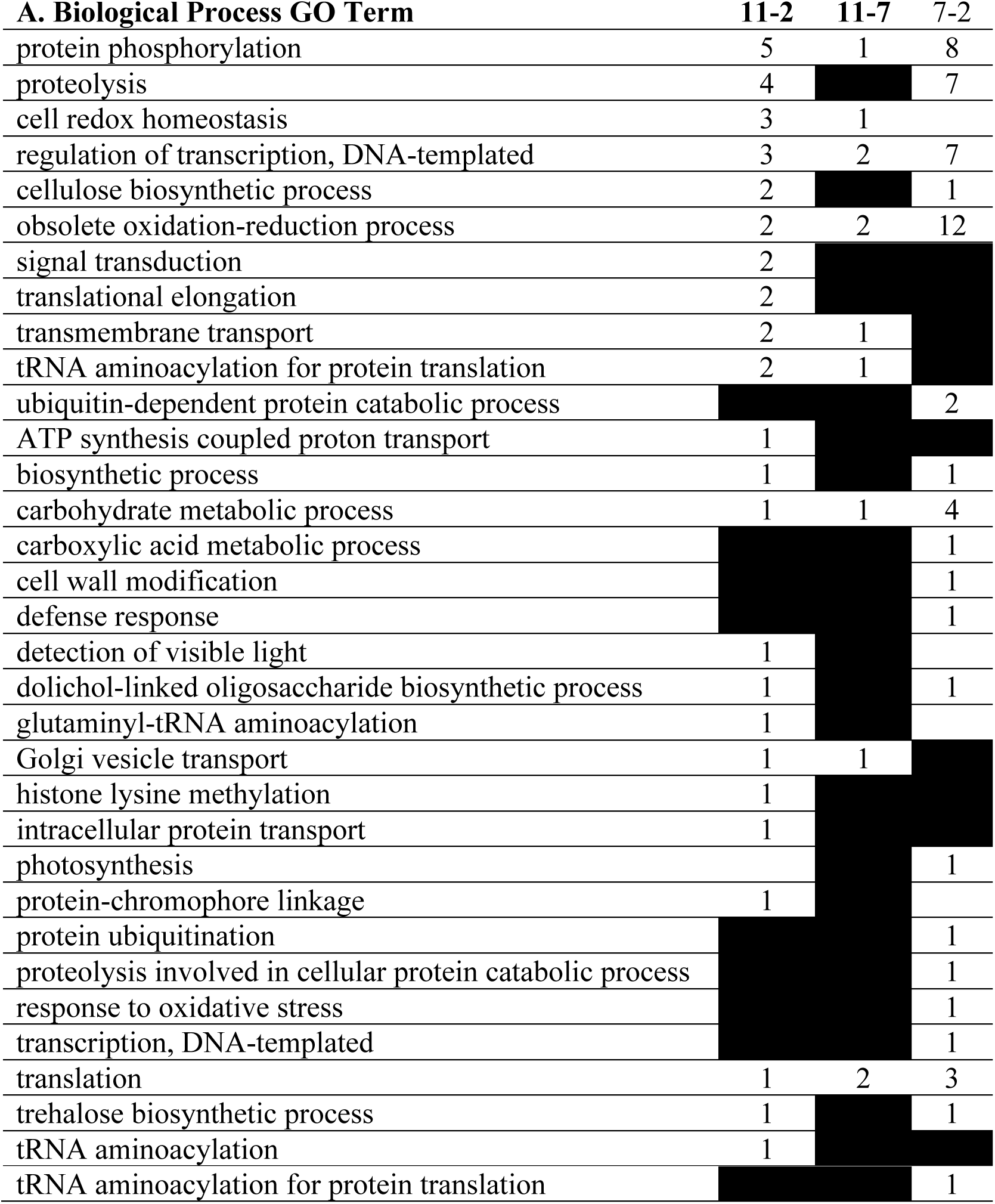
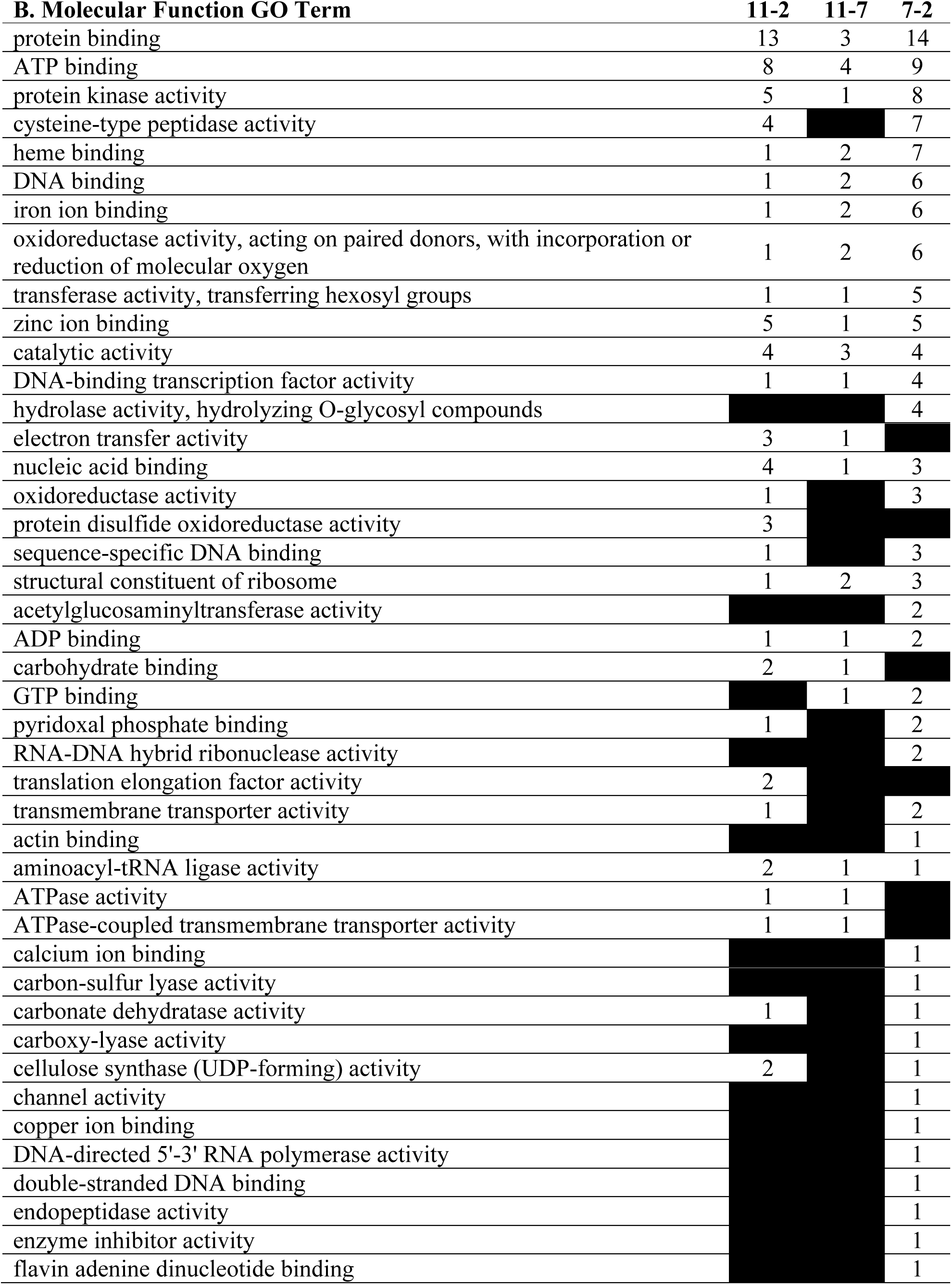
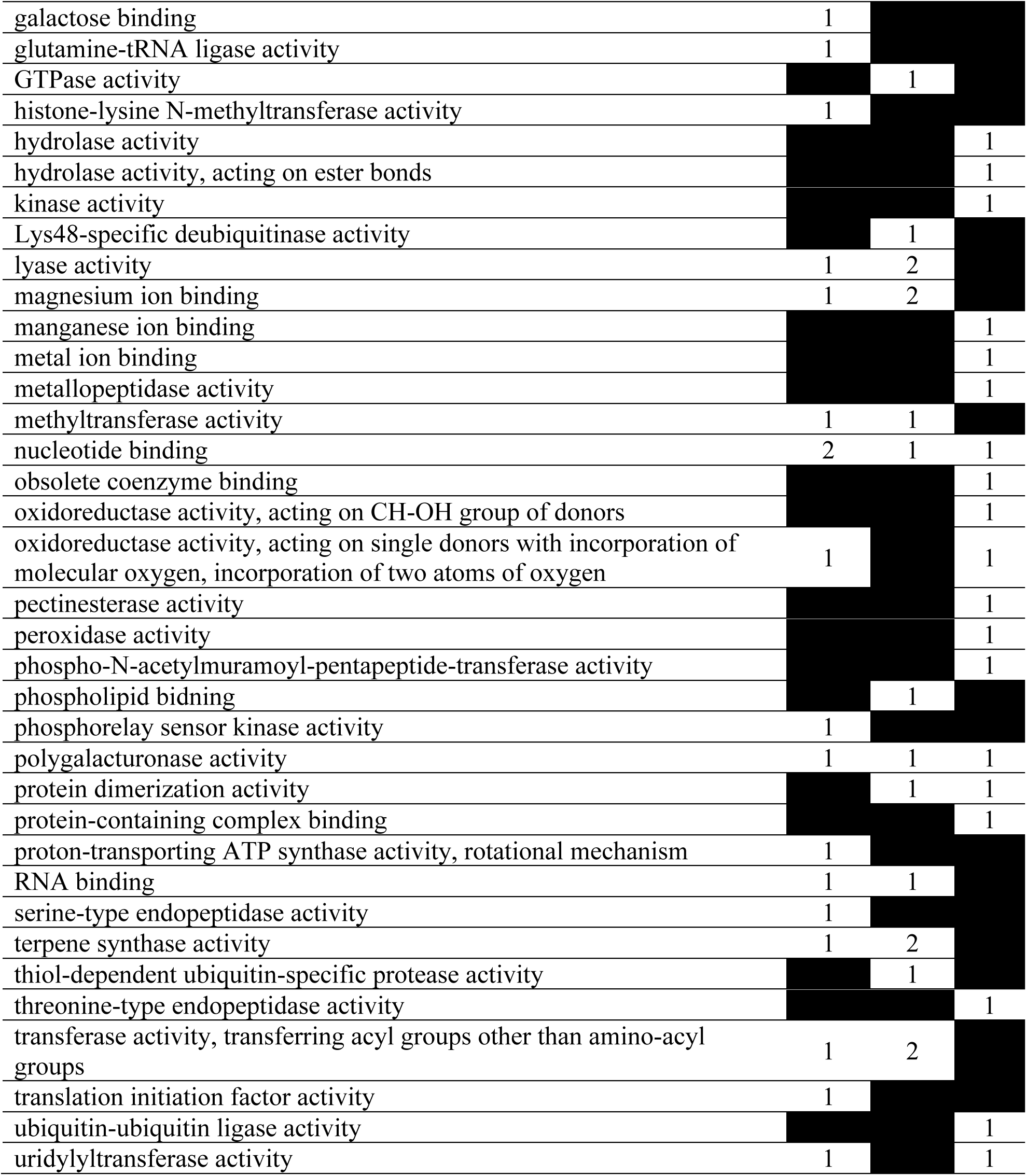
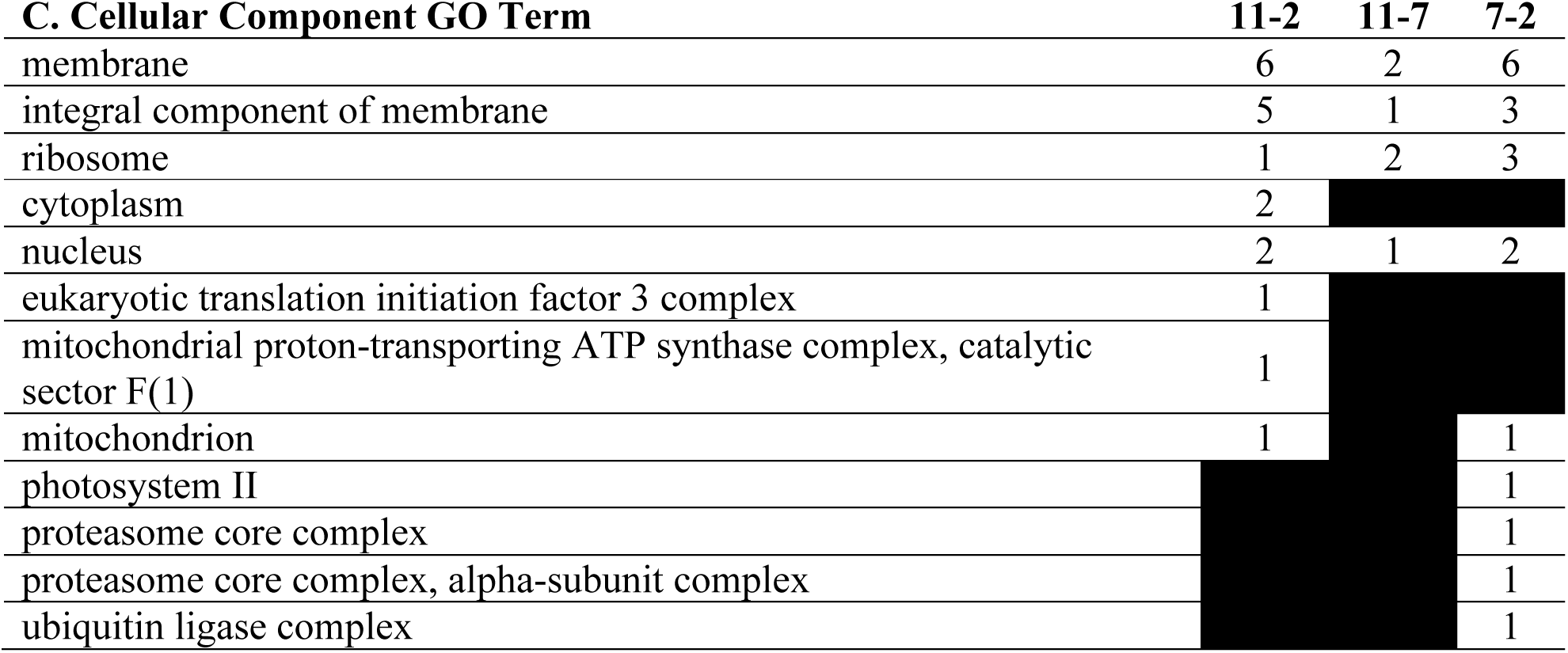
Number of CG hypomethylated DMR-associated genes associated with each biological process (**A)**, molecular function **(B)**, and cellular component **(C)** gene ontology (GO) terms for each of the three age-contrasts (i.e., 11 – 2 year, 11 – 7 year, and 7 – 2 year). Values in each column represent the number of DMR-associated genes that are associated with each GO term. Black squares indicate no genes associated with that contrast were assigned the particular GO term.

**Table S6a-c.** Number of CHG hypermethylated DMR-associated genes associated with each biological process (**A)**, molecular function **(B)**, and cellular component **(C)** gene ontology (GO) terms for each of the three age-contrasts (i.e., 11 – 2 year, 11 – 7 year, and 7 – 2 year). Values in each column represent the number of DMR-associated genes that are associated with each GO term. Black squares indicate no genes associated with that contrast were assigned the particular GO term.

| **A. Biological Process GO Term** | **11-2** | **11-7** | **7-2** |
| --- | --- | --- | --- |
| proteolysis | 16 | 16 | 4 |
| obsolete oxidation-reduction process | 7 | 13 | 2 |
| protein phosphorylation | 5 | 11 | 5 |
| carbohydrate metabolic process | 4 | 4 |  |
| regulation of transcription, DNA-templated | 4 | 4 | 2 |
| transmembrane transport | 5 | 4 | 2 |
| positive regulation of stomatal complex development |  | 3 |  |
| biosynthetic process | 1 | 2 | 1 |
| fatty acid biosynthetic process |  | 2 |  |
| nitrogen compound metabolic process | 2 |  | 1 |
| protein ubiquitination |  | 2 |  |
| response to auxin | 2 |  |  |
| signal transduction | 5 | 2 | 1 |
| translational termination | 2 |  | 1 |
| cation transport |  | 1 |  |
| cell population proliferation |  | 1 |  |
| cell redox homeostasis | 1 | 1 |  |
| cell wall modification |  | 1 | 1 |
| DNA topological change | 1 |  |  |
| drug transmembrane transport |  |  | 1 |
| glycerolipid biosynthetic process | 1 |  |  |
| intracellular protein transport |  | 1 |  |
| intracellular signal transduction | 1 | 1 |  |
| ion transport | 1 |  |  |
| lipid metabolic process | 1 | 1 |  |
| mismatch repair | 1 |  |  |
| nucleoside metabolic process | 1 |  | 1 |
| nucleosome assembly |  | 1 |  |
| potassium ion transport |  | 1 |  |
| recognition of pollen |  |  | 1 |
| response to oxidative stress |  | 1 | 2 |
| terpenoid biosynthetic process | 1 |  |  |
| transcription, DNA-templated |  | 1 | 1 |
| translation | 2 | 1 | 1 |
| vesicle-mediated transport |  | 1 |  |

| **B. Molecular Function GO Term** | **11-2** | **11-7** | **7-2** |
| --- | --- | --- | --- |
| cysteine-type peptidase activity | 14 | 15 | 2 |
| protein binding | 14 | 9 | 7 |
| ATP binding | 10 | 7 | 8 |
| zinc ion binding | 8 | 3 | 6 |
| nucleic acid binding | 7 | 4 | 1 |
| ADP binding | 5 | 5 | 1 |
| protein kinase activity | 5 | 11 | 5 |
| hydrolase activity, hydrolyzing O-glycosyl compounds |  | 4 |  |
| catalytic activity | 3 | 2 |  |
| copper ion binding |  | 3 |  |
| DNA binding | 3 | 2 | 6 |
| RNA-DNA hybrid ribonuclease activity | 3 | 1 |  |
| transmembrane transporter activity | 3 | 3 | 1 |
| actin binding | 2 |  | 1 |
| endopeptidase inhibitor activity | 2 | 2 |  |
| GTP binding | 2 |  |  |
| heme binding | 2 | 5 | 2 |
| hydrolase activity, acting on ester bonds | 2 | 8 | 2 |
| hydrolase activity, acting on glycosyl bonds |  | 2 |  |
| iron ion binding | 2 | 4 |  |
| nickel cation binding | 2 |  |  |
| oxidoreductase activity, acting on paired donors, with incorporation or reduction of molecular oxygen | 2 | 4 |  |
| polygalacturonase activity | 2 |  |  |
| protein dimerization activity | 2 | 3 |  |
| RNA binding |  | 2 |  |
| sequence-specific DNA binding | 2 | 2 |  |
| serine-type endopeptidase activity | 2 | 1 |  |
| structural constituent of ribosome | 2 | 1 | 1 |
| translation release factor activity | 2 |  | 1 |
| ubiquitin-protein transferase activity |  | 2 |  |
| 2-C-methyl-D-erythritol 2,4-cyclodiphosphate synthase activity | 1 |  |  |
| acid phosphatase activity |  |  | 1 |
| alpha-amylase activity | 1 |  |  |
| antioxidant activity |  | 1 |  |
| ATP-dependent peptidase activity | 1 |  |  |
| ATPase activity |  |  | 1 |
| calcium ion binding | 1 | 1 |  |
| diacylglycerol O-acyltransferase activity | 1 |  |  |
| DNA topoisomerase activity | 1 |  |  |
| DNA-binding transcription factor activity | 1 | 1 | 1 |
| DNA-directed 5'-3' RNA polymerase activity |  |  | 1 |
| double-stranded DNA binding | 1 |  |  |
| electron transfer activity | 1 | 1 |  |
| endonuclease activity |  | 1 |  |
| enzyme inhibitor activity |  | 1 | 1 |
| flavin adenine dinucleotide binding | 1 |  |  |
| growth factor activity |  | 1 |  |
| GTPase activator activity | 1 | 1 |  |
| hydrolase activity |  | 1 | 2 |
| ion channel activity | 1 |  |  |
| ionotropic glutamate receptor activity | 1 |  |  |
| lyase activity |  | 1 |  |
| magnesium ion binding |  | 1 |  |
| malonyl-CoA decarboxylase activity |  | 1 |  |
| metal ion binding |  | 1 | 2 |
| mismatched DNA binding | 1 |  |  |
| NAD binding |  | 1 |  |
| nutrient reservoir activity | 1 |  |  |
| oxidoreductase activity | 1 | 6 |  |
| oxidoreductase activity, acting on CH-OH group of donors | 1 |  |  |
| oxidoreductase activity, acting on the aldehyde or oxo group of donors, NAD or NADP as acceptor | 1 |  |  |
| oxidoreductase activity, acting on the CH-CH group of donors, NAD or NADP as acceptor |  | 1 |  |
| pectinesterase activity |  | 1 | 1 |
| peptide-methionine (S)-S-oxide reductase activity | 1 | 1 |  |
| peroxidase activity |  | 1 | 2 |
| phosphotransferase activity, alcohol group as acceptor | 1 |  |  |
| polysaccharide binding | 1 |  |  |
| protein disulfide oxidoreductase activity | 1 | 1 |  |
| protein phosphatase regulator activity | 1 | 1 |  |
| racemase activity, acting on amino acids and derivatives |  |  | 1 |
| ribonuclease activity |  | 1 |  |
| serine-type carboxypeptidase activity | 1 |  | 2 |
| solute:proton antiporter activity |  | 1 |  |
| strictosidine synthase activity |  |  | 1 |
| terpene synthase activity |  | 1 |  |
| transferase activity, transferring acyl groups |  | 1 | 1 |
| transferase activity, transferring acyl groups other than amino-acyl groups | 1 | 2 | 2 |
| transferase activity, transferring hexosyl groups | 1 |  | 1 |
| ubiquitin-like modifier activating enzyme activity | 1 |  | 1 |

| **C. Cellular Component GO Term** | **11-2** | **11-7** | **7-2** |
| --- | --- | --- | --- |
| membrane | 5 | 4 | 1 |
| integral component of membrane | 1 | 3 | 1 |
| AP-5 adaptor complex |  | 1 |  |
| COPI vesicle coat |  | 1 |  |
| extracellular region |  | 1 |  |
| membrane coat |  | 1 |  |
| nucleus | 2 | 1 | 4 |
| protein phosphatase type 2A complex | 1 | 1 |  |
| proton-transporting two-sector ATPase complex, catalytic domain |  | 1 |  |
| ribosome | 2 | 1 | 1 |
| membrane |  | 4 |  |

**Table S7a-c.** Number of CHG hypomethylated DMR-associated genes associated with each biological process (**A)**, molecular function **(B)**, and cellular component **(C)** gene ontology (GO) terms for each of the three age-contrasts (i.e. 11 – 2 year, 11 – 7 year, and 7 – 2 year). Values in each column represent the number of DMR-associated genes that are associated with each GO term. Black squares indicate no genes associated with that contrast were assigned the particular GO term.

| **A. Biological Process GO Term** | **11-2** | **11-7** | **7-2** |
| --- | --- | --- | --- |
| obsolete oxidation-reduction process | 5 | 3 | 9 |
| protein phosphorylation | 5 | 5 | 6 |
| proteolysis | 5 | 8 | 8 |
| regulation of transcription, DNA-templated | 4 | 5 | 2 |
| carbohydrate metabolic process | 2 |  | 2 |
| cysteinyl-tRNA aminoacylation | 2 |  |  |
| phospholipid biosynthetic process |  |  | 2 |
| response to auxin | 2 |  | 1 |
| transmembrane transport | 2 | 1 | 5 |
| amine metabolic process | 1 |  | 1 |
| ATP metabolic process | 1 |  |  |
| autophagy | 1 |  |  |
| base-excision repair |  | 1 |  |
| cell redox homeostasis | 1 |  |  |
| defense response |  |  | 1 |
| drug transmembrane transport | 1 | 1 |  |
| G protein-coupled receptor signaling pathway | 1 |  |  |
| intracellular protein transport |  | 1 |  |
| lipid biosynthetic process | 1 |  | 1 |
| magnesium ion transport | 1 |  |  |
| metal ion transport | 1 | 1 |  |
| mismatch repair |  |  | 1 |
| nucleobase-containing compound metabolic process |  | 1 |  |
| nucleosome assembly |  |  | 1 |
| organic substance metabolic process |  |  | 1 |
| proton transmembrane transport | 1 |  |  |
| purine nucleobase biosynthetic process | 1 | 1 |  |
| regulation of cyclin-dependent protein serine/threonine kinase activity |  |  | 1 |
| response to oxidative stress |  |  | 1 |
| rRNA processing |  |  | 1 |
| steroid biosynthetic process |  | 1 |  |
| transcription, DNA-templated |  | 1 |  |
| translation | 1 | 2 | 2 |
| translational elongation | 1 | 1 |  |
| trehalose biosynthetic process |  |  | 1 |
| tRNA processing |  |  | 1 |
| ubiquitin-dependent protein catabolic process |  |  | 1 |

| **B. Molecular Function GO Term** | **11-2** | **11-7** | **7-2** |
| --- | --- | --- | --- |
| ATP binding | 10 | 5 | 10 |
| protein binding | 9 | 3 | 9 |
| zinc ion binding | 8 | 5 | 8 |
| cysteine-type peptidase activity | 5 | 7 | 7 |
| DNA binding | 5 | 5 | 1 |
| protein kinase activity | 5 | 5 | 6 |
| iron ion binding | 4 | 2 | 5 |
| heme binding | 3 | 2 | 5 |
| nucleic acid binding | 3 | 2 | 5 |
| oxidoreductase activity, acting on paired donors, with incorporation or reduction of molecular oxygen | 3 | 2 | 4 |
| transferase activity, transferring acyl groups other than amino-acyl groups | 3 |  |  |
| 3'-5' exonuclease activity |  | 1 |  |
| catalytic activity | 2 | 3 | 2 |
| cysteine-tRNA ligase activity | 2 |  |  |
| DNA-binding transcription factor activity | 2 | 2 | 1 |
| hydrolase activity | 2 | 2 | 1 |
| hydrolase activity, hydrolyzing O-glycosyl compounds | 2 |  | 1 |
| Lys48-specific deubiquitinase activity | 2 |  |  |
| metal ion binding | 2 | 4 |  |
| methyltransferase activity |  | 2 |  |
| nucleotide binding | 2 |  |  |
| phosphatidylserine decarboxylase activity |  |  | 2 |
| sequence-specific DNA binding | 2 | 1 | 1 |
| serine-type endopeptidase activity | 2 |  |  |
| thiol-dependent ubiquitin-specific protease activity | 2 |  |  |
| acid phosphatase activity | 1 | 2 |  |
| ADP binding | 1 | 1 | 2 |
| aminopeptidase activity |  |  | 1 |
| ATPase activity |  |  | 1 |
| calcium ion binding | 1 | 1 |  |
| calcium-dependent phospholipid binding | 1 | 1 |  |
| carbohydrate binding | 1 | 1 |  |
| carbon-sulfur lyase activity |  |  | 1 |
| chromatin binding |  |  | 1 |
| copper ion binding | 1 |  | 1 |
| DNA-3-methyladenine glycosylase activity |  | 1 |  |
| DNA-directed 5'-3' RNA polymerase activity |  | 1 |  |
| electron transfer activity | 1 |  |  |
| endopeptidase inhibitor activity |  |  | 1 |
| enzyme inhibitor activity |  |  | 1 |
| flavin adenine dinucleotide binding |  |  | 1 |
| glycopeptide alpha-N-acetylgalactosaminidase activity | 1 |  |  |
| GTPase activity |  |  | 1 |
| hydrolase activity, acting on ester bonds | 1 | 1 |  |
| intramolecular transferase activity, phosphotransferases |  |  | 1 |
| ionotropic glutamate receptor activity | 1 | 1 |  |
| lyase activity | 1 | 2 |  |
| magnesium ion binding | 1 | 2 |  |
| magnesium ion transmembrane transporter activity | 1 |  |  |
| mismatched DNA binding |  |  | 1 |
| oxidoreductase activity | 1 | 1 | 2 |
| peroxidase activity |  |  | 1 |
| phospholipid binding | 1 | 1 |  |
| phosphoribosylamine-glycine ligase activity | 1 | 1 |  |
| polysaccharide binding |  | 1 |  |
| primary amine oxidase activity | 1 |  | 1 |
| protein dimerization activity | 1 |  | 2 |
| protein disulfide oxidoreductase activity | 1 |  |  |
| protein kinase binding |  |  | 1 |
| protein serine/threonine kinase activity |  | 1 | 1 |
| quinone binding | 1 |  | 1 |
| RNA-DNA hybrid ribonuclease activity | 1 | 1 | 2 |
| RNA binding |  | 1 |  |
| serine-type carboxypeptidase activity |  | 1 |  |
| structural constituent of ribosome | 1 | 2 | 2 |
| terpene synthase activity | 1 | 2 |  |
| transferase activity, transferring glycosyl groups |  |  | 1 |
| transferase activity, transferring hexosyl groups | 1 | 1 |  |
| translation elongation factor activity | 1 | 1 |  |
| transmembrane transporter activity | 1 |  | 3 |
| tRNA dihydrouridine synthase activity |  |  | 1 |
| ubiquitin protein ligase binding |  |  | 1 |

| **C. Cellular Component GO Term** | **11-2** | **11-7** | **7-2** |
| --- | --- | --- | --- |
| integral component of membrane | 4 | 1 | 3 |
| cytoplasm | 3 |  |  |
| membrane | 1 | 2 | 3 |
| nucleus | 1 | 1 | 3 |
| ribosome | 1 | 2 | 2 |
| anchored component of plasma membrane |  |  | 1 |

**Table S8a-c.** Number of CHH hypermethylated DMR-associated genes associated with each biological process (**A)**, molecular function **(B)**, and cellular component **(C)** gene ontology (GO) terms for each of the three age-contrasts (i.e., 11 – 2 year, 11 – 7 year, and 7 – 2 year). Values in each column represent the number of DMR-associated genes that are associated with each GO term. Black squares indicate no genes associated with that contrast were assigned the particular GO term.

| **A. Biological Process GO Term** | **11-2** | **11-7** | **7-2** |
| --- | --- | --- | --- |
| protein phosphorylation | 14 | 7 | 28 |
| proteolysis | 13 | 6 | 21 |
| obsolete oxidation-reduction process | 8 | 4 | 20 |
| regulation of transcription, DNA-templated | 7 | 2 | 15 |
| translation | 1 | 2 | 8 |
| transmembrane transport | 5 | 4 | 8 |
| signal transduction | 3 | 1 | 6 |
| carbohydrate metabolic process | 2 | 2 | 4 |
| defense response |  | 1 | 3 |
| tRNA aminoacylation for protein translation | 1 |  | 3 |
| cation transport |  | 2 |  |
| cell redox homeostasis | 1 |  | 2 |
| cell wall modification |  | 1 | 2 |
| DNA topological change |  |  | 2 |
| exocytosis |  |  | 2 |
| fatty acid beta-oxidation |  |  | 2 |
| histone lysine methylation |  |  | 2 |
| lipid metabolic process |  | 2 | 2 |
| nitrate assimilation | 3 |  | 2 |
| positive regulation of Notch signaling pathway | 2 |  |  |
| protein ubiquitination |  |  | 2 |
| pyrimidine nucleotide biosynthetic process | 2 |  |  |
| transcription, DNA-templated |  |  | 2 |
| ATP metabolic process |  |  | 1 |
| biosynthetic process | 1 |  | 1 |
| carboxylic acid metabolic process |  | 1 |  |
| cell wall biogenesis | 1 |  |  |
| cell wall macromolecule catabolic process |  |  | 1 |
| cellular amino acid metabolic process |  | 1 |  |
| cellular glucan metabolic process |  | 1 |  |
| cellulose biosynthetic process |  |  | 1 |
| cellulose microfibril organization | 1 |  | 1 |
| chitin catabolic process |  |  | 1 |
| DNA-templated transcription, initiation |  | 1 |  |
| drug transmembrane transport |  |  | 1 |
| endocytosis |  |  | 1 |
| fatty acid biosynthetic process |  | 1 | 1 |
| glutathione biosynthetic process |  | 1 |  |
| glycolytic process | 1 |  | 1 |
| glycyl-tRNA aminoacylation |  |  | 1 |
| guanosine tetraphosphate metabolic process | 1 |  |  |
| intermembrane lipid transfer |  |  | 1 |
| intracellular protein transport | 1 |  | 1 |
| intracellular signal transduction |  |  | 1 |
| ion transport | 1 |  | 1 |
| lipid catabolic process |  | 2 | 1 |
| malate transport |  | 1 |  |
| mannose metabolic process | 2 |  | 1 |
| mRNA export from nucleus | 2 |  | 1 |
| nucleobase-containing compound metabolic process |  |  | 1 |
| nucleosome assembly |  |  | 1 |
| phosphatidylinositol metabolic process | 1 |  | 1 |
| phospholipid biosynthetic process |  | 1 |  |
| photosynthesis |  |  | 1 |
| proteasome assembly | 2 |  | 1 |
| protein kinase C-activating G protein-coupled receptor signaling pathway | 1 |  |  |
| protein neddylation |  |  | 1 |
| protein peptidyl-prolyl isomerization |  | 1 |  |
| proteolysis involved in cellular protein catabolic process |  | 1 |  |
| proton transmembrane transport |  |  | 1 |
| regulation of systemic acquired resistance | 1 |  |  |
| RNA processing | 1 |  |  |
| response to oxidative stress | 1 |  |  |
| retrograde vesicle-mediated transport, Golgi to endoplasmic reticulum |  |  | 1 |
| ribosome biogenesis |  | 1 | 1 |
| rRNA processing |  |  | 1 |
| sulfate transport |  | 1 |  |
| terpenoid biosynthetic process | 1 |  |  |
| transcription by RNA polymerase II |  |  | 1 |
| transcription initiation from RNA polymerase II promoter | 1 | 1 | 1 |
| translational elongation |  |  | 1 |
| tryptophan catabolic process to kynurenine |  |  | 1 |
| tryptophan metabolic process |  | 1 |  |
| ubiquitin-dependent protein catabolic process |  | 1 | 1 |
| valyl-tRNA aminoacylation |  |  | 1 |
| vesicle docking involved in exocytosis |  |  | 1 |
| vesicle-mediated transport | 1 |  | 1 |

| **B. Molecular Function GO Term** | **11-2** | **11-7** | **7-2** |
| --- | --- | --- | --- |
| protein binding | 20 | 6 | 42 |
| ATP binding | 21 | 10 | 37 |
| protein kinase activity | 14 | 7 | 28 |
| cysteine-type peptidase activity | 12 | 4 | 20 |
| DNA binding | 8 | 1 | 19 |
| ADP binding | 5 | 3 | 16 |
| nucleic acid binding | 11 | 5 | 15 |
| zinc ion binding | 8 | 2 | 14 |
| hydrolase activity, acting on ester bonds | 5 | 4 | 13 |
| protein dimerization activity | 5 | 1 | 13 |
| structural constituent of ribosome | 1 | 2 | 8 |
| oxidoreductase activity | 7 | 1 | 7 |
| DNA-binding transcription factor activity | 2 | 1 | 6 |
| RNA binding | 1 |  | 6 |
| acetylglucosaminyltransferase activity | 2 | 1 | 5 |
| RNA-DNA hybrid ribonuclease activity | 4 |  | 5 |
| 3-hydroxyisobutyryl-CoA hydrolase activity |  |  | 4 |
| heme binding | 2 | 1 | 4 |
| iron ion binding | 1 | 1 | 4 |
| nucleotide binding | 1 |  | 4 |
| oxidoreductase activity, acting on paired donors, with incorporation or reduction of molecular oxygen | 1 | 1 | 4 |
| polysaccharide binding | 1 | 2 | 4 |
| sequence-specific DNA binding | 2 | 1 | 4 |
| transferase activity, transferring acyl groups | 2 |  | 4 |
| aminoacyl-tRNA ligase activity | 1 |  | 3 |
| ATPase activity | 5 | 1 | 3 |
| calcium ion binding | 1 |  | 3 |
| catalytic activity | 4 | 3 | 3 |
| electron transfer activity | 1 |  | 3 |
| GTPase activity | 3 |  |  |
| hydrolase activity |  |  | 3 |
| oxidoreductase activity, acting on the CH-CH group of donors |  |  | 3 |
| serine-type endopeptidase activity | 1 | 1 | 3 |
| transferase activity, transferring hexosyl groups | 4 | 1 | 3 |
| ubiquitin-like modifier activating enzyme activity | 1 |  | 3 |
| acyl-CoA oxidase activity |  |  | 2 |
| DNA topoisomerase activity |  |  | 2 |
| flavin adenine dinucleotide binding |  |  | 2 |
| GTP binding | 5 | 1 | 2 |
| helicase activity |  |  | 2 |
| histone-lysine N-methyltransferase activity |  |  | 2 |
| hydrolase activity, hydrolyzing O-glycosyl compounds |  | 1 | 2 |
| metal ion binding |  | 2 | 2 |
| methyltransferase activity |  |  | 2 |
| molybdenum ion binding | 3 |  | 2 |
| nutrient reservoir activity | 2 |  |  |
| oxidoreductase activity, acting on CH-OH group of donors |  |  | 2 |
| pectinesterase activity |  | 1 | 2 |
| protein disulfide oxidoreductase activity | 1 |  | 2 |
| solute:proton antiporter activity |  | 2 |  |
| transferase activity, transferring glycosyl groups |  |  | 2 |
| ubiquitin-protein transferase activity |  |  | 2 |
| 1-deoxy-D-xylulose-5-phosphate synthase activity | 1 |  |  |
| 3'-5' exonuclease activity |  |  | 1 |
| alpha-amylase activity | 1 |  |  |
| alpha-mannosidase activity | 2 |  | 1 |
| antioxidant activity |  |  | 1 |
| arylformamidase activity |  |  | 1 |
| ATPase-coupled transmembrane transporter activity |  |  | 1 |
| calcium-dependent phospholipid binding |  |  | 1 |
| carboxy-lyase activity |  | 1 |  |
| cellulose synthase (UDP-forming) activity |  |  | 1 |
| channel activity |  | 1 | 1 |
| chitin binding |  |  | 1 |
| chitinase activity |  |  | 1 |
| chromatin binding |  | 2 | 1 |
| copper ion binding | 1 |  | 1 |
| cysteine-type endopeptidase inhibitor activity |  |  | 1 |
| diacylglycerol kinase activity | 1 |  |  |
| DNA topoisomerase type II (double strand cut, ATP-hydrolyzing) activity |  |  | 1 |
| DNA-directed 5'-3' RNA polymerase activity |  |  | 1 |
| endopeptidase activity |  | 1 |  |
| enzyme inhibitor activity |  | 2 | 1 |
| galactoside 2-alpha-L-fucosyltransferase activity | 1 |  |  |
| glutathione synthase activity |  | 1 |  |
| glycine-tRNA ligase activity |  |  | 1 |
| glycopeptide alpha-N-acetylgalactosaminidase activity | 1 |  |  |
| heat shock protein binding |  |  | 1 |
| ion channel activity | 1 |  | 1 |
| lipid transfer activity |  |  | 1 |
| magnesium ion binding | 2 |  | 1 |
| mannosyl-glycoprotein endo-beta-N-acetylglucosaminidase activity |  |  | 1 |
| microtubule minus-end binding |  |  | 1 |
| NEDD8 activating enzyme activity |  |  | 1 |
| O-methyltransferase activity |  |  | 1 |
| obsolete coenzyme binding |  | 1 | 1 |
| peptidyl-prolyl cis-trans isomerase activity |  | 1 |  |
| peroxidase activity | 1 |  |  |
| phosphatidylinositol binding |  |  | 1 |
| phosphatidylinositol phosphate kinase activity | 1 |  | 1 |
| phosphatidylserine decarboxylase activity |  | 1 |  |
| polygalacturonase activity |  |  | 1 |
| potassium ion binding | 1 |  | 1 |
| protein serine/threonine kinase activity |  | 1 |  |
| proton-transporting ATPase activity, rotational mechanism |  | 1 |  |
| pyridoxal phosphate binding |  | 1 | 1 |
| pyruvate kinase activity | 1 |  | 1 |
| ribonuclease T2 activity |  |  | 1 |
| serine-type carboxypeptidase activity | 1 | 1 |  |
| serine-type exopeptidase activity |  | 1 |  |
| serine-type peptidase activity |  | 1 |  |
| sulfate transmembrane transporter activity |  | 1 |  |
| TBP-class protein binding |  | 1 |  |
| threonine-type endopeptidase activity |  | 1 |  |
| thiamine pyrophosphate binding | 1 |  |  |
| thiolester hydrolase activity |  |  | 1 |
| transaminase activity |  |  | 1 |
| transferase activity, transferring acyl groups other than amino-acyl groups | 1 | 2 | 1 |
| translation elongation factor activity |  |  | 1 |
| transmembrane transporter activity | 2 | 1 | 1 |
| tRNA binding |  |  | 1 |
| tryptophan synthase activity |  | 1 |  |
| unfolded protein binding |  |  | 1 |
| valine-tRNA ligase activity |  |  | 1 |
| xyloglucan:xyloglucosyl transferase activity |  | 1 |  |

| **C. Cellular Component GO Term** | **11-2** | **11-7** | **7-2** |
| --- | --- | --- | --- |
| integral component of membrane | 3 | 5 | 15 |
| membrane | 10 | 4 | 14 |
| ribosome | 1 | 2 | 8 |
| nucleus |  | 1 | 7 |
| cytoplasm |  |  | 4 |
| chloroplast |  |  | 2 |
| chromosome |  |  | 2 |
| exocyst |  |  | 2 |
| peroxisome |  |  | 2 |
| anchored component of membrane | 1 |  | 1 |
| apoplast |  | 1 |  |
| cell wall |  | 1 |  |
| endoplasmic reticulum membrane | 1 |  |  |
| extrinsic component of membrane |  |  | 1 |
| Golgi transport complex |  |  | 1 |
| nucleolus |  | 1 |  |
| nucleosome |  |  | 1 |
| photosystem II |  |  | 1 |
| photosystem II oxygen evolving complex |  |  | 1 |
| proteasome core complex |  | 1 |  |
| proteasome core complex, alpha-subunit complex |  | 1 |  |
| proteasome regulatory particle, lid subcomplex | 2 |  | 1 |
| proton-transporting two-sector ATPase complex, catalytic domain |  | 1 |  |
| transcription factor TFIIA complex | 1 |  | 1 |

**Table S9a-c.** Number of CHH hypomethylated DMR-associated genes associated with each biological process (**A)**, molecular function **(B)**, and cellular component **(C)** gene ontology (GO) terms for each of the three age-contrasts (i.e., 11 – 2 year, 11 – 7 year, and 7 – 2 year). Values in each column represent the number of DMR-associated genes that are associated with each GO term. Black squares indicate no genes associated with that contrast were assigned the particular GO term.

| **A. Biological Process GO Term** | **11-2** | **11-7** | **7-2** |
| --- | --- | --- | --- |
| protein phosphorylation | 2 | 9 | 1 |
| regulation of transcription, DNA-templated | 6 | 9 |  |
| proteolysis | 2 | 5 | 4 |
| transmembrane transport | 2 | 5 |  |
| carbohydrate metabolic process |  | 3 | 1 |
| obsolete oxidation-reduction process | 5 | 3 | 6 |
| translation | 1 | 3 |  |
| ATP metabolic process |  | 2 |  |
| base-excision repair | 1 | 2 |  |
| nucleobase-containing compound metabolic process |  | 2 |  |
| proton transmembrane transport |  | 2 |  |
| response to hormone |  | 2 |  |
| response to oxidative stress | 2 |  | 2 |
| cell redox homeostasis |  |  | 1 |
| cell wall macromolecule catabolic process |  | 1 |  |
| chitin catabolic process |  | 1 |  |
| defense response |  | 1 |  |
| DNA repair |  | 1 |  |
| DNA topological change |  |  | 1 |
| exocytosis |  | 1 |  |
| intermembrane lipid transfer |  |  | 1 |
| intracellular protein transport | 1 | 1 |  |
| lipid metabolic process | 1 |  | 1 |
| metal ion transport | 1 | 1 | 1 |
| nitrogen compound metabolic process |  |  | 1 |
| phytochromobilin biosynthetic process |  | 1 |  |
| protein folding | 1 |  |  |
| rRNA processing |  |  | 1 |
| signal transduction |  | 1 |  |
| steroid biosynthetic process |  |  | 1 |
| sucrose metabolic process |  |  | 1 |
| superoxide metabolic process | 1 |  |  |
| translational initiation |  | 1 |  |

| **B. Molecular Function GO Term** | **11-2** | **11-7** | **7-2** |
| --- | --- | --- | --- |
| ATP binding | 3 | 14 | 3 |
| DNA binding | 5 | 13 | 2 |
| protein binding | 4 | 9 | 3 |
| protein kinase activity | 2 | 9 | 1 |
| zinc ion binding |  | 6 | 2 |
| cysteine-type peptidase activity | 2 | 5 | 4 |
| RNA binding |  | 5 |  |
| sequence-specific DNA binding | 4 | 5 |  |
| DNA-binding transcription factor activity | 5 | 4 |  |
| metal ion binding | 1 | 4 |  |
| ATPase activity |  | 3 |  |
| ATPase-coupled transmembrane transporter activity |  | 3 |  |
| protein dimerization activity | 1 | 3 | 1 |
| structural constituent of ribosome | 1 | 3 |  |
| 3'-5' exonuclease activity |  | 2 |  |
| ADP binding | 1 | 2 |  |
| DNA-3-methyladenine glycosylase activity | 1 | 2 |  |
| glutathione peroxidase activity | 2 |  | 2 |
| heme binding |  |  | 2 |
| hydrolase activity | 1 | 2 | 2 |
| hydrolase activity, acting on ester bonds |  | 2 | 3 |
| iron ion binding |  |  | 2 |
| nucleic acid binding |  | 2 | 4 |
| oxidoreductase activity, acting on paired donors, with incorporation or reduction of molecular oxygen |  |  | 2 |
| polygalacturonase activity |  | 2 |  |
| pyridoxal phosphate binding |  | 2 |  |
| transaminase activity |  | 2 |  |
| 3-beta-hydroxy-delta5-steroid dehydrogenase activity |  |  | 1 |
| acetylglucosaminyltransferase activity | 1 | 1 | 2 |
| acid phosphatase activity |  | 1 |  |
| carbohydrate binding |  | 1 |  |
| catalytic activity | 1 | 1 | 2 |
| chitin binding |  | 1 |  |
| chitinase activity |  | 1 |  |
| cobalt ion binding |  | 1 |  |
| DNA topoisomerase activity |  |  | 1 |
| flavin adenine dinucleotide binding | 1 |  |  |
| hydrolase activity, hydrolyzing O-glycosyl compounds |  | 1 | 1 |
| intramolecular lyase activity |  | 1 |  |
| lipid transfer activity |  |  | 1 |
| Lys48-specific deubiquitinase activity |  | 1 |  |
| metal ion transmembrane transporter activity | 1 |  | 1 |
| methyltransferase activity |  | 1 | 1 |
| microtubule minus-end binding |  | 1 |  |
| O-methyltransferase activity |  | 1 |  |
| obsolete coenzyme binding |  |  | 1 |
| oxidoreductase activity | 2 |  | 1 |
| oxidoreductase activity, acting on the CH-OH group of donors, NAD or NADP as acceptor |  |  | 1 |
| oxidoreductase activity, acting on single donors with incorporation of molecular oxygen, incorporation of two atoms of oxygen |  | 1 |  |
| oxidoreductase activity, acting on the CH-CH group of donors, iron-sulfur protein as acceptor |  | 1 |  |
| polysaccharide binding | 1 |  | 1 |
| racemase activity, acting on amino acids and derivatives |  |  | 1 |
| sucrose synthase activity |  |  | 1 |
| superoxide dismutase activity | 1 |  |  |
| thiol-dependent ubiquitin-specific protease activity |  | 1 |  |
| transferase activity, transferring acyl groups other than amino-acyl groups |  | 1 |  |
| transferase activity, transferring hexosyl groups | 1 | 1 |  |
| translation initiation factor activity |  | 1 |  |

| **C. Cellular Component GO Term** | **11-2** | **11-7** | **7-2** |
| --- | --- | --- | --- |
| integral component of membrane |  | 5 |  |
| ribosome | 1 | 3 |  |
| membrane | 2 | 2 | 4 |
| nucleus |  | 2 | 1 |
| cytoplasm |  |  | 1 |
| exocyst |  | 1 |  |

**Table S10.**
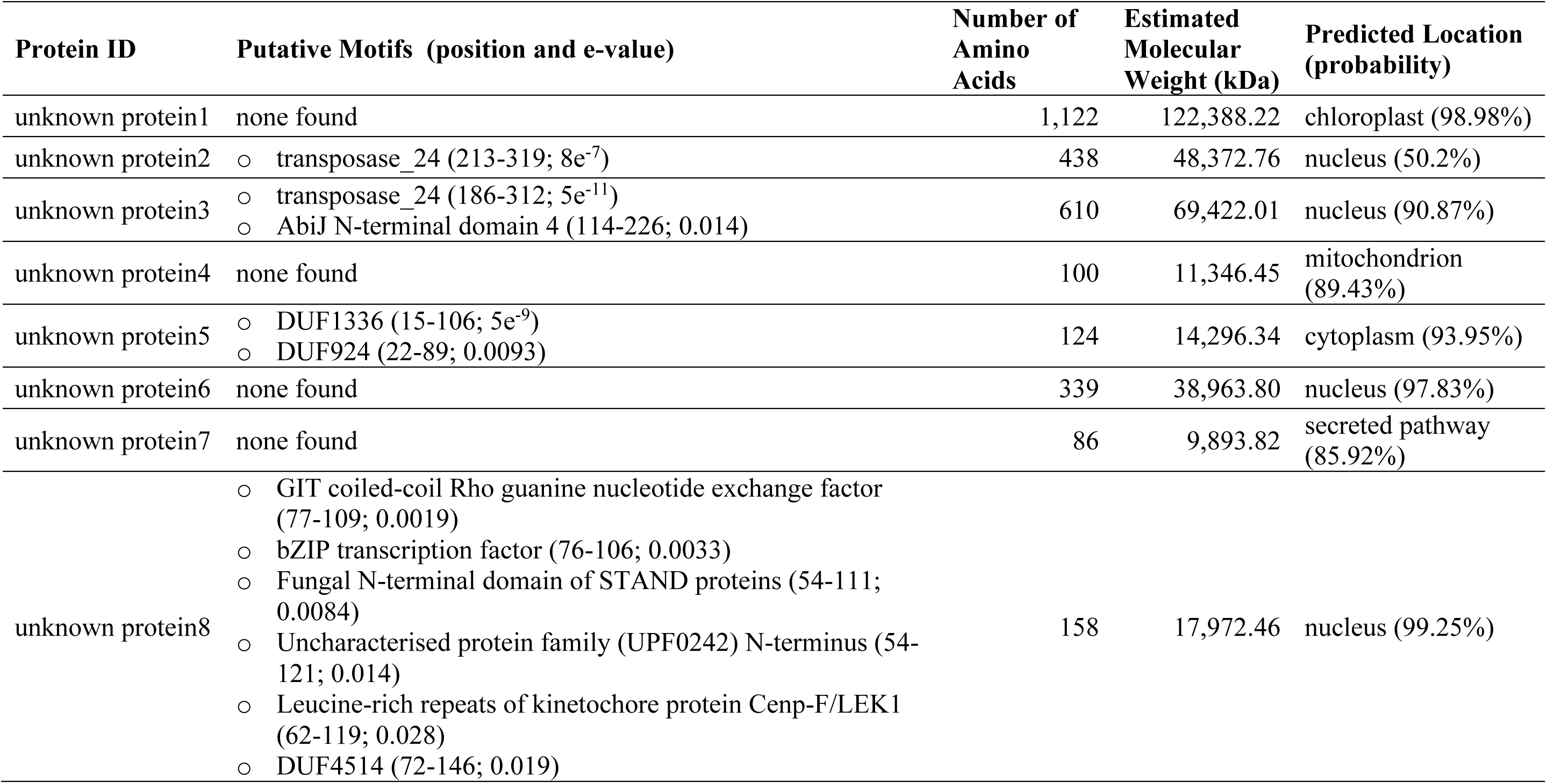
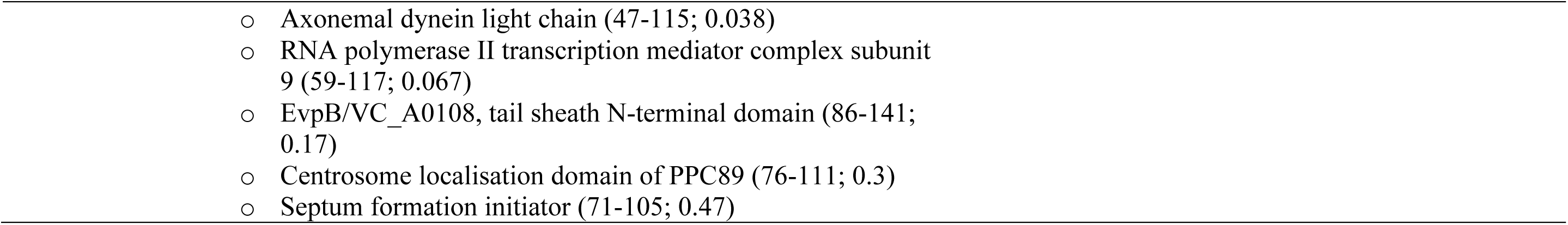
Characterization of eight unknown proteins associated with the 17 shared DMR sequences identified among the three age contrasts. The unknown protein ID corresponds to the unknown proteins associated with the shared DMR sequences. The putative motifs identified within each protein sequence include the position of the motif within the sequence and the e-value. The number of amino acids and estimated molecular weight (kDa) are also included for each protein sequence. Finally, the predicted localization of each protein is provided, as well as the calculated probability of this prediction.

| Protein ID | Putative Motifs (position and e-value) | Number of Amino Acids | Estimated Molecular Weight (kDa) | Predicted Location (probability) |
| --- | --- | --- | --- | --- |
| unknown protein1 | none found | 1,122 | 122,388.22 | chloroplast (98.98%) |
| unknown protein2 | ○ transposase_24 (213-319; $8e^{-7}$) | 438 | 48,372.76 | nucleus (50.2%) |
| unknown protein3 | ○ transposase_24 (186-312; $5e^{-11}$)<br>○ AbiJ N-terminal domain 4 (114-226; 0.014) | 610 | 69,422.01 | nucleus (90.87%) |
| unknown protein4 | none found | 100 | 11,346.45 | mitochondrion (89.43%) |
| unknown protein5 | ○ DUF1336 (15-106; $5e^{-9}$)<br>○ DUF924 (22-89; 0.0093) | 124 | 14,296.34 | cytoplasm (93.95%) |
| unknown protein6 | none found | 339 | 38,963.80 | nucleus (97.83%) |
| unknown protein7 | none found | 86 | 9,893.82 | secreted pathway (85.92%) |
| unknown protein8 | ○ GIT coiled-coil Rho guanine nucleotide exchange factor (77-109; 0.0019)<br>○ bZIP transcription factor (76-106; 0.0033)<br>○ Fungal N-terminal domain of STAND proteins (54-111; 0.0084)<br>○ Uncharacterised protein family (UPF0242) N-terminus (54-121; 0.014)<br>○ Leucine-rich repeats of kinetochore protein Cenp-F/LEK1 (62-119; 0.028)<br>○ DUF4514 (72-146; 0.019) | 158 | 17,972.46 | nucleus (99.25%) |

**Table S11.**
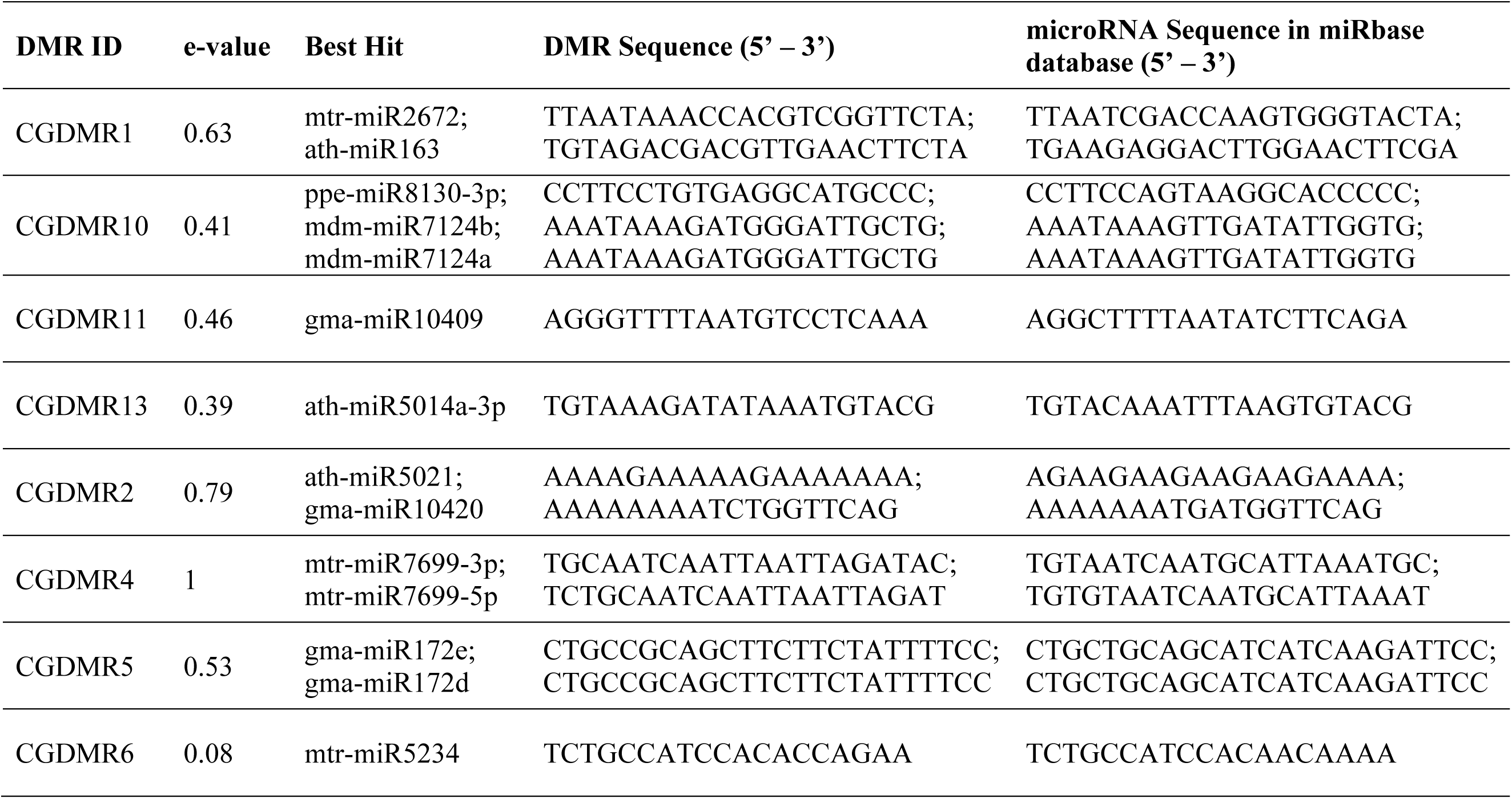
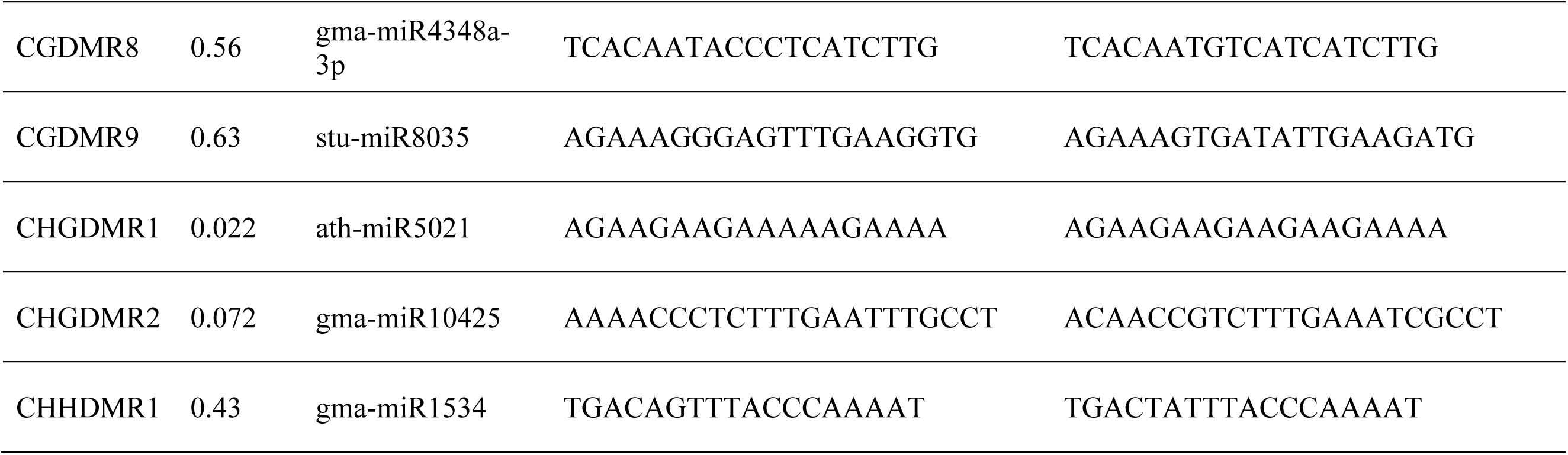
Identification of microRNAs found within the 17 identified DMR sequences based on BLAST analysis using a modified miRbase database containing plant-only entries. The BLAST results were filtered to include only those regions within the DMR that are n – 2 bp in length in relation to the closest associated microRNA and are the best hit based on the associated e-value to each microRNA within the database.s

## Data S1

**Hypermethylation is associated with increased age in almond (Prunus dulcis [Mill.] D.A. Webb) accessions**

**Katherine M. D’Amico-Willman1, Chad E. Niederhuth2, Elizabeth S. Anderson3, Thomas M. Gradziel4, Jonathan Fresnedo Ramírez1,3* 1Translational Plant Sciences Graduate Program, The Ohio State University, Columbus, OH 43210**

**2Department of Plant Biology, Michigan State University, East Lansing, MI 48824**

**3Department of Horticulture and Crop Science, Ohio Agricultural Research and Development Center, The Ohio State University, Wooster, OH 44691**

**4Department of Plant Sciences, University of California, Davis, CA 95616**

**\*For correspondence**

DataS1. File containing sequnecing statistics, conversion efficiencies, total percent methylation, and percent methylation within each context (CG, CHG, CHH) for almond accessions presented in this study.

Contents:

**AgeCohort_MethylSeq**:

samp_ID - the unique sample identifier assigned to each almond accession; age - in years; num_reads - the number of reads generated in total for each individual; cov_seq - the coverage across the reference ’Nonpareil’ almond genome based on the number of reads generated by HiSeq4000 sequencing; num_reads_align - the number of reads that aligned to the reference ’Nonpareil’ almond genome; cov_align - the coverage across the reference ’Nonpareil’ almond genome based on the number of reads that aligned following Bismark alignment; per_align - mapping efficiency; per_CG - percent methylation in the CG context; per_CHG - percent methylation in the CHG context; per_CHH - percent methylation in the CHH context; per_lambda - percent methylated cytosines in the CG context following alignment to the Lambda reference; per_puc19 - percent methylated cytosines in the CG context following alignment to the pUC19 reference; per_meth - total percent methyaltion following alignment to the ’Nonpareil’ almond reference genome; Female_parent: female progenitor of the accession; Male_parent: male progenitor of the accession.

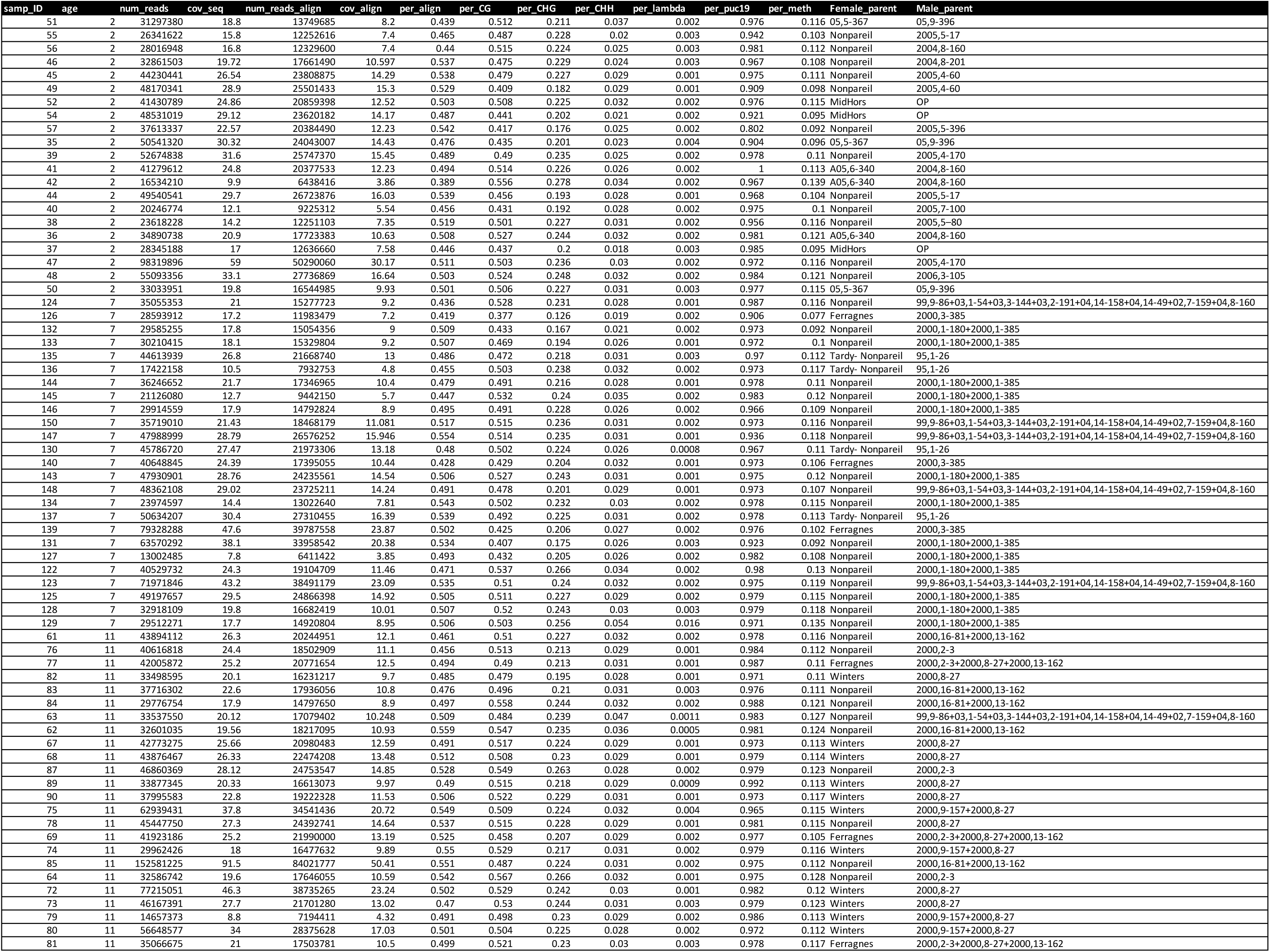

## Data S2

>CGDMR1 CGCCGTTAAATATAATTTTCACGACGTTATGGAAAACTACTCTGTCTCTTATTGAAAACTACTACGTCTTTAAATACATGAGAAGACGACGTTATTGAAAACTACTACGTCTCTTATTTACAAAAGACCGTCGTGAAATAAGTTTTCCACTTCGATTGGTCTAAAAAAGATGACGTAGATATAGCTGTCTGTGTCATTATTTACGACGTATGTATTTATGTTGTTGACGTTGAAATGCGAAACAACGTCGGTGGCATTTATATAGTCGACGTTGTATTTATATGTATTCAGTACTCACGACGTACTTATTAATGTAGACGACGTTGAACTTCTAAACAACGTCAATATTTACGACGCATTTAATAAACCACGTCGGTTCTAAATTAATGTTCAGATATTTAATCTCATTTTTAGACGTCGGTATTATGGGATTGGAGTCGTGTTATTTTGGATTACATGGCTGTTCTTAATCCTTCCCCGACGTGATATCCTGGATAATTGCTTCACAATAGCGTCGTGGTTCTTCATTGTTACGTTGCTTTAAGACGTCGGTAAATATGCTGAATAAATGGTTAACCATAACGTC

>CGDMR2 CGCGCGGCTATACTATTTGTAAATATAAAAAGCCCAGCCCACAGAGCAGGTACCACGTGTCAAAGAGGTACAGCCGGGCGGCTGTACTATTTTTTTAAAAGATAACACGAGTATAGCCGCGCGGCTATACTATTGCTTGGAAAAAAAAAAAAAAAAAAAAAAGAGAATCTGGTATAGCCCGGCCCAGGCGCCATGTGCGCCACGTGGCAAACAAGTAGAGCCGAACGGCTATACTATTTTTAAAAGCAATTAAGGGAAAAATGAGTATAGCCGCACGGCTATACTATTTCTTTAAAAAAGAAAAAGAAAAAAAGGATCTGGTACAGCCCGGCCCAGGCGCCATGTGCGCCACGTGGCAAGCAAGTAGAGCCGAACGGCTACTCTATTTTTAAAAAGCAATTAAAGGAAAACTGAGTATAGCCGCACGGCTATACTATTCCTTGAAAGAAAAAAAAAATCTGGTTCAGCCCGTGCGCGCCACGTGGCAAACAAGTAGAGCCG

>CGDMR3 CGATCGATTTGGTGGAGAAACGAAGGAGAATTTCGAGCTGGAAGTTCGGGTGGCTTTGCAGGAACCTCCGTCGGAAAACATGGTTTTCCGGCCAACTCAGAGCGGCCCCGGGGTTGAGATCTGTCAGGTTGTGACGGCGACACGAAGGCGGACCCGTAGGTACCAACGGCGGAGATCGGCTCGGGCCTGTGGTGGCCGGGCCGTCCCTTGGAAGGCGAAGGGTCGCACGACCCCAAATCGCGCGAAGGAGAGAGAGGGAGGAGAGAGAAAGTGACGGGGAGAAGAGAGAGAGGGGGTATAAAATCTGACTTTTTGGCCAAATTACCATTTTGCCCTTCGCGGTTTTTAGACCATAACTTCTTCGTTACTGCTCCGATTCGGGCCTACTCCGTGTCTACGAACTCCTTTCG

>CGDMR4 CGTCAACGACTCTTCGTCGAGATCCTCCAAGTTATAGTTCGTCTGCAATCAATTAATTAGATACGTTAAGTAATAGAAATATACACAATTTTAAAGAAAAATAATGAAAATGTAAGGTACATACCGACAACTGGCCGCGAACCTCCGCCTTGATCTCGTCAGGCATGACCCTCCAAGACTTCCATTGCATCGGGCAATGGGTCCGCACGACGTGGCCAATGTCGTGGGCCAAGGAGCTATGCAACTCCGCCGTCGGTGCAGCCCGATGGCGCTCGTCGTATCCGATGCTAATACGACTGTTGGTCACCCGGGTGACCTTCGCCGTCTTCAGCTGACGACACGGTCCCCGGGTGTTCTTCTTCG

>CGDMR5 GCGGGGTGCCCCTGATTTTTCAGGGTAAAGTTGGGTGTACTCTTTCTTCGAGTTTGACCAATTAAATTAATTAAAATTAATATAATAGAAAATTATTTCCCTGCGAAACTGCCGCAGCTTCTTCTATTTTCCTGCGAGTTTGACACCCGCCGCTTCTTCTCTATTTTCCTGCGAAACTGCCGCAGCTGGAGGCG

>CGDMR6 CGCTCGTAAGGAGTCCTATTGAAACCAAGAAATATAGGCCTGTCTGCCATCCACACCAGAATAAATGAAGTTTTTTTTACTCTAACCCTTTTCGAAATAATTACAGTTTTGCCATTGATAATGTTTTGATCACCAAATCTTCGTTACAACTCCGTTTCAAGCCTACCGCGTGTCTACAAATTCGTCTTAGTACCACCTACCTAAAAATACCAATCGTGTTCCCAAAATCCTTCCG

>CGDMR7 CGTCGACTATATAAATGCCACCGACGTTGTTTCGCATTTCAACGTCAACAACATAAATACATACGTCGTAAATAATGACACAGACAGCTATATCTACGTCATCTTTTTTAGACAAATCGAAGTGGAAAACTTATTTCACGACGGTCGTTTGTAAATAAGAGACGTAGTAGTTTTCAATAACGTCGTCTTCTCATGTATTTAAAGAC

>CGDMR8 CGTTTTGCCCTCATTCGCAGCAGCAGGCAATCGATTAATGAGATGGCTTGCATAAGTAATCGCCTCAGCCCAAAACGCCTTGCCTAAACCAGCATTAGACAACATACACCGAACTTTCTCAAGCAAAGTACGGTTCATGCGCTCTGCCACCCCATTCTGTTGCGGTGTCTCCCTAACCGTGAAGTGCCTCACAATACCCTCATCTTGACAAACTTTCAAGAAAGGATCAGACTTATATTCACCACCATTGTCTGATCTGAGAGTCTTGATCTTTCGACCGCTTTGC

>CGDMR9 CGTTCCTCGTCAAGTATTAGGGTTAAGGGTGACTGTGGGTATCGCAAGAGAGAAGAAAGGGAGTTTGAAGGTGAGAAGGA TGAGAGTAACTATGACTGGGTATCGCAAGAGAGAAGAGAGGGAGTCTGAAATATTCACGAGGGAGTGAAATTTTTCAGCG

>CGDMR10 GTGCAGCTTTTAATGGGAAACCTGAATATGGCATACCTCCCGAGCCATTAACCGGAGAAGAAGTGCTGCATATGGTTGAAAATGGTGACAGAGTTTGTTGGAAGAAGAAATCAATATTCTTTGATCTCGAGTATTGGAAATACCTTCCTGTGAGGCATGCCCTAGATGTTATGCATATTGAGAAGAATGTTTGCGATAGTATCATTGGTACATTGCTGGAGATCCCTGGAAAAAATAAAGATGGGATTGCTGCTCGATTAGATTTATTGAACATGGGGGTCAAAACTGATTTGCAACCCGAGTATGGAGAAAGACGTACTC

>CGDMR11 CGGCCCGATAAGTAATATATTTATTTTTAAGTAGTTTATAATATAATATAACCATGTATGATTATAGGGTTTTAATGTCCTCAAACTTCGAGTTGTATTTTGGTCCCTCAACTAAATTATTCG

>CGDMR12 CGTAGTTGTATGACGTCACAAGGAGGCCGGAGTAATTTTCGGGGCATTTGTAGGAATTTCCTACTATTTATTACCTATTATCAAGGATTTTTCCACGAAAATAGCCAACCCTAATTGCAAAAACACACCTGGCGCAATCTCAGCAGGTAACCTTTCAGCACTTGCACAAGTGTAC

>CGDMR13 CGAGGGTTTGTCTATATTTCAATATAGAGTATTCAAATAAGCATGTAAAAATCTACTGATGACTGTGCTTGTATAACTGAACAAATATGTAAAGATATAAATGTACG

>CHGDMR1 CAGTAACAAAAAAAAGGGAGAAGGCTTCCACTGAAAAGAAAACAAACGGAAGAAGAGTAAAAAAAAAAGGAAAGTGGCGTCAGCCACTTTGTTTCAAAAGGAAGAAGAAAGCGTCAGCCACTTTGTTTCAAAAGGAAGAAGAAATTCAAAGAAGAAAAAAAAAGAGAAGAGAGTGGCTGATGCCATTTTGCTGAAATTGGAAGAAAAGCAAAGCAAAGGACTTGGTGTTGCAGATCAAGAAGAAGAAAAAGAAAAGTCAAAACAAGGACAAAAAAGAACAAGGTGGCTGACGCCACTTTTGGAGTTGGAAGAAAAAAAGGGACACAAGAGACTTGCTGTTGCAGAGAAAAAATGAGGAAAAGGACAAAAAAAAAAGTCTGAAACCTCTTGGGTGTTGCGGCAGTTTTTCTCTCTTTCTTCTGGCTTTCTTTTCTTCTTGTTTGCAAAGGCTG

>CHGDMR2 CTGGCTTGTGCTTTTTGGCAGCTATCTGGTATTGGGTGTATTGGGATCTAGAAATATTTATTGATGAACATACAGGAAAACCCTCTTTGAATTTGCCTAAGATCTTTGGAATTCATTTATTTCTATCCGGGGTGGCTTGCTTTGGTTTTGGTGCATTTCATGTAACAGGATTGTATGGTCCTGGAAGGTTCTACCTCCTATTTTTTATGAAGAGAATGAATCTTTTTATGGAAGGATCAGAAAAAAATGGGTCCGGACCTCCTGCGAGAATGATTTGGAAGATCCAAAACCAAAAATAGTGGTATTTGCTAGCAACAACATAGTTAGTAAGAGGGATCTTGAACTAAGAAATAGATTCTAGAAGCTAAAAAAGGGTATCC

>CHGDMR3 GCACCTTCTATAGTGCACCAGGTGCATCCATGCATCATTTGTCATTTATAATTGACAGGACCCAACCCAATTTCCACTTTGAAATTCGAGCCAAGTCCTGCGCGTGTCCGACACCTGGCGAATGTCGGGCACAATTGTCCTTTTTACCCTTCTTACTTCAATTCTTCTTTAAAATTGCCTTAGACTTCTGCCGAAAATTCGGCAGAGTCTCCCCTGTATTTTTGACTTATCCCAAAATTTTCACCTG

>CHHDMR1 GTTTGGCGTCCATCCTTTGTATCCCAAAATCGTCATCTCACAGTTAATGATTCTGTGATGATGAATGATGCTACTGCTGTCACAGTAGCTAGGAATTTCATTATTCCAATGGATGAAATGCTGTTGACAGGGAGGTCTGAAGAAGAGGCTATTGAGGACTCAATGGCTTTTAGCATTCAGAGTGCTGCTTCTGTTTCTAACATGGCTGATCGTTTGCGTGCTAGAGCAAACGAGGTTCAGAAGCTAACAACGAAAAATTCGTCTCTCCAAAGAATACTTCATGAGTCTCAAAAAGAGGTTGAGAAACTTAAAGGAGAGAATAATTCCTTGTTGAAACTGGTGAGTTCGTATTCTGTTGATACACAGAGGAAGCTAGACATGCTGCAGGTCTCAAATGAAAGAATTTTGGGAGACCACGAGAGGCTCATGGCTAGGCTTAAGAAGCGCCGTCCTCTTCCTTCAGAGGCTTCCAGAACATAATGTAATTTTATAGATTTTACAGGGCCTGCACCTTCATTGCAGGTGGAAAAATCTATCTGTTGTATGTTCATTTGCTGTTATAATAATTGTACATGTTCTTAAACTTGCATCTGTGGTTTTTACGTCTTTTCAAAATGACGGTTTGGAACCTTGTGCCTTATAGGTTCAAATAACCACATCAAATCTCTCATATTTCCATGCATATGAGCCCAGAGCTTTTGGTCTGGGTTAAACCCAAACACATAATTAATTCGCCATTTTCAAATGGAGAGCATTAACTACAATGTACCCACAACTTCAAGTTTTAGGATCTCTCATATATTTGGATCCATGGGCTTCCGGCCCAGATATAACAAAATATGTGGGGAGCCTCAATTCATTATTTGAGGTTTATATTGATATTATCCATTTCGCGGTGTATTCTTAACAACCGGAATTCACAAAATATATTTCTTCCTTGAGGTGTCGATTATAACAGAATCGAACTTCATTAAATTCATCATCTTCTTATGCCAAAGAAATATGTGGCATATCACAATTTGCAATAATACCTCAAGGGTTGTCCATTTAATTGTTGGAACTTCAGGTTCTCAATACTGTTAGATTTTGAACTTCGGGCCAAAATCACATATTCTCATGGTATGGACATTTTTACAATTTTCTGTACATATTTTGGGACTTCAAGTCCTTACATAATTGTCCATATTTTGAGGAACTTCTGGCATCTCATTTAATTGCTCATCCATGAGTGTAAGGAACTGCAGGTTCCCTTTTGTATATAGTGACAGTTTACCCAAAATGGTTAATATTTATGCATACGTCACTATTCATGTGAATAGTACTATTCATCAATCATGAATACATATCTATTCATATGTGC AGTACATTTGCCAGTACAGTTATTATTCATGTGTACGGCACTTTTAACCAATACGGTACTGTTACATCATTAAGGAAACCCAGGTCCTTATTCACATGTCAGGGATCAAGGACCCTTAAGTCCGATCACATGTTTACAAATATAG

## Data S3

**Katherine M. D’Amico-Willman1, Chad E. Niederhuth2, Elizabeth S. Anderson3, Thomas M. Gradziel4, Jonathan Fresnedo Ramírez1,3\***

**1Translational Plant Sciences Graduate Program, The Ohio State University, Columbus, OH 43210**

**2Department of Plant Biology, Michigan State University, East Lansing, MI 48824**

**4Department of Plant Sciences, University of California, Davis, CA 95616**

**\*For correspondence**

DataS3. File containing general infromation on DMRs overlapping transposable elements.

Contents:

**AgeCohortRep_DMRs_classified**:

Chromosome - chromosome where DMR is located; Start - starting position of the DMR; End - ending position of the DMR; DMR_length- lenght of the DMR; ID - identificator f the DMR; ChrAligned - Chromosome where the transposable element is located; Source - software used for the prediction of TEs (EDTA); TEType - type of transposable element; TeStart - begining of the TE; TEEnd - end of the TE; TE_length - actual length of the TE; TE_ID - identifyier of the specific transposable lement in the Nonpareil almond genome; Parent - main repetitive region for TE; Name - Name of homologous TE in database; Calssifciation - class in which the TE falls, Seq_Ontology - ontology for the TE, Method - whether the TE was predicted based on homology or structural features; Motif - sequence motifi for TE, TSD - Target Site Duplication, TIR - Terminal Inverted Repeat; Overlpa - overlap in bp between the DMR and the TE

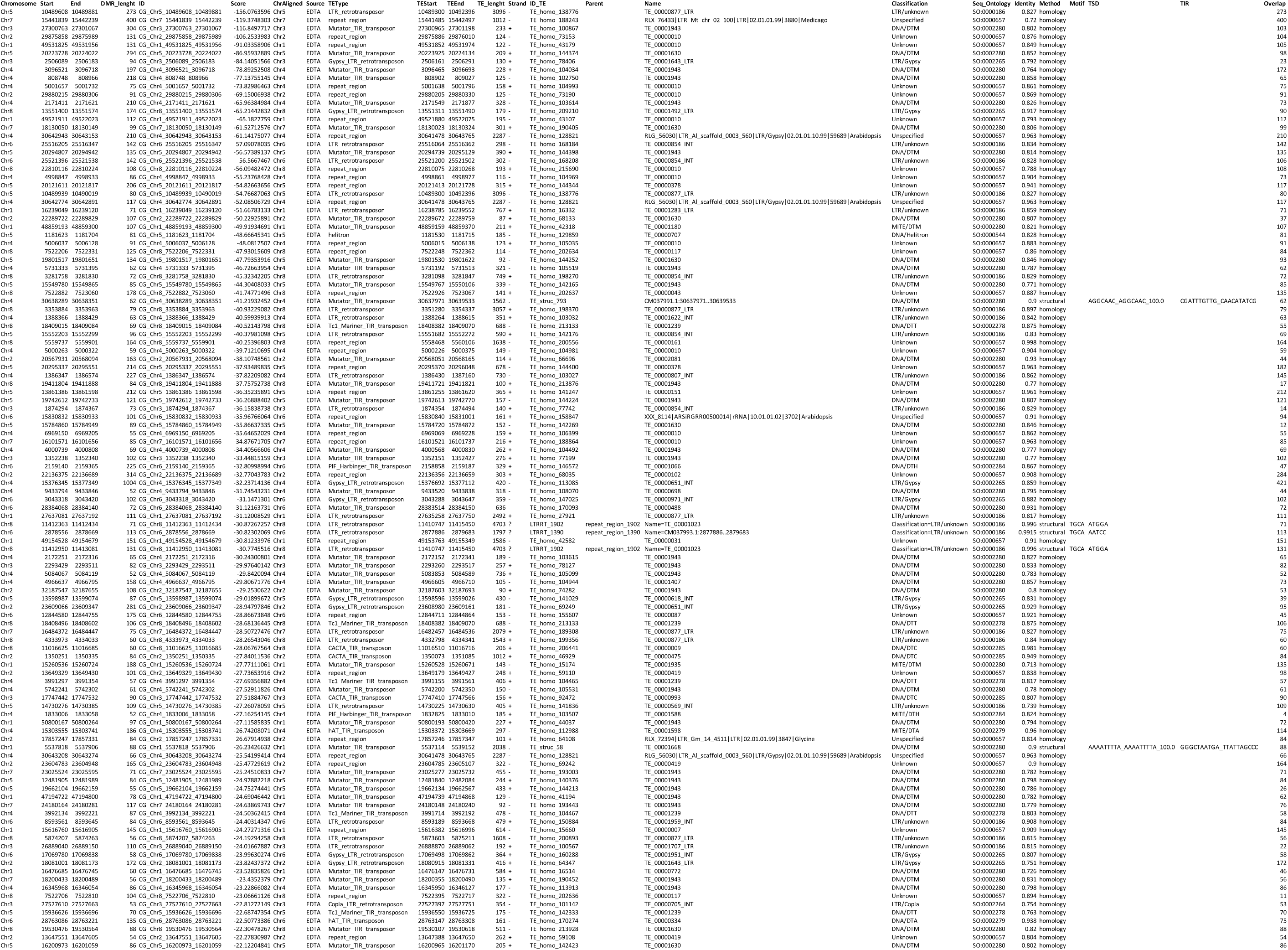
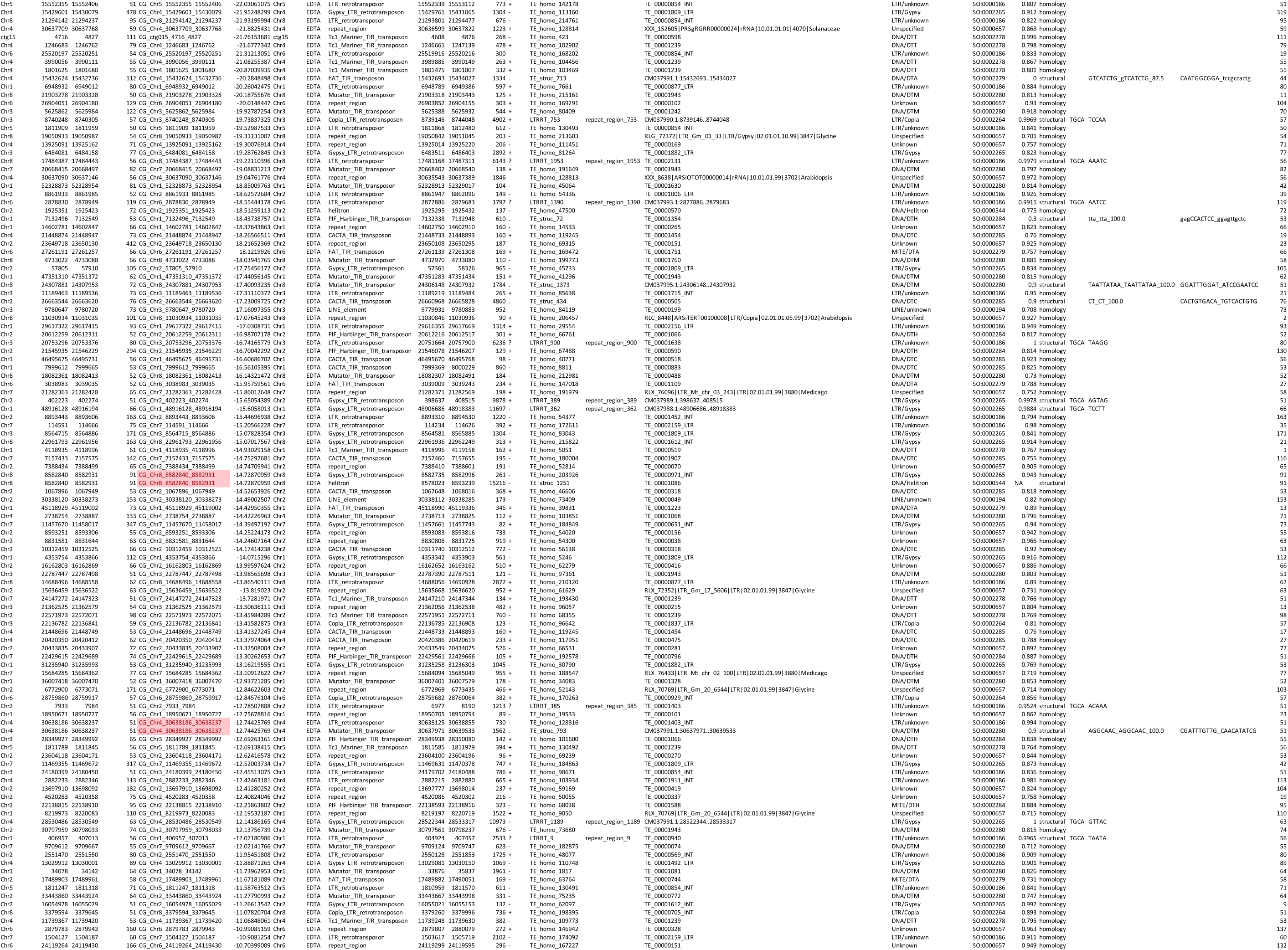
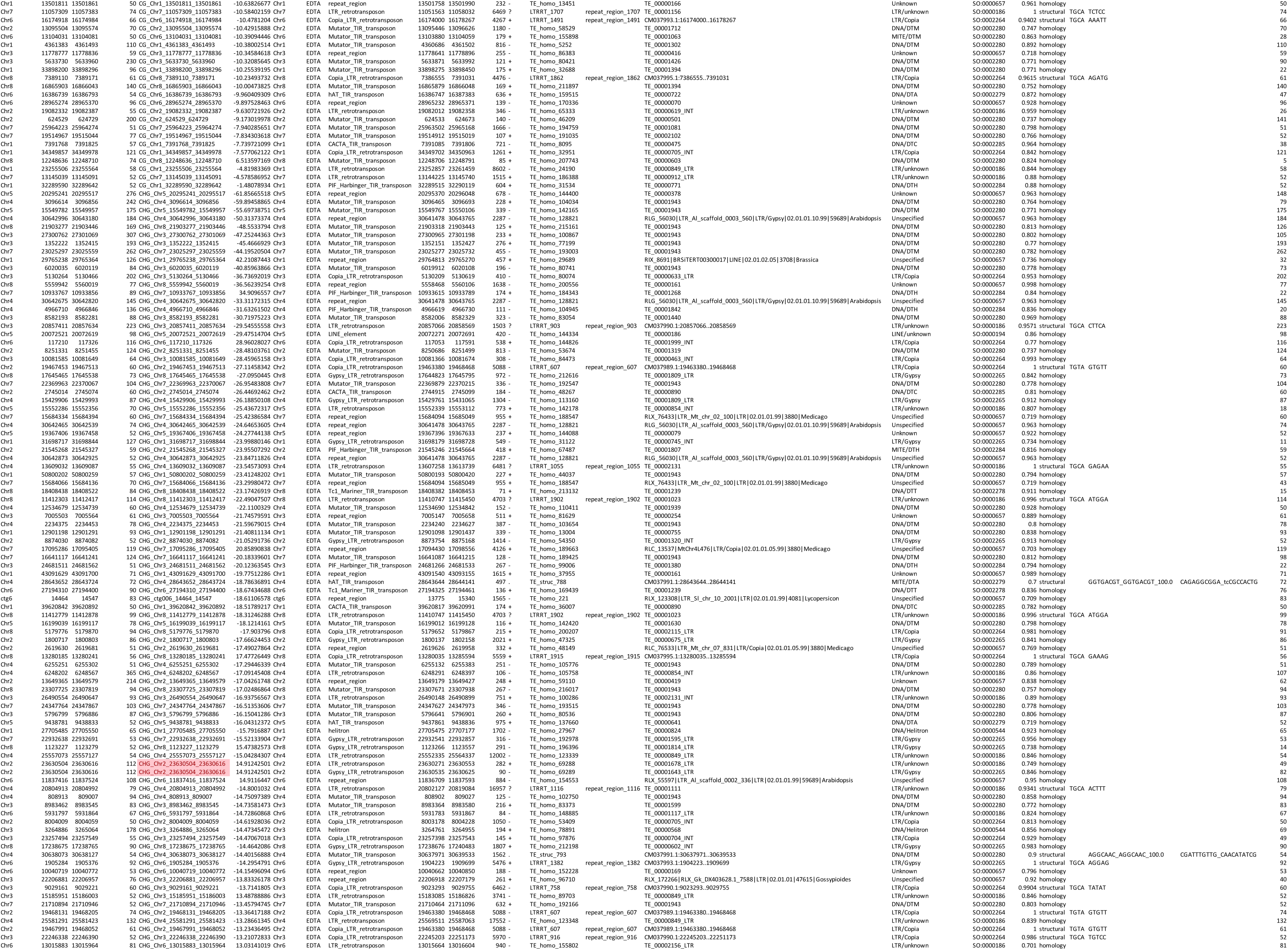
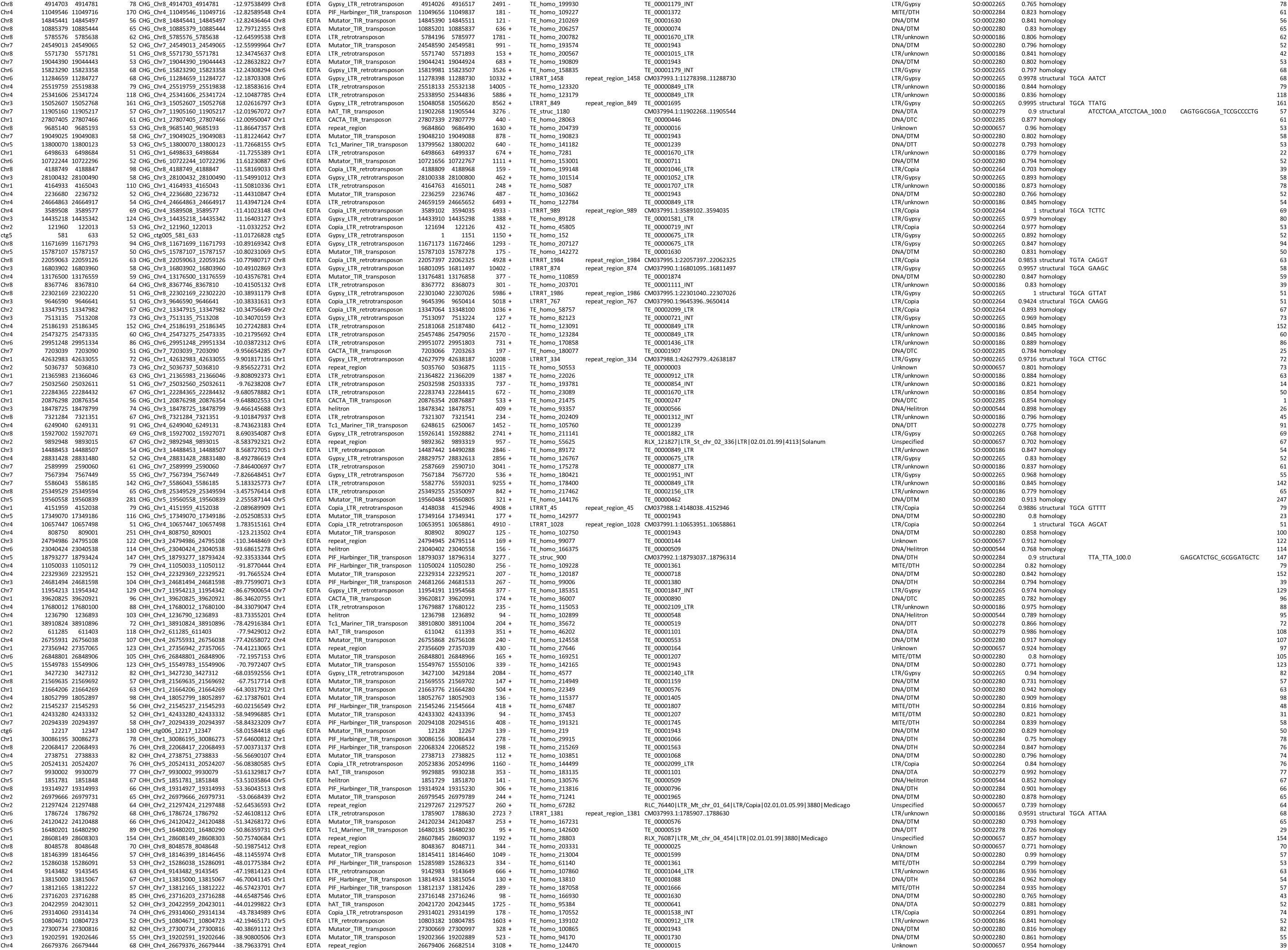
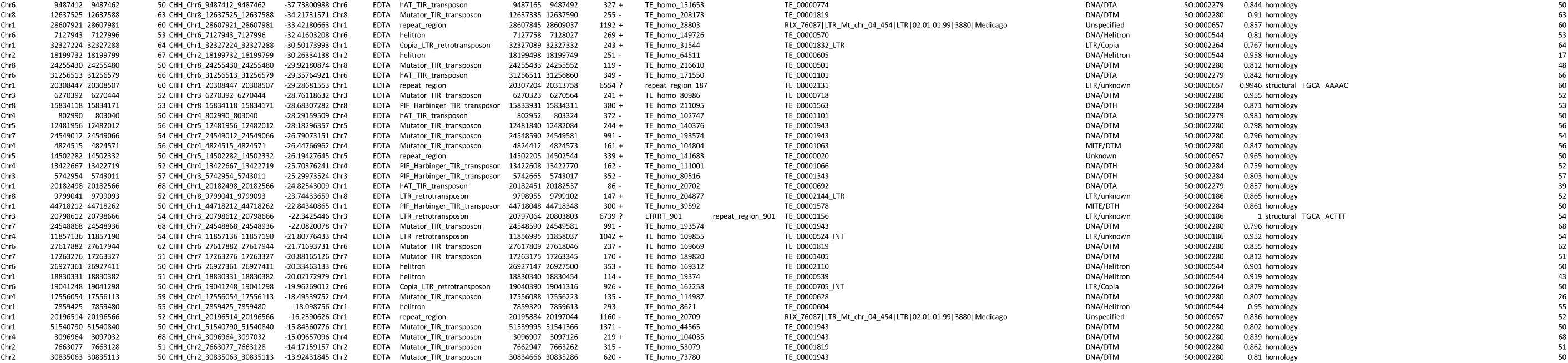

